# Living materials with programmable functionalities grown from engineered microbial co-cultures

**DOI:** 10.1101/2019.12.20.882472

**Authors:** Charlie Gilbert, Tzu-Chieh Tang, Wolfgang Ott, Brandon A. Dorr, William M. Shaw, George L. Sun, Timothy K. Lu, Tom Ellis

## Abstract

Biological systems assemble tissues and structures with advanced properties in ways that cannot be achieved by man-made materials. Living materials self-assemble under mild conditions, are autonomously patterned, can self-repair and sense and respond to their environment. Inspired by this, the field of engineered living materials (ELMs) aims to use genetically-engineered organisms to generate novel materials. Bacterial cellulose (BC) is a biological material with impressive physical properties and low cost of production that is an attractive substrate for ELMs. Inspired by how plants build materials from tissues with specialist cells we here developed a system for making novel BC-based ELMs by addition of engineered yeast programmed to add functional traits to a cellulose matrix. This is achieved via a synthetic ‘symbiotic culture of bacteria and yeast’ (Syn-SCOBY) approach that uses a stable co-culture of *Saccharomyces cerevisiae* with BC-producing *Komagataeibacter rhaeticus* bacetria. Our Syn-SCOBY approach allows inoculation of engineered cells into simple growth media, and under mild conditions materials self-assemble with genetically-programmable functional properties in days. We show that co-cultured yeast can be engineered to secrete enzymes into BC, generating autonomously grown catalytic materials and enabling DNA-encoded modification of BC bulk material properties. We further developed a method for incorporating *S. cerevisiae* within the growing cellulose matrix, creating living materials that can sense chemical and optical inputs. This enabled growth of living sensor materials that can detect and respond to environmental pollutants, as well as living films that grow images based on projected patterns. This novel and robust Syn-SCOBY system empowers the sustainable production of BC-based ELMs.

## Introduction

The nascent field of engineered living materials (ELMs) aims to recapitulate the desirable properties of natural living biomaterials to create novel, useful materials using genetically-engineered organisms^1–4^. Most ELMs to date have either synthesised novel materials from natural macromolecules and polymers harvested from microbial cells^5–14^ or have made use of the multiple functionalities of living cells by embedding these within man-made hydrogels^15–20^. However, the long-term goal of ELMs research is to use engineered cells, rationally reprogrammed at the DNA level, to both make the material and incorporate novel functionalities into it at the same time – thus ‘growing’ functional biomaterials *in situ*^3^.

Most natural examples of living materials achieve their advanced properties by relying on the division of labour afforded by specialised cells performing particular functions. Plant leaves, for example, are self-assembling living materials in which specialised cells are responsible for many complex and useful traits (Fig. 1a). Inspired by the strategy seen in plants, we sought to develop a microbial ELM that utilised a similar approach, dividing bulk material production and functional modification between co-cultured cell types suited for each specialism.

**Fig. 1.**
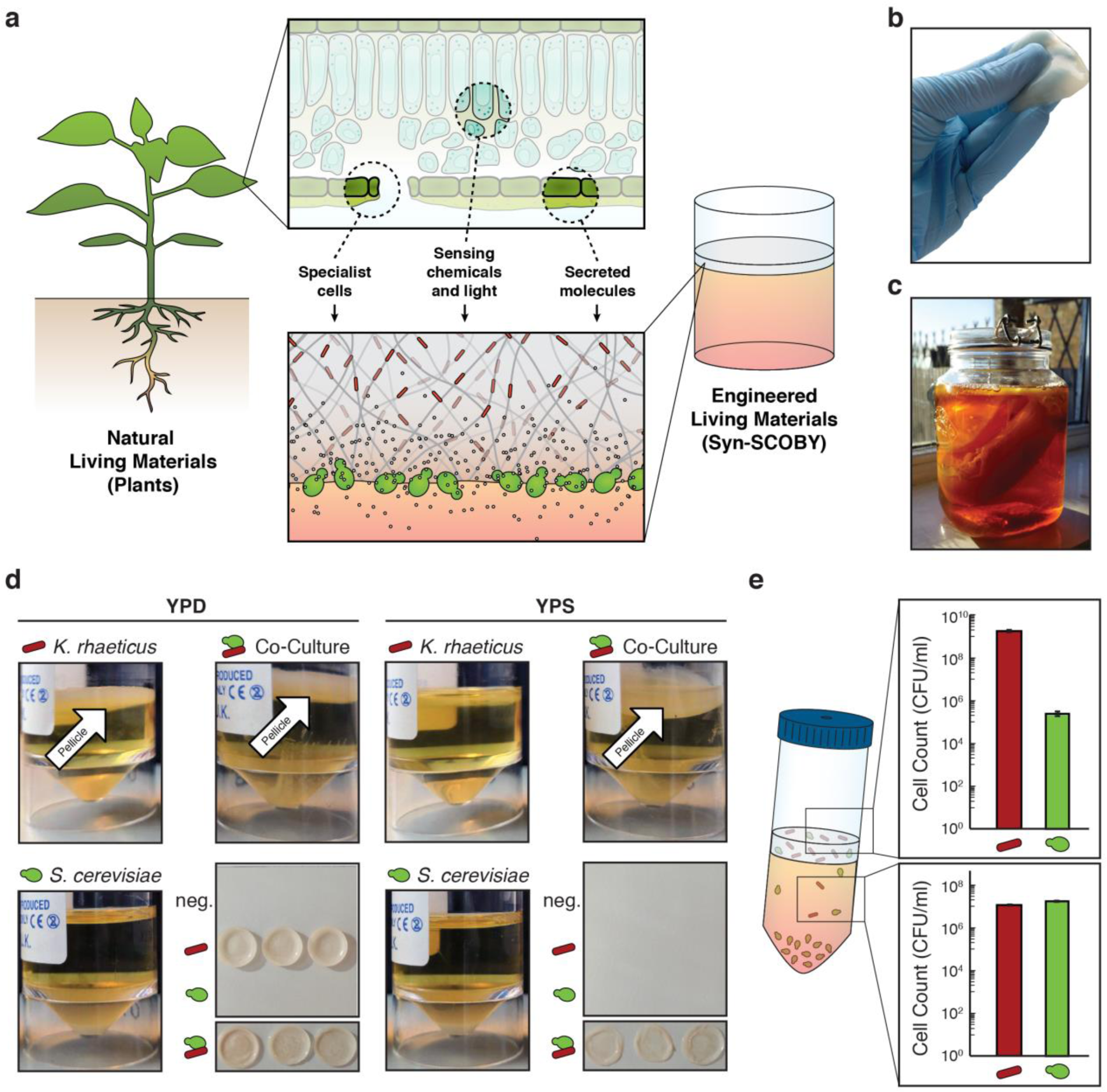
Generating Syn-SCOBY co-cultures with *S. cerevisiae* and *K. rhaeticus*. **(a)** Schematic showing analogies between natural living materials (plants) and engineered living Syn-SCOBY materials, yeast cells depicted in green and bacteria depicted in red. **(b)** An image of a BC pellicle, a flexible but tough material. **(c)** Home-brewed kombucha tea. Both a large submerged mass of BC from previous rounds of fermentation as well as a newly-formed thinner layer at the surface can be seen. **(d)** Images of mono-cultures and co-cultures of *K. rhaeticus* and *S. cerevisiae* grown for 3 days. *S. cerevisiae* grows well in both YPD and YPS media, forming a sediment at the base of the culture. *K. rhaeticus* grew well in YPD medium, forming a thick pellicle layer at the air-water interface (green arrow), but failed to form a pellicle in YPS medium. When co-cultured, in both YPD and YPS, a thick pellicle layer was formed as well as a sediment layer at the base of the culture, indicating both *S. cerevisiae* and *K. rhaeticus* had grown (white arrow). **(e)** Cell counts of a co-culture consisting of *K. rhaeticus* Kr RFP and *S. cerevisiae* yWS167, grown in YPS media. Cell counts were determined by plating and counting the numbers of cells present in the two phases of co-cultures – the liquid layer and the pellicle layer. Samples prepared in triplicate, data represent the mean ±1 SD.

To do this, we turned to a bacterial biofilm material, bacterial cellulose (BC), as our bulk material production system. BC has recently emerged as a promising alternative model ELM system due to high production yields that achieve >10 grams per litre from just a few days growth in low cost, simple sugar media^21^. Various species of Gram-negative acetic acid bacteria – particularly members of the *Komagataeibacter* and *Gluconacetobacter* genera – produce high quantities of extracellular cellulose that yield continuous BC sheets with large surface areas (>1000 cm^2^) when grown in shallow trays containing static liquid^22^. These bacteria secrete cellulose as numerous individual glucan chains bundled into fibrils. Over several days, a thick floating mat forms consisting of a network of BC fibrils around embedded BC-producing bacteria. The resulting biofilm material, referred to as a pellicle (Fig. 1b), is a dense network of ribbonlike cellulose fibrils, each ~50 nm wide and up to 9 µm in length, held together tightly by van der Waals’ forces and hydrogen-bonds^22^.

The ultrapure nature and high crystallinity of BC affords excellent mechanical properties, with individual nanofibers estimated to have tensile strength of at least 2 GPa and Young’s modulus of ~138 GPa^23,24^. It has high porosity, high water retention and a very large surface area. It is both biodegradable and biocompatible and can be made at scale with minimal equipment and low environmental impact and cost. Consequently, BC-based materials have attracted interest for use as surgical and wound dressings, as acoustic diaphragms for headphones and speakers, as battery separators, as additives to cosmetics and as scaffolds for tissue engineering^25^. For example, a BC-based wound-dressing is currently used in the treatment of burns and ulcers^26^.

Genetic modification of BC-producing bacteria has previously been used to alter BC material properties, engineering the bacteria to grow non-native chitin-cellulose^27^ and curdlan-cellulose^28^ co-polymer materials. Modular genetic toolkits for engineering BC-producing bacteria^29–32^ have also been developed but remain minimal in comparison to those for synthetic biology hosts. Crucially, to our knowledge, no tools have ever been reported for recombinant protein secretion from BC-producing bacteria, and their ability to be reprogrammed to sense chosen external cues is also poor. These factors severely limit the engineerability and versatility of BC-based ELMs.

To create an ELM system that leverages specialist engineered cells to expand the functionalities of BC-based ELMs, we took inspiration from kombucha tea to develop a co-culture system that enables autonomous self-assembly and growth of ELMs. Kombucha is a fermented beverage produced by the action of a microbial community commonly referred to as a symbiotic culture of bacteria and yeast (SCOBY) that typically consists of at least one species of BC-producing bacteria and at least one species of yeast (Fig. 1c). Notably, one of the yeast species often found in kombucha fermentations is the model eukaryote *Saccharomyces cerevisiae*^33^.

Based on this observation, we set out here to recreate co-cultures of a BC-producing bacterium, *Komagataeibacter rhaeticus*^29^, with the engineered lab strains of *S. cerevisiae* in order to develop a Synthetic SCOBY (Syn-SCOBY) system. Stable integration of yeast cells among the bacteria during their cellulose biofilm production phase provides us with an engineerable host cell within the growing material that can be rationally programmed at the DNA level for dedicated tasks (Figure 1a). Engineered *S. cerevisiae* therefore act as specialist cells in the system, secreting proteins, sensing chemical and physical signals, and modifying the material properties of surrounding cellulose. In light of the goals of ELMs research, we show that all these functionalities can be achieved in materials that grow and self-assemble in their entirety from only simple growth media in a few days.

## Results and Discussion

### Establishing conditions for stable Syn-SCOBY growth

To introduce yeast as reprogrammable specialist cells within a bacterial culture used for BC material production, we first sought to identify conditions in which *K. rhaeticus* and *S. cerevisiae* are efficiently co-cultured. This required screening for growth at a range of inoculation ratios and in different growth media (Supplementary Fig. 1 and Supplementary Discussion). This initial screen revealed that *K. rhaeticus* grew poorly in sucrose-containing media compared to glucose-containing media, failing to form a pellicle after 3 days (Fig. 1d). However, co-culturing the bacteria with *S. cerevisiae* in the same sucrose media resulted in growth and pellicle formation, revealing a specific growth condition where *K. rhaeticus* growth and BC formation is dependent on the continued presence of the yeast (Supplementary Fig. 2).

**Fig. 2.**
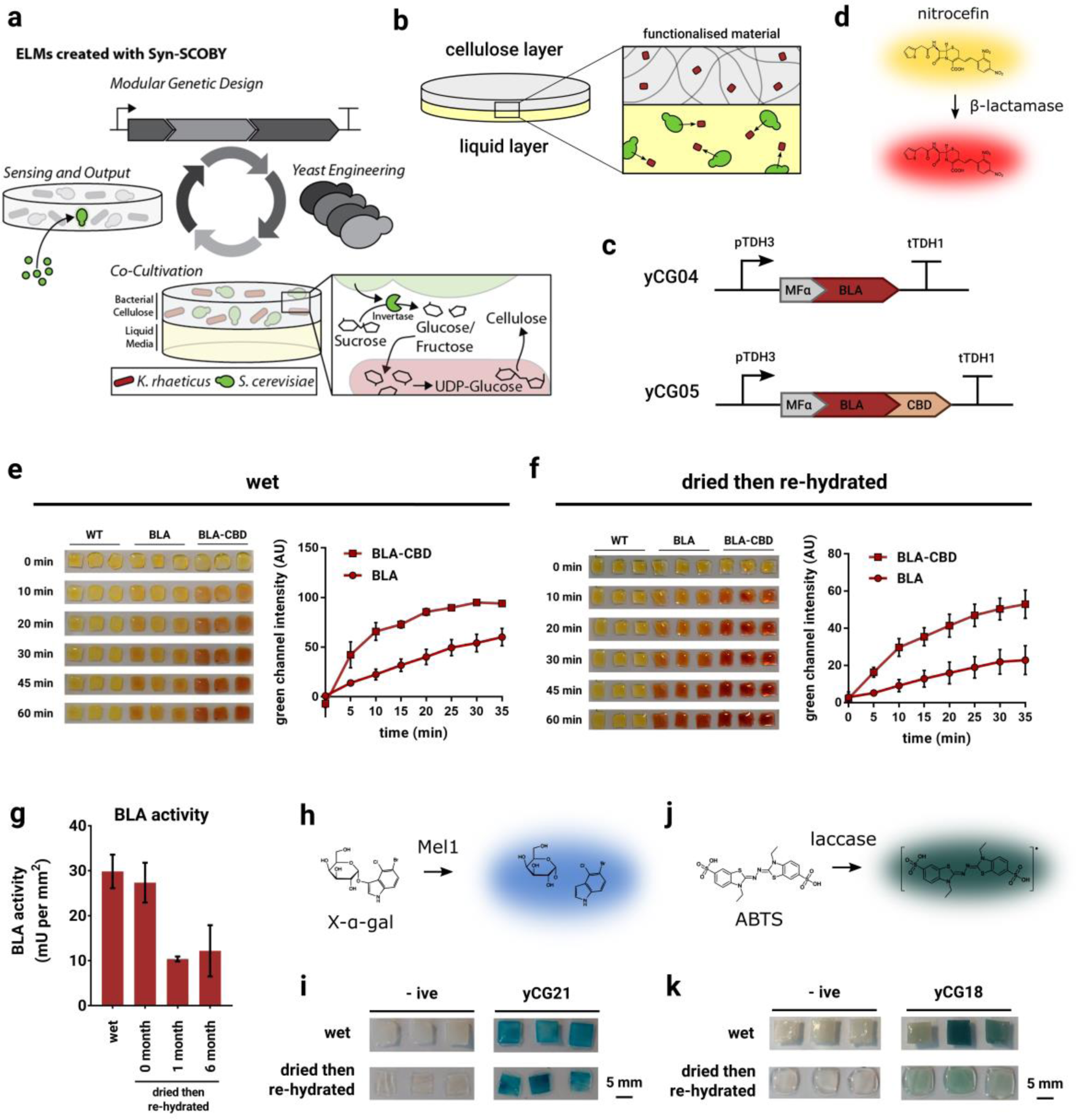
Syn-SCOBYs produce enzyme-functionalised BC materials. **(a)** Schematic of co-culturing BC-producing bacteria with engineerable *S. cerevisiae* yeast to create ELMs, such as modular living biosensors. **(b)** Schematic illustrating concept of functionalisation. *S. cerevisiae* cells, largely found in the liquid layer, secrete a protein that then incorporates into the BC layer, conferring a new functional property. **(c)** BLA-secreting yeast strains yCG04 (BLA) and yCG05 (BLA-CBD). **(d)** Nitrocefin is converted from a yellow substrate to a red product in the presence of β-lactamase enzyme. **(e)** The nitrocefin assay was performed with cut 5×5 mm native wet pellicle samples **(f)** and with dried then re-hydrated pellicle samples from co-cultures with *S. cerevisiae* BY4741 (WT), yCG04 (BLA) and yCG05 (BLA-CBD). Images are shown of pellicles after indicated time points during the assay. In addition, the yellow-to-red colour change was then quantified as green channel intensity using ImageJ software. Samples prepared in triplicate, data represent mean ±1 SD. **(g)** Absolute β-lactamase activities were calculated from native hydrated pellicles (wet) and from pellicles dried then re-hydrated after the indicated time periods. Samples presented here are from pellicles grown in co-culture with yCG05 (BLA-CBD). As pellicle liquid volume is altered by drying, β-lactamase activity is represented by enzyme activity units per unit of pellicle area, to enable cross-comparison. **(h)** X-α-gal is converted from a colourless substrate to a blue product by Mel1 enzyme. **(i)** The X-α-gal assay was performed with wet and dried then re-hydrated pellicle samples from co-cultures with the GFP-secreting strain yCG01 (-ive) or a strain that secretes Mel1 (yCG21), samples prepared in triplicate. **(j)** ABTS is converted from a colourless substrate to a green product by laccase enzyme. **(k)** The ABTS assay was performed with wet and dried then re-hydrated pellicle samples from co-cultures with the GFP-secreting strain yCG01 (-ive) or a strain that secretes CtLcc1 (yCG18), samples prepared in triplicate.

We next characterised a range of co-culture parameters using these standard conditions (see Supplementary Discussion). Pellicle yields, accounting for both cellular and cellulosic biomass both of which are integral components of BC-ELMs, were found to plateau after 3 days of growth (Supplementary Fig. 3). Similar to SCOBYs from brewed kombucha tea, co-cultures remained stable over multiple rounds of passage (Supplementary Figs. 4 and 5). Consistent with a hypothesis in which *S. cerevisiae*-secreted invertase enzymes breakdown sucrose to glucose and fructose and thus facilitate *K. rhaeticus* growth, purified *S. cerevisiae* invertase was shown to improve growth of *K. rhaeticus* when added to sucrose media (Supplementary Fig. 6). Cell counting in the liquid and pellicle layers in co-cultures also revealed that the majority of *S. cerevisiae* cells are found in the liquid below the pellicle, and the majority of *K. rhaeticus* cells are found in the BC pellicle itself (Fig. 1e and Supplementary Fig. 7). Finally, we measured the variability of pellicle yields and cell counts obtained from co-cultures prepared on separate occasions in order to verify the robustness and repeatability of our synthetic SCOBY system (Supplementary Fig. 8).

**Fig. 3.**
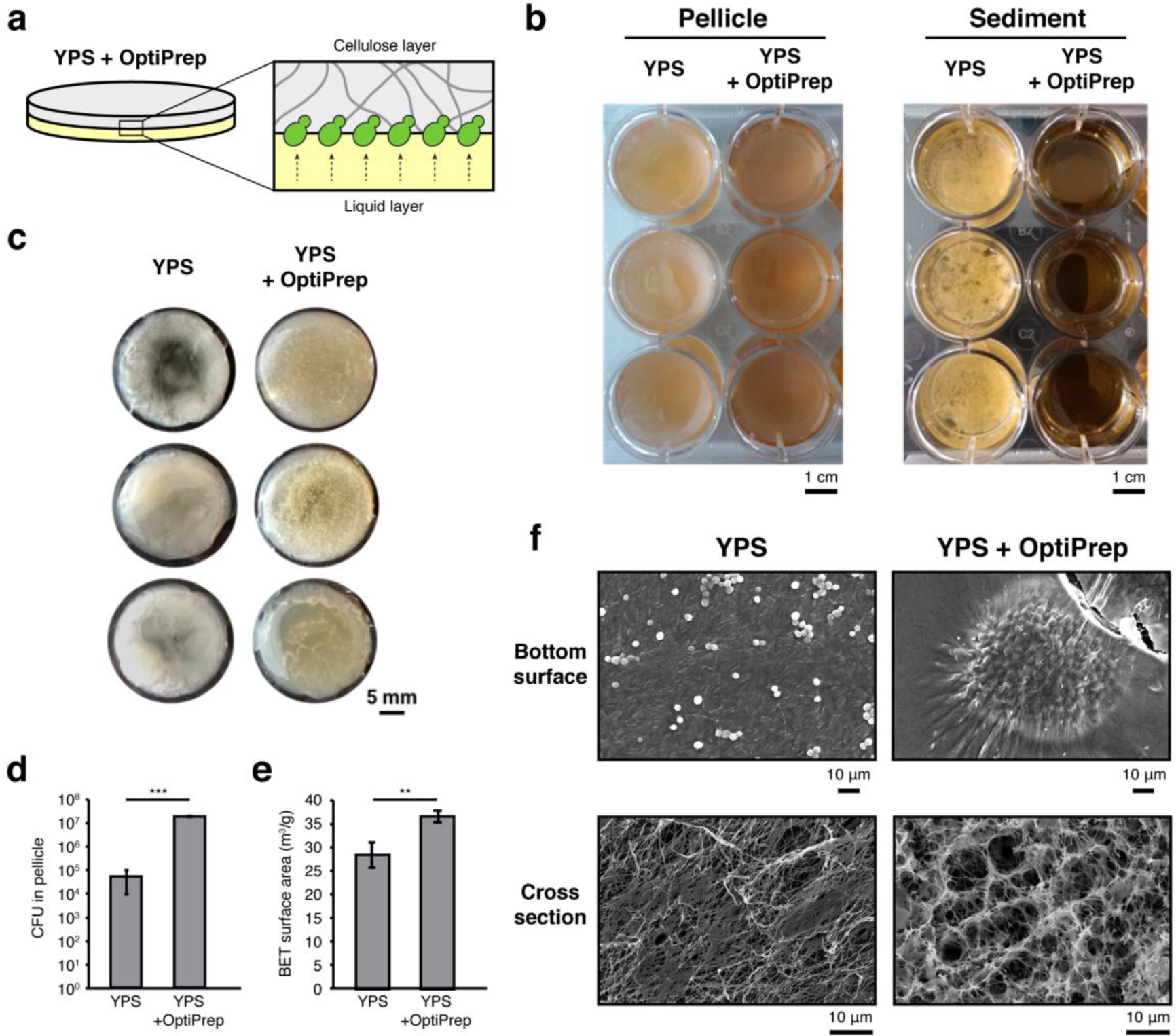
Incorporation of *S. cerevisiae* cells within BC material. **(a)** Schematic demonstrating that modified media density facilitates incorporation of *S. cerevisiae* cells into the pellicle. **(b)** Co-cultures of *K. rhaeticus* Kr RFP and *S. cerevisiae* yWS167 were prepared in YPS media with or without 45% OptiPrep. Images show the pellicles formed at the air-water interface and the liquid below the pellicle, following pellicle removal. **(c)** Isolated pellicles from YPS and YPS+OptiPrep co-cultures of *K. rhaeticus* Kr RPF and *S. cerevisiae* yWS167. Pellicles isolated from co-cultures with OptiPrep have a speckled appearance, due to *S. cerevisiae* colonies. **(d)** Yeast colony forming unit (CFU) counts from pellicles grown in YPS and YPS+OptiPrep using *K. rhaeticus* Kr RPF and *S. cerevisiae* yWS167. Data represent mean and SD. *** p<0.001. **(e)** Brunauer-Emmett-Teller (BET) surface area of the pellicles. Data represent mean and SD. ** p<0.01. **(f)** Sample SEM images of the bottom surface (top) and cross-section (bottom) of pellicles.

**Fig. 4.**
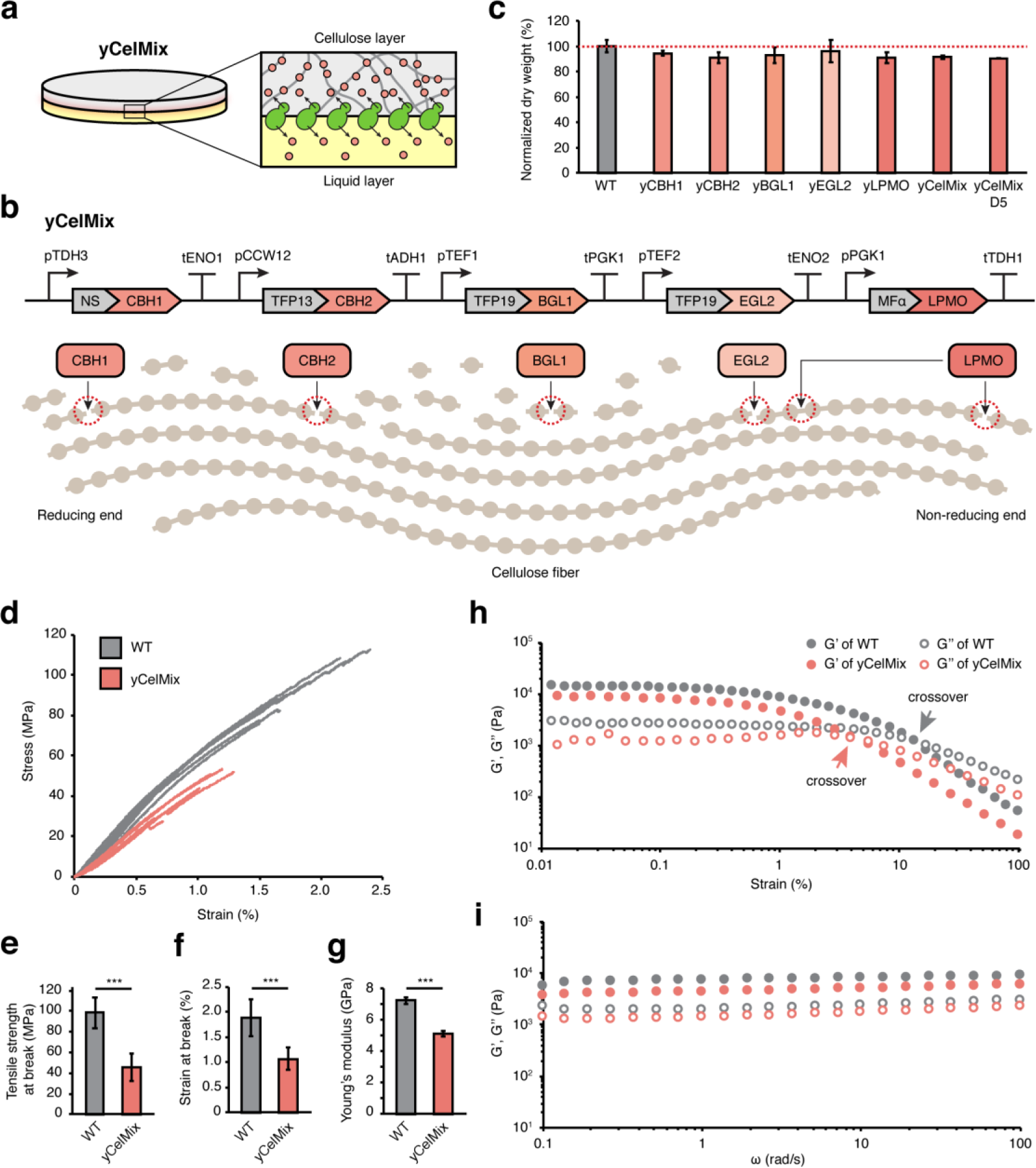
Modifying BC physical material properties. **(a)** Schematic showing yCelMix cells secreting cellulases into the surrounding microenvironment from the bottom surface of the pellicle. **(b)** Schematic illustrating architecture of yCelMix cellulase secretion strain. Expression of each cellulase is controlled by distinct combination of strong promoter and terminator to prevent homologous recombination. CBH1 (cellobiohydrolase from *Chaetomium thermophilum*), CBH2 (cellobiohydrolase from *Chrysosporium lucknowense*), BGL1 (β-glucosidase from *Saccharomycopsis fibuligera*), and EGL2 (endoglucanase from *Trichoderma reesei*) are fused with optimal secretion signals as determined by Lee *et al*^48^ while LPMO (LPMO9H from *Podospora anserine*^47^) was fused with *S. cerevisiae* MFα signal peptide. Signal peptides are colored grey (NS: native signal sequence; TFP13: translational fusion partner 13; TFP19: translational fusion partner 19). Cleavage sites for the cellulases on cellulose polymer are marked as red circles. Each circle represents a monosaccharide unit in the cellulose polymer. **(c)** Normalized pellicle dried weight after 2 day growth of *K. rhaeticus* with different cellulase-secreting *S. cerevisiae*. Red line is weight of wildtype (WT) yeast pellice. yCelMix D5 is after 5 day growth. **(d)** Stress-strain curves of dried WT and yCelMix pellicles. Samples are from 7 WT and 6 yCelMix co-cultures. **(e-g)** Tensile strength at break, strain at break, and Young’s modulus calculated from the data in **(d)**. Data represent the mean and SD. *** p<0.001. **(h, i)** Rheological properties of WT and yCelMix pellicles measured by **(h)** strain sweep at 1 rad/s and **(i)** frequency sweep at 1% strain. Arrows in **(i)** indicate the crossover of the storage modulus (G’, elastic deformation) and loss modulus (G’’, viscous deformation).

**Fig. 5.**
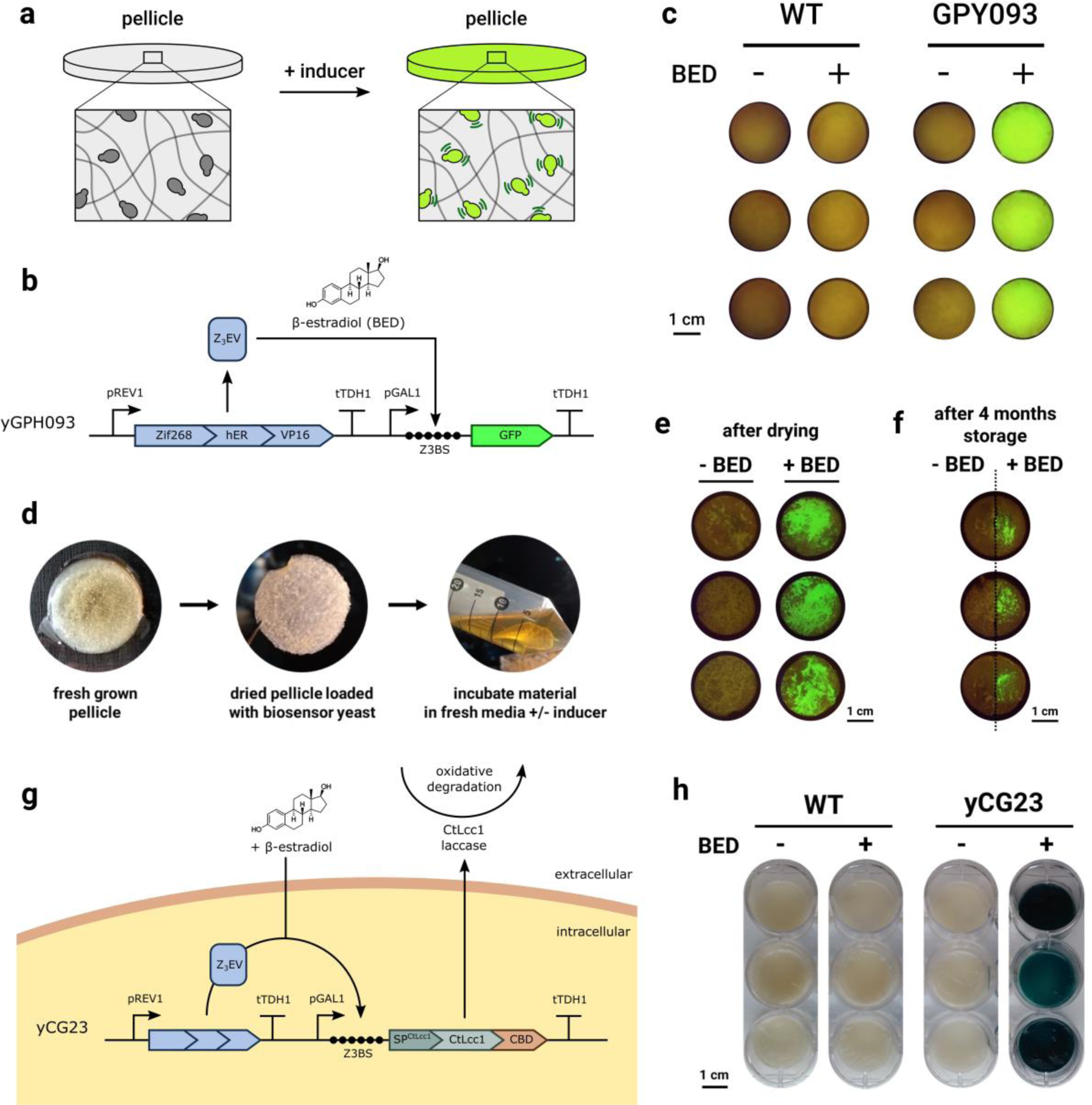
Engineered co-cultures grow living BC sense-and-respond materials. **(a)** Schematic illustrating sense- and-respond pellicle function. Pellicles were grown containing engineered *S. cerevisiae* capable of detecting environmental stimuli and responding by gene expression. **(b)** Schematic showing genetic yGPH093 circuit. The engineered *S. cerevisiae* (yGPH093) senses the presence of the chemical inducer β-estradiol (BED) and in response, produces the reporter protein GFP. The Z_3_ EV synthetic transcription factor is expressed from constitutive promoter pREV1. On addition of β-estradiol Z_3_ EV binds Z_3_ EV binding sites (Z3BSs) in the pGAL1 promoter activating GFP expression. **(c)** Testing biosensor pellicles. Pellicles with either BY4741 (WT) or β-estradiol (BED) responsive (yGPH093) yeast incorporated within the BC matrix were grown with OptiPrep. Triplicate samples were washed, then incubated with agitation in fresh media with or without BED. After 24 h pellicles were washed and imaged for GFP fluorescence by transilluminator. **(d)** Pellicles into which *S. cerevisiae* have been incorporated can be dried, stored, and then revived by incubating in fresh media with or without inducer. Dried pellicles into which yGPH093 was incorporated, were incubated in fresh media in the presence or absence of BED following ambient storage for 1 day **(e)** or 4 months **(f)**. After 24 h, pellicles were imaged for GFP fluorescence by transilluminator. Samples prepared in triplicate. **(g)** Schematic illustrating yCG23 construct design. Similar to yGPH093, yCG23 enables BED-inducible expression and then secretion of CtLcc1 laccase. Extracellular laccases can then be used oxidise a broad range of substrates, enabling degradation of environmental pollutants. **(h)** Native wet pellicles from YPS-OptiPrep co-cultures of the GFP-secreting yCG01 strain (WT) and yCG23 were harvested, washed and inoculated into fresh medium with or without BED. After 24 h growth, pellicles were again harvested and washed and assayed for laccase activity using the colourimetric ABTS assay. Samples prepared in triplicate.

**Fig. 6.**
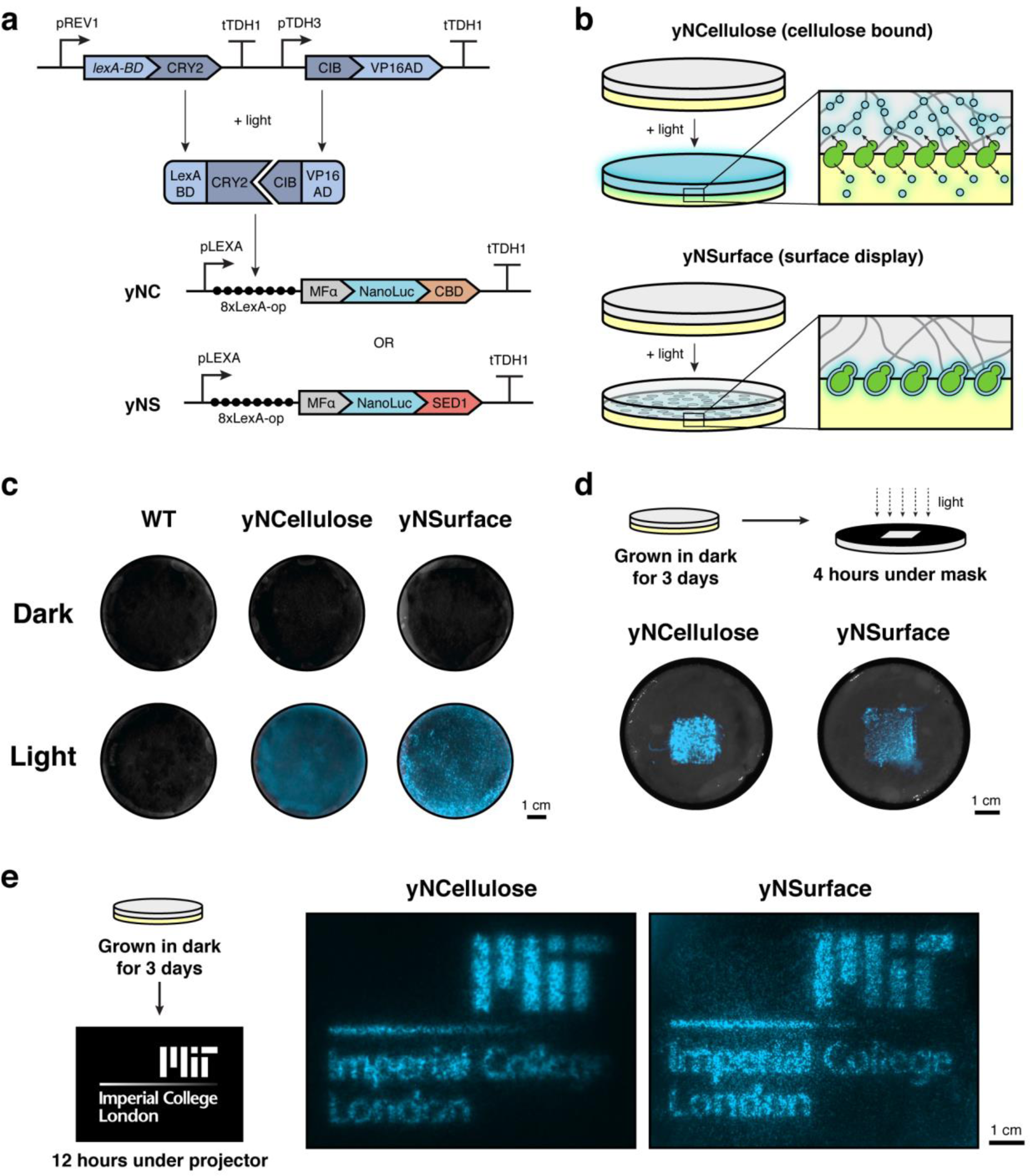
Optical patterning of enzymatically functionalized BC materials. **(a)** Schematic showing the optogenetic circuit. Engineered *S. cerevisiae* strains (yNCellulose and yNSurface) sense blue light and in response express the reporter protein NanoLuc. The LexA-CRY2 and VP16-CIB synthetic transcription factors are expressed from constitutive promoters pREV1 and pTDH3, respectively. Upon exposure to light, the dimer binds LexA-binding-sites (8xLexA-op) in the pLEXA promoter, activating transcription of the NanoLuc gene. **(b)** Schematic illustrating the two modes of functionalisation. The yNCellulose strain secrete NanoLuc-CBD which diffuse into and eventually binds the surrounding cellulose matrix, while the yNSurface strain display NanoLuc-SED1 on the yeast cell surface. **(c)** The pellicles grown in dark or light after 3 days. **(d)** yNCellulose and yNSurface pellicles were grown in dark and then exposed to light under a mask for 4 hours. Pellicles were flipped so that the lower surface, where the majority of yeast cells are localised, was closest to the light source. NanoLuc substrate was applied in the end for visualisation of the pattern created by masking. **(e)** yNCellulose and yNSurface pellicles grown in the dark were exposed to a complicated pattern created by a projector. NanoLuc substrate was applied in the end for visualisation of the pattern created by masking.

### Engineering BC material functionalisation

Having developed and characterised a method for *S. cerevisiae*-*K. rhaeticus* co-culture, we set out to next engineer *S. cerevisiae* to act as specialist cells conferring novel functional properties for BC-based ELMs (Fig. 2a). Unlike BC-producing bacteria, yeasts have a high capacity for recombinant protein secretion, which offers many potential functions. Therefore, the presence of yeast in Syn-SCOBY co-cultures offers a route to BC functionalisation, for example by using *S. cerevisiae* strains engineered to secrete proteins that form part of grown BC materials (Fig. 2b). We assessed this capability using the β-lactam hydrolysing enzyme TEM1 β-lactamase (BLA). Using the yeast toolkit (YTK) system for modular genetic engineering^34^, the BLA catalytic region was cloned downstream of the *S. cerevisiae* mating factor alpha (MFα) pre-pro secretion signal peptide under the control of a strong constitutive promoter (pTDH3) to create yeast strain yCG04 (Fig. 2c). However, as the pellicle makes up only a fraction of the co-culture volume, we hypothesised that addition of a cellulose-binding domain (CBD) to this enzyme might increase the proportion of secreted protein incorporated within the BC layer. Therefore, a second strain (yCG05) was engineered in which a CBD was fused to the C-terminus of the BLA protein (Fig. 2c). The 112 amino acid region from the C-terminus of the Cex exoglucanase from *Cellulomonas fimi* (CBDcex)^35^ was chosen based on previous work demonstrating its ability to bind BC^29^. Culture supernatants from mono-cultures of yCG04 and yCG05 were screened for BLA activity using the colourimetric nitrocefin assay (Fig. 2d), which confirmed secretion from the engineered yeast of active BLA and BLA-CBD, respectively (Supplementary Fig. 9). Next, co-cultures were grown with wild type, BLA-secreting (yCG04) or BLA-CBD-secreting (yCG05) *S. cerevisiae* strains and *K. rhaeticus* bacteria and the resultant BC pellicles were screened for β-lactamase activity. While pellicles grown with wild type yeast showed no BLA activity, clear activity was observed from pellicles from co-cultures with BLA-secreting and BLA-CBD-secreting strains (Fig. 2e), demonstrating direct BC functionalisation. Notably, fusion of the CBD to the enzyme resulted in a 3-fold increase in observed β-lactamase signal from the BC samples, presumably due to the enzyme attaching tightly to the growing material. This increase is seen even though the fusion of the CBD decreases BLA secretion from yeast by around 30% (Supplementary Fig. 9). The value of the CBD-fusion was further demonstrated by assaying the material after washing, where BLA-CBD functionalised pellicles showed a higher activity than those with only BLA (Supplementary Fig. 10).

We then tested whether enzyme-functionalised BC could be dried and re-hydrated while retaining activity. Pellicles produced by co-culturing were dried to create thin, paper-like materials (Supplementary Fig. 11) and later rehydrated and assayed for β-lactamase activity. Re-hydrated pellicles functionalised with both BLA and BLA-CBD demonstrated clear activity after rehydration (Fig. 2f). For absolute quantification of BLA activity, assays were run with pellicles derived from co-cultures with WT yeast spiked with known concentrations of commercial BLA enzyme to create standard curves (Supplementary Fig. 12). This revealed that the drying process itself had little effect on the BLA activity of the material: 29.8 ±3.7 mU/mm^2^ before drying and 27.3 ±4.4 mU/mm^2^ after (Fig. 2g). Identical assays were performed following storage of materials for 1 month or 6 months at room temperature without desiccant. After long-term storage enzymatic activity was retained, although reduced to approximately one-third the original level (Fig. 2g). Overall these experiments show that functionalised materials can be grown and stored at room temperature, retaining their activity for later rehydration and deployment.

To demonstrate the generalizability of our approach, we grew functionalised BC with two other enzymes: alpha-galactosidase enzyme Mel1 (a native *S. cerevisiae* secreted protein) and laccase enzymes from either *Myceliophthora thermophila* (MtLcc1) and *Coriolopsis troggi* (CtLcc1)^36^. All were cloned with a C-terminal CBD and either their native secretion signal peptide or the *S. cerevisiae* MFα signal peptide at the N-terminus (Supplementary Fig. 13a and 14a). Yeast secreting Mel1 fused to the MFα signal peptide and CtLcc1 fused to its native signal peptide exhibited the highest secretion yields in colourimetric plate-based assays (Supplementary Fig. 13b and 14b). These yeast strains were then co-cultured with *K. rhaeticus* and the grown pellicles assayed for enzyme activity. Laccase and α-galactosidase activities were detected in native, wet pellicles (Fig. 2i and 2k) and pellicles functionalised with CtLcc1 or Mel1 retained their activity after drying and re-hydration (Fig. 2i and 2k).

Our Syn-SCOBY approach therefore enables self-assembled production of enzyme-functionalised BC materials grown under mild conditions from simple raw materials. Importantly, these materials retain activity following drying and re-hydration storage under ambient conditions. Our approach could be applied to the production of immobilised enzyme materials used in various industrial processes, such as lipases in the interesterification of food fats and oils^37^, laccases in bioremediation of industrial waste products^38–42^ or β-lactamases in decontamination of antibiotic-contaminated soil and wastewater^43,44^.

Further, we found that an engineered GFP-expressing strain using an identical promoter continually produced functional protein over the course of 16 passages (48 days), indicating our system might allow simple scalable production for many days by passaging (Supplementary Fig. 4). Notably the biological material assembly process is likely to be more sustainable than synthetic methods, as it occurs autonomously, using simple chemical feedstocks under mild conditions and without the need for complex manufacturing steps, such as separate enzyme purification and chemical bonding to the material.

### Incorporating living *S. cerevisiae* within BC materials

The presence of living cells within ELMs during their growth and use enables many of the exciting possibilities for this new class of material. With that in mind, we sought methods to boost the number of yeast cells incorporated into the BC pellicle during its production. *S. cerevisiae* settles to the bottom of static liquid culture as the density of yeast cells is greater than that of water: ~1.11 g/mL compared to 1 g/mL^45^. We hypothesised that increasing the density of the culture medium to >1.11 g/mL would float *S. cerevisiae* cells to the surface, forcing their incorporation into the newly-forming pellicle at the air-water interface (Fig. 3a).

To increase the density of the YPS medium we used OptiPrep, a metabolically-inert aqueous solution of 60% iodixanol with a density of 1.32 g/mL. An initial screen, in which increasing concentrations of OptiPrep were added to YPS medium, revealed that higher density media showed less cell sedimentation (Supplementary Fig. 15). Based on this, co-cultures grown in media with 45% OptiPrep (v/v) were compared to those grown without, and images were taken of resulting materials. Under both conditions, BC pellicles formed, with pellicle thicknesses and yields slightly reduced when grown with OptiPrep (Fig. 3b and Supplementary Figure 16). Interestingly, when pellicles were removed from the cultures, there was a complete absence of sediment in the YPS-OptiPrep medium, in contrast to dense sediment in standard YPS medium (Fig. 3b). Compared to the homogenous surface of pellicles grown in YPS, pellicles grown in YPS-OptiPrep had a speckled appearance due to embedded yeast colonies (Fig. 3c). Addition of OptiPrep gave a ~340-fold increase in estimated *S. cerevisiae* cell count within the pellicle layer, from 5.50 × 10^4^ cfu/mL (±4.58 × 10^4^) to 1.87 × 10^7^ cfu/mL (±1.15 × 10^6^) (Fig. 3d) and thus offers a simple approach to direct incorporation of *S. cerevisiae* cells into the growing material layer under static conditions. Although commercial sources of OptiPrep are relatively expensive, wholesale sources of non-clinical-grade purity iodixanol, used to make OptiPrep, are orders of magnitude less expensive.

In testing, we found that pellicles formed with OptiPrep present exhibited a larger total surface area, as determined by Brunauer-Emmett-Teller (BET) measurement (Fig. 3e). This increase may benefit BC enzyme functionalisation approaches as the internal catalytic surface in cellulose network is expanded. Our observations were confirmed by scanning electron microscopy (SEM) images of dried pellicles and pellicle cross-sections where enlarged macroporous structures are seen when grown in YPS-OptiPrep (Fig. 3f and Supplementary Fig. 17). Furthermore, when grown in YPS, small numbers of *S. cerevisiae* cells were loosely attached to the bottom surface of the pellicle. In contrast, when grown in YPS+OptiPrep, yeast were again localised to the bottom surface of pellicles, but now formed larger foci reminiscent of colonies, containing many cells covered by cellulose fibrils.

To ensure that the drying process did not change the native structure of the pellicles, we also imaged pellicles without lyophilisation by environmental SEM (Supplementary Fig. 18) and observed the same surface morphologies and localisation of yeast at the bottom surface. However, different drying conditions are known to affect the material properties of BC^46^. Fluorescence scans of the pellicle containing mScarlet-expressing yeast showed punctate, colony-like lateral distribution (Supplementary Fig. 19) and even after washing of these pellicles more than half of the yeast were still present, demonstrating their incorporation within the BC material (Supplementary Fig. 20).

### Modified BC physical properties via programmed enzyme secretion

To change the bulk physical properties of the BC during its growth, we next explored modifying the cellulosic matrix by having cellulases secreted from engineered *S. cerevisiae* cells during co-culturing. We constructed a *S. cerevisiae* strain, yCelMix, in which cellobiohydrolases (CBH1 and CBH2), endoglucanase (EGL2), β-glucosidase (BGL1), and lytic polysaccharide monooxygenases (LPMO) are optimized to be secreted simultaneously for synergistic cellulose degradation, as described in previous work^47,48^ (Fig. 4a-b and Supplementary Fig. 21). Co-cultures of *K. rhaeticus* and yCelMix yeast still formed pellicles after two days, albeit with slightly decreased dried weight (Fig. 4c), indicating that rates of cellulase secretion and action were slower than the rate of bacterial cellulose biosynthesis. Extending the incubation time to five days did not further reduce the cellulose mass, potentially due to the limited diffusion of enzymes, low pH, local accumulation of reaction products, or cellulase deactivation at the air-liquid interface in static culture^49^. Interestingly, the total surface area of yCelMix pellicles was smaller than that of pellicles grown with normal wildtype yeast (Supplementary Fig. 22). This may result from a decrease in structure due to cellulose degradation. Indeed, SEM imaging of yCelMix reveals that a loose fibrous network replaces the densely-packed cellulosic matrix at both the top and bottom surfaces (Supplementary Fig. 23).

Although cellulose yield was not greatly reduced by cellulase secretion, the mechanical properties of the pellicles were altered. Stress-strain curves from tensile tests demonstrated a clear difference between WT pellicle and yCelMix pellicle (Fig. 4d). Specifically, while both were brittle (as determined by fracture strength), the yCelMix pellicle could only sustain 45.7 MPa while the WT pellicle can bear 98.3 MPa of stress before fracturing (Fig. 4e). The yCelMix pellicle could also only be stretched to half of the length that the WT pellicle could be elongated to before breaking (Fig. 4f). Furthermore, secreted cellulases reduced the stiffness of the cellulose matrix, lowering the Young’s modulus from 7.2 GPa (WT) to 5.1 GPa (yCelMix) (Fig. 4g and Supplementary Fig. 24). Although the underlying mechanisms by which these enzymes modify the material properties of the BC matrix require further investigation, our results are consistent with other reports that the continuity of the cellulose fibrils and integrity of network structure are essential for the strength and stiffness of bacterial cellulose materials^50,51^.

The weakening of microstructure in the yCelMix pellicle was also reflected by its rheological properties. In the strain sweep experiment (Fig. 4h), both pellicles showed gel-like behavior at low strain, as G’ always dominates over G’’. The G’ and G’’ of yCelMix pellicle were both lower than those of WT pellicles, again demonstrating that cellulases reduced the stiffness of the BC material (Fig. 4h) in a frequency-independent fashion (Fig. 4i). As the applied strain increased beyond the G’ G’’ crossover point, where microcracks accumulated and major rupture appeared, the pellicles switched to a viscoelastic liquid and started to flow. Notably, yCelMix pellicle had an earlier onset of crossover, indicating a faster breakdown of network structure in the matrix. In addition, the more pronounced G’’ maximum in the yCelMix pellicle (Fig. 4i) demonstrates that deformation energy was converted into friction heat from the free broken fibrils near the microcracks, which originated from a disintegrated and weaker cellulose network. Our results suggest cellulases secreted from the yeast can effectively weaken the mechanical and viscoelastic properties of BC materials, complementing recent studies showing carbohydrate binding domain (CBD) protein additives can enhance the strength of cellulosic materials^52,53^.

### Engineering living BC materials to sense-and-respond

*S. cerevisiae* strains are routinely engineered in synthetic biology as sensors that respond to chemical and physical stimuli and trigger changes in gene expression. Our modular co-culture Syn-SCOBY approach enables easy incorporation of such biosensor yeast into grown BC materials (Fig. 5a). To demonstrate this, we selected a chemically-inducible system where the oestrogen steroid hormone β-estradiol (BED) triggers activation of transcription from a target promoter^54,55^. Specifically, we used an *S. cerevisiae* strain (yGPH093) expressing the BED-activated synthetic transcription factor Z_3_ EV and a GFP reporter under control of the Z_3_ EV-responsive promoter (Fig. 5b)^56^. Pellicles grown from *K. rhaeticus* with yGPH093 yeastproduced a strong GFP signal when exposed to exogenous BED (Fig. 5c), demonstrating co-cultures that self-assembly living BC materials able sense-and-respond to environmental stimuli.

To test viability of these BC-yeast living materials after drying and long-term storage (Fig. 5d), pellicles containing a GFP-expressing yeast strain (yWS167) were grown from co-culture, dried and stored at room temperature under ambient conditions. They were then degraded enzymatically and cell counts of *S. cerevisiae* determined by plating. Although drying resulted in a substantial loss of viable yeast within the BC, viable cells could be recovered even after 1 month of storage under ambient conditions (Supplementary Fig. 25).

To demonstrate that these small numbers of viable cells were still sufficient for biosensor materials to remain functional, pellicles grown with yGPH093 were dried, stored and then revived by being incubated in fresh medium for 24 hours in the presence or absence of chemical inducer. Pellicles containing yGPH093 once again yielded a clear GFP signal in the presence of BED (Fig. 5e). Further, re-hydrated dried pellicles yielded a detectable GFP signal in the presence of BED even after ambient storage for 4 months (Fig. 5f). While these sense-and-response functions require the re-addition of growth media, diverse sample types could be used by supplementing with concentrated nutrient stocks. A similar non-ELMs approach has previously enabled *S. cerevisiae* biosensor strains to function in blood, urine and soil^57^.

As a demonstration of further biosensor capabilities, we grew and verified BC sensors where the yeast sense a protein via a G protein coupled receptor (GPCR). Specifically, we used biosensor strain yWS890 to detect the *S. cerevisiae* MFα peptide and produce GFP in response^58^ (Supplementary Fig. 26). GPCRs are the major class of membrane protein receptors across eukaryotes and detect a remarkable range of different chemical and physical stimuli. Our modular approach opens up the opportunity to directly use previously-developed biosensing strains^57,59,60^ and grow them into functionalised BC materials.

With living cells in materials, it is possible to not only sense the environment, but to also react to sensed cues. Future ELMs will be able to dynamically remodel and adapt their properties in response to environmental changes. To demonstrate a step towards this, we linked our sensing systems in yeast to the production of a functional response. Since laccase enzymes have been previously shown to degrade BED^61^, we engineered yeast strain yCG23 to secrete laccase CtLcc1 from the BED-inducible promoter (Fig. 5g and Supplementary Fig. 27). The *K. rhaeticus* and yCG23 co-culture grew a BC-based living material that could simultaneously detect the presence of BED and in response secrete active laccase enzyme (Fig. 5h). Notably, BED is an important environmental pollutant, with potential effects on exposed aquatic species and humans^62,63^. Our approach shows that co-cultures can grow functional living materials engineered to sense changes in their environment and respond accordingly.

### Spatial patterning of catalytic living materials by optogenetics

As an alternative to sensing chemical inputs, we also investigated the use of optogenetics to develop light-responsive BC-based ELMs. For this, we implemented and optimized a blue light sensing system in *S. cerevisiae* based on the CRY2/CIB transcription system^64^ (Supplementary Fig. 28). DNA programs added to the yeast used this light-inducible promoter to trigger expression of NanoLuc, a luciferase reporter enzyme. Two versions were designed, a yNSurface strain that displays NanoLuc on the yeast cell surface, and yNCellulose which secretes a NanoLuc-CBD version into the culture to bind to the BC (Fig. 6a and b).

To demonstrate that Syn-SCOBY pellicles can be engineered to respond an optical input, *K. rhaeticus* was co-cultured with either yeast strain, with or without exposure to white light. After 3 days incubation in light, BC pellicles grown from both co-cultures formed material layers that exhibited high bioluminescence when substrate was applied. Equivalent pellicles grown in the dark for 3 days showed nearly zero luciferase activity (Fig. 6c). Notably, the yNCellulose pellicle exhibited evenly-distributed NanoLuc activity across the material surface, whereas the yNSurface pellicle had localized foci, corresponding to yeast distribution. This demonstrates that the local distribution of protein functionalisation in BC materials can be altered by the design of the DNA construct placed into the yeast cells.

We further explored the responsiveness of these pellicles to light patterns created by masking and projecting. With Syn-SCOBYs containing either yNSurface or yNCellulose yeast, we grew “living films” in the dark for 3 days before further growth with light exposure (Fig. 6d). With light patterning done by masking, both BC pellicles exhibited bright foci within their patterns, likely because there was insufficient time for the NanoLuc to diffuse through the cellulose matrix away from the yNCellulose cells in this experiment. When the pattern was instead projected, both grown pellicles nicely reproduced the desired patterns in their final luciferase activity output (Fig. 6e). In this case, yNSurface produced a sharper pattern compared with yNCellulose, which is expected due to the ability of NanoLuc to diffuse in the former case.

Notably, areas more intricately patterned showed poorer resolution, possibly due to the internal light scattering from the opaque cellulose matrix. In addition, changing the *S. cerevisiae* density in the co-culture by either slowing down yeast growth rate or increasing the incubation time in the dark was able to decrease (Supplementary Fig. 29b), or increase (Supplementary Fig. 29c) the resolution of the patterns, respectively. These knobs, along with different patterning modes (light mask for rapid, large scale prototyping and projection for complicated patterning), provide a basic toolset for optogenetic control of the functionalization of BC-based ELMs. We anticipate that this system could be readily expandable to multi-color optogenetics by further incorporating alternative light-based dimerizing systems^64^ and by linking these to other enzymatic outputs, enabling spatially segregated enzymatic cascade in BC materials.

## Conclusions

Bacterial cellulose is a promising biological material for the development of ELMs. Inspired by plants, we here developed a novel Syn-SCOBY approach to introduce specialised cells into the material production culture, and in doing so improved the engineerability and functionalisation of BC. We developed standard conditions to enable reproducible co-culturing leading to self-assembly of BC-based ELMs. Our Syn-SCOBY system allowed us to leverage the large body of well-described *S. cerevisiae* genetic parts and synthetic biology tools to create a range of BC-based ELMs. Notably, Syn-SCOBY is not only valuable for the development of biological ELMs, but also offers a potential model system for synthetic ecology, in which microbe-microbe interactions could be investigated and engineered.

As our approach is highly modular, we can incorporate existing *S. cerevisiae* strains and synthetic biology tools to genetically program functional properties into grown BC materials. Enzymes secreted from *S. cerevisiae* are incorporated into BC and functionalise the material with the desired catalytic activities as the material forms. This avoids the need to introduce the high-cost, low-sustainability steps of separate protein purification and enzyme-material chemical bonding into the manufacturing process, achieving in one-step what would normally require separate and expensive manufacturing processes. The functionalised BC materials we generated here could be applied to the degradation of β-lactam antibiotics or oestrogen hormones present in wastewater streams, both of which are environmental pollutants. Notably, these applications may be facilitated by the fact that functionalised BC materials could be dried and later rehydrated, retaining catalytic activity. Further, BC materials are biocompatible, produced by growth under mild conditions with simple culture media and made in high yield at little cost from microbes already commonly-used in both the food and healthcare industries.

The approach is also highly adaptable; numerous other protein targets – be they enzymes, binding or adhesion proteins, or even structural proteins – could be secreted from *S. cerevisiae* to add various desired biological properties to the material. However, the feasibility of these approaches will depend on the secretion yields for given proteins – which may be lower under static growth conditions compared to shake-flask or bioreactor growth – as well as the stability of the proteins outside the cell or after any required sterilisation procedures.

Where living cells are permitted in applications, *S. cerevisiae* biosensor strains offer a fast way to incorporate a sensing function into a material. The yeast can modify bulk mechanical properties of BC as well as create living materials able to respond to environmental changes. Sense-and-respond ELMs could find application in a variety of contexts, in biosensing, bioremediation or creation of patterned materials. Since our approach is highly-adaptable, numerous other *S. cerevisiae* biosensor strains able to detect pathogens^57^, environmental pollutants^59^ or biomarkers^65^ could be used in conjunction with our co-culture method.

Organisms are remarkable material-producing systems, capable of converting simple feedstocks into complex materials with diverse chemical and physical properties, while simultaneously controlling their morphology over multiple length scales, and remodelling material properties in response to environmental cues. Synthetic material systems capable of recreating all of these behaviours do not exist. As such, the ability to genetically control the process of material self-assembly with the same level of sophistication seen in natural biological materials could revolutionise the manufacture of products for use in numerous arenas of human life and society. This work represents a substantial step towards the ability to genetically program biological material assembly. In summary, our Syn-SCOBY approach showcases the viability of microbial co-cultures combined with synthetic biology tools to design, grow and test functional BC-based materials.

## Acknowledgements

The authors thank Dr. Georgios Pothoulakis, Dr. Carlos Bricio-Garberi, Prof. Benjamin E. Wolfe, and Elizabeth Landis for advice and discussions, Jelle van der Hilst for contributions to co-culture methods, and Bolin An for assisting with photo taking. Work at Imperial College London was funded by UK Engineering and Physical Sciences Research Council (EPSRC) awards EP/M002306/1 and EP/N026489/1 and an Imperial College London President’s Scholarship to CG. WO was supported by a research fellowship (OT 577/1-1) from the German Research Foundation (DFG). Work at MIT was funded by Army Research Office award W911NF-11-1-0281 and Institute for Soldier Nanotechnologies award W911NF-13-D-0001, T.O. 4. TCT was supported by the MIT J-WAFS Fellowship. Work across both institutions was funded by the MIT-MISTI MIT-Imperial College London Seed Fund.

## References

(1) Chen, A. Y., Zhong, C., and Lu, T. K. (2015) Engineering living functional materials. ACS Synth. Biol. 4, 8–11.

(2) Nguyen, P. Q. (2017) Synthetic biology engineering of biofilms as nanomaterials factories. Biochem. Soc. Trans. 45, 585–597.

(3) Nguyen, P. Q., Courchesne, N. D., Duraj-thatte, A., Praveschotinunt, P., and Joshi, N. S. (2018) Engineered Living Materials: Prospects and Challenges for Using Biological Systems to Direct the Assembly of Smart Materials. Adv. Mater. 1704847, 1–34.

(4) Gilbert, C., and Ellis, T. (2019) Biological Engineered Living Materials: Growing Functional Materials with Genetically Programmable Properties. ACS Synth. Biol. 8, 1–15.

(5) Blanco, L. P., Evans, M. L., Smith, D. R., Badtke, M. P., and Chapman, M. R. (2012) Diversity, biogenesis and function of microbial amyloids. Trends Microbiol. 20, 66–73.

(6) Kalyoncu, E., Ahan, R. E., Olmez, T. T., and Safak Seker, U. O. (2017) Genetically encoded conductive protein nanofibers secreted by engineered cells. RSC Adv. 7, 32543–32551.

(7) Seker, U. O. S., Chen, A. Y., Citorik, R. J., and Lu, T. K. (2017) Synthetic Biogenesis of Bacterial Amyloid Nanomaterials with Tunable Inorganic-Organic Interfaces and Electrical Conductivity. ACS Synth. Biol. 6, 266–275.

(8) Dorval Courchesne, N.-M., DeBenedictis, E. P., Tresback, J., Kim, J. J., Duraj-Thatte, A., Zanuy, D., Keten, S., and Joshi, N. S. (2018) Biomimetic engineering of conductive curli protein film. Nanotechnology 29, 509501.

(9) Chen, A. Y., Deng, Z., Billings, A. N., Seker, U. O. S., Lu, M. Y., Citorik, R. J., Zakeri, B., and Lu, T. K. (2014) Synthesis and patterning of tunable multiscale materials with engineered cells. Nat. Mater. 13, 515–23.

(10) Moser, F., Tham, E., González, L. M., Lu, T. K., and Voigt, C. A. (2019) Light-Controlled, High-Resolution Patterning of Living Engineered Bacteria Onto Textiles, Ceramics, and Plastic. Adv. Funct. Mater. 1901788.

(11) Nguyen, P. Q., Botyanszki, Z., Tay, P. K. R., and Joshi, N. S. (2014) Programmable biofilm-based materials from engineered curli nanofibres. Nat. Commun. 5, 4945.

(12) Nussbaumer, M. G., Nguyen, P. Q., Tay, P. K. R., Naydich, A., Hysi, E., Botyanszki, Z., and Joshi, N. S. (2017) Bootstrapped Biocatalysis: Biofilm-Derived Materials as Reversibly Functionalizable Multienzyme Surfaces. ChemCatChem 9, 4328–4333.

(13) Duraj-Thatte, A. M., Courchesne, N. D., Praveschotinunt, P., Rutledge, J., Lee, Y., Karp, J. M., and Joshi, N. S. (2019) Genetically Programmable Self-Regenerating Bacterial Hydrogels. Adv. Mater. 1901826.

(14) Dorval Courchesne, N.-M., Duraj-Thatte, A., Tay, P. K. R., Nguyen, P. Q., and Joshi, N. S. (2016) Scalable Production of Genetically Engineered Nanofibrous Macroscopic Materials via Filtration. ACS Biomater. Sci. Eng. acsbiomaterials.6b00437.

(15) Park, S. J., Gazzola, M., Park, K. S., Park, S., Di Santo, V., Blevins, E. L., Lind, J. U., Campbell, P. H., Dauth, S., Capulli, A. K., Pasqualini, F. S., Ahn, S., Cho, A., Yuan, H., Maoz, B. M., Vijaykumar, R., Choi, J. W., Deisseroth, K., Lauder, G. V., Mahadevan, L., and Parker, K. K. (2016) Phototactic guidance of a tissue-engineered soft-robotic ray. Science (80-.). 353, 158–162.

(16) Van Tittelboom, K., De Belie, N., De Muynck, W., and Verstraete, W. (2010) Use of bacteria to repair cracks in concrete. Cem. Concr. Res. 40, 157–166.

(17) Wang, J., Van Tittelboom, K., De Belie, N., and Verstraete, W. (2012) Use of silica gel or polyurethane immobilized bacteria for self-healing concrete. Constr. Build. Mater. 26, 532–540.

(18) Gerber, L. C., Koehler, F. M., Grass, R. N., and Stark, W. J. (2012) Incorporating microorganisms into polymer layers provides bioinspired functional living materials. Proc. Natl. Acad. Sci. U. S. A. 109, 90–94.

(19) Gerber, L. C., Koehler, F. M., Grass, R. N., and Stark, W. J. (2012) Incorporation of Penicillin-Producing Fungi into Living Materials to Provide Chemically Active and Antibiotic-Releasing Surfaces. Angew. Chemie 124, 11455–11458.

(20) Liu, X., Tang, T.-C., Tham, E., Yuk, H., Lin, S., Lu, T. K., and Zhao, X. (2017) Stretchable living materials and devices with hydrogel–elastomer hybrids hosting programmed cells. Proc. Natl. Acad. Sci. 114, 2200–2205.

(21) Chawla, P. R., Bajaj, I. B., Survase, S. A., and Singhal, R. S. (2009) Microbial cellulose: Fermentative production and applications. Food Technol. Biotechnol.

(22) Huang, Y., Zhu, C., Yang, J., Nie, Y., Chen, C., and Sun, D. (2014) Recent advances in bacterial cellulose. Cellulose 21, 1–30.

(23) Hsieh, Y. C., Yano, H., Nogi, M., and Eichhorn, S. J. (2008) An estimation of the Young’s modulus of bacterial cellulose filaments. Cellulose 15, 507–513.

(24) Kondo, T., and Rytczak, P. (2016) Bacterial NanoCellulose Characterization. Bact. Nanocellulose 59–71.

(25) Wang, J., Tavakoli, J., and Tang, Y. (2019) Bacterial cellulose production, properties and applications with different culture methods – review. Carbohydr. Polym. 219, 63–76.

(26) Ludwicka, K., Jedrzejczak-Krzepkowska, M., Kubiak, K., Kolodziejczyk, M., and Pankiewicz, T. (2016) Medical and Cosmetic Applications of Bacterial NanoCellulose. Bact. Nanocellulose 145–165.

(27) Yadav, V., Paniliatis, B. J., Shi, H., Lee, K., Cebe, P., and Kaplan, D. L. (2010) Novel in vivo- degradable cellulose-chitin copolymer from metabolically engineered gluconacetobacter xylinus. Appl. Environ. Microbiol. 76, 6257–6265.

(28) Fang, J., Kawano, S., Tajima, K., and Kondo, T. (2015) In Vivo Curdlan/Cellulose Bionanocomposite Synthesis by Genetically Modified Gluconacetobacter xylinus. Biomacromolecules 16, 3154–3160.

(29) Florea, M., Hagemann, H., Santosa, G., Abbott, J., Micklem, C. N., Spencer-Milnes, X., de Arroyo Garcia, L., Paschou, D., Lazenbatt, C., Kong, D., Chughtai, H., Jensen, K., Freemont, P. S., Kitney, R., Reeve, B., and Ellis, T. (2016) Engineering control of bacterial cellulose production using a genetic toolkit and a new cellulose-producing strain. Proc. Natl. Acad. Sci. 201522985.

(30) Teh, M. Y., Ooi, K. H., Danny Teo, S. X., Bin Mansoor, M. E., Shaun Lim, W. Z., and Tan, M. H. (2019) An Expanded Synthetic Biology Toolkit for Gene Expression Control in Acetobacteraceae. ACS Synth. Biol. 8, 708–723.

(31) Jacek, P., Ryngajłło, M., and Bielecki, S. (2019) Structural changes of bacterial nanocellulose pellicles induced by genetic modification of Komagataeibacter hansenii ATCC 23769. Appl. Microbiol. Biotechnol. 1–15.

(32) Walker, K. T., Goosens, V. J., Das, A., Graham, A. E., and Ellis, T. (2018) Engineered cell-to-cell signalling within growing bacterial cellulose pellicles. Microb. Biotechnol. 0, 1–9.

(33) Jayabalan, R., Malini, K., Sathishkumar, M., Swaminathan, K., and Yun, S. E. (2010) Biochemical characteristics of tea fungus produced during kombucha fermentation. Food Sci. Biotechnol. 19, 843–847.

(34) Lee, M. E., DeLoache, W. C., Cervantes, B., and Dueber, J. E. (2015) A Highly-characterized Yeast Toolkit for Modular, Multi-part Assembly. ACS Synth. Biol. 4, 975–986.

(35) Ong, E., Gilkes, N. R., Miller, R. C., Warren, R. a, and Kilburn, D. G. (1993) The cellulose-binding domain (CBD(Cex)) of an exoglucanase from *Cellulomonas fimi*: production in *Escherichia coli* and characterization of the polypeptide. Biotechnol. Bioeng. 42, 401–9.

(36) Antošová, Z., Herkommerová, K., Pichová, I., and Sychrová, H. (2018) Efficient secretion of three fungal laccases from *Saccharomyces cerevisiae* and their potential for decolorization of textile industry effluent-A comparative study. Biotechnol. Prog. 34, 69–80.

(37) DiCosimo, R., McAuliffe, J., Poulose, A. J., and Bohlmann, G. (2013) Industrial use of immobilized enzymes. Chem. Soc. Rev. 42, 6437.

(38) Shah, N., Ul-Islam, M., Khattak, W. A., and Park, J. K. (2013) Overview of bacterial cellulose composites: A multipurpose advanced material. Carbohydr. Polym. 98, 1585–1598.

(39) Wu, S.-C., and Lia, Y.-K. (2008) Application of bacterial cellulose pellets in enzyme immobilization. J. Mol. Catal.. Enzym. 54, 103–108.

(40) Wu, S.-C., Wu, S.-M., and Su, F.-M. (2017) Novel process for immobilizing an enzyme on a bacterial cellulose membrane through repeated absorption. J. Chem. Technol. Biotechnol. 92, 109–114.

(41) Chen, L., Zou, M., and Hong, F. F. (2015) Evaluation of Fungal Laccase Immobilized on Natural Nanostructured Bacterial Cellulose. Front. Microbiol. 6, 1245.

(42) Viswanath, B., Rajesh, B., Janardhan, A., Kumar, A. P., and Narasimha, G. (2014) Fungal laccases and their applications in bioremediation. Enzyme Res. 2014.

(43) Allen, H. K., Donato, J., Wang, H. H., Cloud-Hansen, K. A., Davies, J., and Handelsman, J. (2010) Call of the wild: antibiotic resistance genes in natural environments. Nat. Rev. Microbiol. 8, 251–259.

(44) Crofts, T. S., Wang, B., Spivak, A., Gianoulis, T. A., Forsberg, K. J., Gibson, M. K., Johnsky, L. A., Broomall, S. M., Rosenzweig, C. N., Skowronski, E. W., Gibbons, H. S., Sommer, M. O. A., and Dantas, G. (2018) Shared strategies for β-lactam catabolism in the soil microbiome. Nat. Chem. Biol. 14, 556–564.

(45) Baldwin, W., and Kubitschek, H. E. (1984) Buoyant Density Variation During the Cell Cycle of Saccharomyces cerevisiae. J. Bacteriol. 158, 701–704.

(46) Clasen, C., Sultanova, B., Wilhelms, T., Heisig, P., and Kulicke, W.-M. (2006) Effects of Different Drying Processes on the Material Properties of Bacterial Cellulose Membranes. Macromol. Symp. 244, 48–58.

(47) Villares, A., Moreau, C., Bennati-Granier, C., Garajova, S., Foucat, L., Falourd, X., Saake, B., Berrin, J.- G., and Cathala, B. (2017) Lytic polysaccharide monooxygenases disrupt the cellulose fibers structure. Sci. Rep. 7, 40262.

(48) Lee, C.-R., Sung, B. H., Lim, K.-M., Kim, M.-J., Sohn, M. J., Bae, J.-H., and Sohn, J.-H. (2017) Co-fermentation using Recombinant Saccharomyces cerevisiae Yeast Strains Hyper-secreting Different Cellulases for the Production of Cellulosic Bioethanol. Sci. Rep. 7, 4428.

(49) Bhagia, S., Dhir, R., Kumar, R., and Wyman, C. E. (2018) Deactivation of Cellulase at the Air-Liquid Interface Is the Main Cause of Incomplete Cellulose Conversion at Low Enzyme Loadings. Sci. Rep. 8, 1350.

(50) Yamanaka, S., Watanabe, K., Kitamura, N., Iguchi, M., Mitsuhashi, S., Nishi, Y., and Uryu, M. (1989) The structure and mechanical properties of sheets prepared from bacterial cellulose. J. Mater. Sci. 24, 3141–3145.

(51) Soykeabkaew, N., Sian, C., Gea, S., Nishino, T., and Peijs, T. (2009) All-cellulose nanocomposites by surface selective dissolution of bacterial cellulose. Cellulose 16, 435–444.

(52) Shi, X., Zheng, F., Pan, R., Wang, J., and Ding, S. (2014) Engineering and Comparative Characteristics of Double Carbohydrate Binding Modules as a Strength Additive for Papermaking Applications. BioResources 9, 3117–3131.

(53) Butchosa, N., Leijon, F., Bulone, V., and Zhou, Q. (2019) Stronger cellulose microfibril network structure through the expression of cellulose-binding modules in plant primary cell walls. Cellulose 26, 3083–3094.

(54) McIsaac, R. S., Gibney, P. A., Chandran, S. S., Benjamin, K. R., and Botstein, D. (2014) Synthetic biology tools for programming gene expression without nutritional perturbations in Saccharomyces cerevisiae. Nucleic Acids Res. 42, 1–8.

(55) McIsaac, R. S., Silverman, S. J., McClean, M. N., Gibney, P. A., Macinskas, J., Hickman, M. J., Petti, A. A., and Botstein, D. (2011) Fast-acting and nearly gratuitous induction of gene expression and protein depletion in *Saccharomyces cerevisiae*. Mol. Biol. Cell (Boone, C., Ed.) 22, 4447–4459.

(56) Pothoulakis, G., and Ellis, T. (2018) Synthetic gene regulation for independent external induction of the Saccharomyces cerevisiae pseudohyphal growth phenotype. Commun. Biol. 1, 7.

(57) Ostrov, N., Jimenez, M., Billerbeck, S., Brisbois, J., Matragrano, J., Ager, A., and Cornish, V. W. (2017) A modular yeast biosensor for low-cost point-of-care pathogen detection. Sci. Adv. 3, e1603221.

(58) Shaw, W. M., Yamauchi, H., Mead, J., Wigglesworth, M., Ladds, G., and Correspondence, T. E. (2019) Engineering a Model Cell for Rational Tuning of GPCR Signaling. Cell 177, 782–796.e27.

(59) Jarque, S., Bittner, M., Blaha, L., and Hilscherova, K. (2016) Yeast Biosensors for Detection of Environmental Pollutants: Current State and Limitations. Trends Biotechnol. 34, 408–419.

(60) Adeniran, A., Sherer, M., and Tyo, K. E. J. (2015) Yeast-based biosensors: Design and applications. FEMS Yeast Res. 15, 1–15.

(61) Cardinal-Watkins, C., and Nicell, J. A. (2011) Enzyme-Catalyzed Oxidation of 17β-Estradiol Using Immobilized Laccase from Trametes versicolor. Enzyme Res. 2011, 725172.

(62) Adeel, M., Song, X., Wang, Y., Francis, D., and Yang, Y. (2017) Environmental impact of estrogens on human, animal and plant life: A critical review. Environ. Int. 99, 107–119.

(63) Avar, P., Zrínyi, Z., Maász, G., Takátsy, A., Lovas, S., G.-Tóth, L., and Pirger, Z. (2016) β-Estradiol and ethinyl-estradiol contamination in the rivers of the Carpathian Basin. Environ. Sci. Pollut. Res. 23, 11630–11638.

(64) Pathak, G. P., Strickland, D., Vrana, J. D., and Tucker, C. L. (2014) Benchmarking of Optical Dimerizer Systems. ACS Synth. Biol. 3, 832–838.

(65) Adeniran, A., Stainbrook, S., Bostick, J., and Tyo, K. (2018) Detection of a peptide biomarker by engineered yeast receptors. ACS Synth. Biol. acssynbio.7b00410.

