## Supplement for "Living materials with programmable functionalities grown from engineered microbial co-cultures"

### Supplementary Information

#### Screening *S. cerevisiae-K. rhaeticus* co-culture conditions

To find optimal conditions for co-culture we screened a panel of conditions for *K. rhaeticus* and *S. cerevisiae* growth. Specifically, we screened growth over a range of *S. cerevisiae* inoculation ratios and in two different media: standard rich yeast medium with glucose (YPD) or sucrose (YPS) as the carbon source and standard medium for cultivation of BC-producing bacteria with glucose (HS-glucose) or sucrose (HS-sucrose) as the carbon source (Supplementary Fig. 1). Our screen led to a number of observations regarding the growth of *S. cerevisiae* and *K. rhaeticus*. Firstly, we found that, at low *S. cerevisiae* inoculation densities, co-cultures could be established in all media types. Secondly, thicker BC pellicles were obtained in yeast media (YPS and YPD) than in HS media. Thirdly, in both glucose and sucrose media, high inoculation densities of *S. cerevisiae* abolished pellicle formation, consistent with either nutrient competition or suppression of BC production by *S. cerevisiae*. Lastly, we found that, in mono-culture, *S. cerevisiae* grew well in all media types, forming a dense sediment at the base of the culture well. In contrast, in sucrose-containing media, *K. rhaeticus* grew poorly compared to glucose-containing media, failing to form a pellicle after 3 days. But, when co-cultured with *S. cerevisiae* inoculated at low density, the growth of *K. rhaeticus* in sucrose media was substantially increased, indicating that the presence of *S. cerevisiae* has some stimulatory effect on the growth of *K. rhaeticus*.

Given our aim of establishing a robust method for co-culturing *S. cerevisiae* alongside *K. rhaeticus*, the observed beneficial interaction between *K. rhaeticus* and *S. cerevisiae* in sucrose media can be considered a useful trait. Specifically, since *K. rhaeticus* growth is dependent on the growth of *S. cerevisiae*, these co-culture conditions effectively ensure that *K. rhaeticus* cannot outcompete *S. cerevisiae*. Co-culture in YPS following this protocol was therefore defined as our standard co-culture condition.

This interaction likely represents either a commensal symbiotic relationship, where one partner benefits from the interaction while the other is unaffected, or a parasitic symbiotic relationship, in which one partner benefits from the interaction while the other is detrimentally affected. A more desirable co-culture system might incorporate an obligate mutualistic symbiosis, where both species are unable to survive without the other. In this case, neither species can outcompete the other, resulting in a stable co-culture system. Although we do not explore this here, in the future it may be possible to engineer further co-dependence between *K. rhaeticus* and *S. cerevisiae*.

**Co-culture characterisation**

First, we followed a time course of pellicle formation to determine the optimal incubation time. Co-cultures yields plateaued after 3 days at a level approximately equivalent to 2.25 g/L (Supplementary Fig. 3).

Next, since yeast and bacterial communities co-exist stably over many cycles of passage during kombucha tea brewing, we wished to determine to what extent our co-culture system constitutes a similarly stable co-culture. To assay long-term co-culture dynamics, our co-culture consisted of genetically-engineered versions of *K. rhaeticus* (Kr RFP) and *S. cerevisiae* (yWS167) that respectively express red and green fluorescent proteins (RFP and GFP) and so can be individually visualised. We used a serial passage approach, in which the liquid below mature pellicles was inoculated into fresh YPS media and allowed to grow for 3 days (Supplementary Fig. 4A). This process was repeated over 16 rounds (48 days). During each round of serial passage, cultures produced new BC pellicles, confirming the presence of *K. rhaeticus* throughout serial passage (Supplementary Fig. 4B). To confirm that *S. cerevisiae* was also maintained throughout passage and to rule out the possibility of contamination with another yeast species, samples from the liquid below the pellicle and from pellicles degraded with commercial cellulase enzyme were plated onto YPD-agar and the resultant colonies imaged for GFP expression (Supplementary Fig. 4C). We observed that the original GFP-tagged *S. cerevisiae* strain, yWS167, was indeed maintained throughout the 16 rounds of serial passage. In addition, in a repeat of the passage experiment, changes in yeast cell counts and pellicle dry weight over 10 passages indicated an increase in the number of *S. cerevisiae* compared to *K. rhaeticus* (Supplementary Fig. 5).

We next wished to investigate the possible causes for the observed stimulatory effect of *S. cerevisiae* on the growth of *K. rhaeticus* in sucrose medium. Notably, *K. rhaeticus* grows well in glucose-containing medium, but much worse in similar sucrose-containing media. One possible explanation is that *S. cerevisiae* converts sucrose to a carbon source which *K.*

*rhaeticus* can consume more efficiently. Although the exact nature of the interactions between kombucha microbes remains unclear, yeasts in kombucha fermentations are known to hydrolyse the majority of carbon source, sucrose, to form extracellular glucose and fructose through the action of the secreted enzyme invertase (Supplementary Fig. 6A)[1]. To explore whether the conversion of sucrose to glucose and fructose causes the observed symbiotic interaction, we tested whether purified *S. cerevisiae* invertase could enhance *K. rhaeticus* growth in YPS medium (Supplementary Fig. 6B). As before, *K. rhaeticus* failed to produce a pellicle when grown in YPS. However, when grown in YPS spiked with invertase enzyme, *K. rhaeticus* produced thick BC pellicles after 3 days of incubation, like those produced under co-culture in YPS. This is consistent with a mechanism in which the secretion of invertase by *S. cerevisiae*, results in the accumulation of extracellular glucose and fructose which *K. rhaeticus* can more efficiently metabolise.

Another key property of our co-culture system affecting the downstream development of BC ELMs is the distribution of *S. cerevisiae* and *K. rhaeticus* between the liquid below the pellicle and the pellicle layer itself. To assess the distribution of cells, mono-cultures and co-cultures of Kr RFP bacteria and yWS167 yeast were prepared and counts of viable cells obtained from the liquid and pellicle layers. Importantly, as described in the methods section, since the degraded pellicle volume was not measured, cell counts in pellicles were estimated by assuming a fixed material volume. In all conditions, the majority of *K. rhaeticus* cells were found in the pellicle layer, while the majority of *S. cerevisiae* cells were found in the liquid layer (Fig. 1e, Supplementary Fig 7). As before, *K. rhaeticus* formed no pellicle in mono-culture in YPS. Notably, *K. rhaeticus* reached similar estimated cell densities in both the pellicle and liquid layers when grown in mono-culture in YPD and in co-culture in YPS. By contrast, *S. cerevisiae* grew to a reduced cell density when co-cultured with *K. rhaeticus* in YPS compared to mono-culture in YPS, indicating that *K. rhaeticus* either competes with *S. cerevisiae* for some nutrient in the medium or creates conditions in the co-culture that inhibit *S. cerevisiae* growth. Importantly, *S. cerevisiae* still grows to reasonably high cell densities under co-culture conditions, reaching a cell density in the liquid layer of $1.78 \times 10^7$ cells/mL ($\pm 2.42 \times 10^6$ cells/mL).

Finally, to give an idea of the robustness of our co-culture method, we set out to determine reproducibility. To achieve this, identical co-cultures were prepared following our standard protocol on three separate occasions and two parameters were measured: pellicle yields and cell counts. We found that pellicle yields tended to be consistent within triplicate samples, but variable between co-cultures set up on different occasions (Supplementary Fig. 8A). Estimated cell counts for *K. rhaeticus* were consistent in the pellicle layer, where the majority of cells were detected, but varied by up to an order of magnitude in the liquid layer (Supplementary Fig. 8C and 8E). Similarly, *S. cerevisiae* cell counts were consistent in the liquid layer, where the majority of cells were detected, but more variable in the pellicle layer (Supplementary Fig. 8D and 8F).

**Spatial patterning of catalytic living materials**

To optimise the behaviour of the light-inducible promoter system, we investigated what effect of different promoter strengths driving the expression of the DNA-binding component (LexA-CRY2) and the transcriptional activation component (VP16-CIB1) had on system performance (Supplementary Fig. 28a). We found that light-induced GFP expression exhibited the lowest background and highest expression levels when driving LexA-CRY2 expression with a weak constitutive promoter (pREV1) and VP16-CIB1 with a strong constitutive promoter (pTDH3) (Supplementary Fig. 28b). Using this information, we constructed yeast strains in which the GFP output was replaced with the luciferase enzyme, NanoLuc. Two strains were generated, each with an N-terminal MFα signal peptide and C-terminal fusions partners of either SED1 for cell surface display[2] (yNSurface strain) or CBD for cellulose binding (yNCellulose strain) (Fig. 6a and b). Notably the luciferase reporter output outperforms the previous fluorescence output, since the catalytic step greatly increases the sensitivity and reduces background noise (Supplementary Fig. 29a).

### Materials and Methods

#### Strains, constructs and DNA assembly

Strains used in this study are listed in Supplementary Table 1. All plasmids used in this study are listed in Supplementary Table 2, with links provided to their complete sequences. All plasmids constructed in this study were constructed using standard cloning techniques. Oligonucleotides were obtained from IDT. Restriction endonucleases, Phusion-HF DNA polymerase and T7 DNA ligase were obtained from NEB. Unless stated, all plasmids were transformed into *E. coli* turbo (NEB) for amplification and verification before transforming into *S. cerevisiae* for protein expression and secretion. Constructs were verified by restriction enzyme digestion and Sanger sequencing (Source Bioscience).

*S. cerevisiae* constructs and strains were generated using the yeast toolkit (YTK) system developed by the Dueber lab[3]. The YTK system uses Golden Gate assembly to combine pre-assembled, defined parts into single gene cassettes and multi-gene cassettes. The final positions of pre-assembled parts within constructs are determined by the sequences of 4 bp overhangs created by digestion with type IIS restriction enzymes (BsaI or BsmBI). Users can therefore pick and choose from pre-assembled promoter, terminator and protein-coding parts to create expression cassettes. The identity of constituent YTK DNA parts used for all single-gene and multi-gene cassettes are listed in Supplementary Tables 3 and 4, respectively. All parts were cloned into pre-assembled backbone plasmids containing genetic elements enabling cloning in *E. coli* and later integrative transformation into the URA3 locus (pYTK096) or the HO locus (pWS473) in *S. cerevisiae*. Type 2, 3 and 4 parts were cloned into pre-assembled backbones. To create more complex fusion proteins, additional subparts were used (e.g. 3a and 3b parts). New parts were codon optimised for *S. cerevisiae* expression, synthesised commercially by GeneArt or IDT and cloned into the YTK system entry vector, pYTK001, for storage and verification. All other parts were taken from the YTK or from published work. Golden Gate assembly reactions were performed as described in Lee et al.[3]

#### Culture conditions and media

Yeast extract peptone dextrose (YPD) and yeast extract peptone sucrose (YPS) media were prepared with 10 g/L yeast extract, 20 g/L peptone from soybean and 20 g/L glucose or sucrose. Synthetic complete (SC) dropout media were prepared with 1.4 g/L yeast synthetic dropout medium supplements, 6.8 g/L yeast nitrogen base without amino acids and 20 g/L glucose. Depending on the required selection, SC media were supplemented with stock solutions of one or more of uracil (final concentration 2 g/L), tryptophan (final concentration 50 mg/L), histidine (final concentration 50 mg/L) and leucine (final concentration 0.1 g/L). Hestrin–Schramm (HS) media were prepared with 5 g/L yeast extract, 5 g/L peptone from soybean, 2.7 g/L $Na_2HPO_4$, 1.5 g/L citric acid and 20 g/L glucose or sucrose. Where required, media were supplemented with 20 g/L bacteriological agar.

*E. coli* was grown in LB medium at 37°C, supplemented with appropriate antibiotics at the following concentrations: chloramphenicol 34 µg/mL, kanamycin 50 µg/mL. For biomass accumulation, *K. rhaeticus* was grown at 30°C in yeast extract peptone dextrose (YPD) medium supplemented with 34 µg/mL chloramphenicol and 1% (v/v) cellulase from *T. reesei* (Sigma Aldrich, C2730). Notably, we found that the growth of *K. rhaeticus* liquid cultures was significantly more reliable when inoculated from glycerol stock, rather than from colonies. Therefore, unless otherwise indicated, all *K. rhaeticus* cultures were inoculated from glycerol stocks. *S. cerevisiae* was grown at 30°C in rich YPD medium or selective SC medium lacking the appropriate supplements, each supplemented with 50 µg/mL kanamycin.

**Co-culture condition screen**

Triplicate samples of *K. rhaeticus* Kr RFP were inoculated from glycerol stocks into 5 mL YPD medium supplemented with cellulase (1% v/v) and grown in shaking conditions for 3 days. Triplicate samples of *S. cerevisiae* yWS167 were inoculated from plates into 5 mL YPD medium and grown in shaking conditions for 24 hours. To prepare screens, *K. rhaeticus* and *S. cerevisiae* were inoculated into 2 mL volumes of YPD, YPS, HS-glucose or HS-sucrose media in 24-well cell culture plates. *K. rhaeticus* cultures were diluted 1/50 into fresh media. *S. cerevisiae* cultures were inoculated over a range of dilutions: 1/100, 1/1000, 1/10,000, 1/100,000 and 1/1,000,000. To enable pellicle formation, plates were incubated for 4 days under static conditions at 30°C. After 4 days of incubation, cultures were photographed under identical conditions. Where present, pellicle layers were removed from the culture surface and photographed.

**OptiPrep concentration screen**

Triplicate samples of *K. rhaeticus* Kr were inoculated from glycerol stocks into 5 mL YPD medium supplemented with cellulase (1% v/v) and grown in shaking conditions for 3 days. Triplicate samples of *S. cerevisiae* BY4741 were inoculated from plates into 5 mL YPD medium and grown in shaking conditions for 24 hours. To prepare screens, *K. rhaeticus* and *S. cerevisiae* were inoculated into 10 mL volumes of YPS or YPS plus 0%, 25%, 35%, 45%, 55%, 65% (w/v) OptiPrep in 10 mLtubes. *K. rhaeticus* cultures were diluted 1/50 into fresh YPS media. *S. cerevisiae* cultures were inoculated at dilution of 1/10,000. To enable pellicle formation, tubes were incubated for 3 days under static conditions at 30°C. After 3 days of incubation, cultures were photographed under identical conditions.

**OptiPrep co-culture condition screen**

Triplicate samples of *K. rhaeticus* Kr were inoculated from glycerol stocks into 5 mL YPD medium supplemented with cellulase (1% v/v) and grown in shaking conditions for 3 days. Triplicate samples of *S. cerevisiae* BY4741 were inoculated from plates into 5 mL YPD medium and grown in shaking conditions for 24 hours. To prepare screens, *K. rhaeticus* and *S. cerevisiae* were inoculated into 10 mL volumes of YPS or YPS plus 40% (w/v) OptiPrep in 50 mL tubes with breathable caps. *K. rhaeticus* cultures were diluted 1/50 into fresh YPS media. *S. cerevisiae* cultures were inoculated over a range of dilutions: final $OD_{600}$ = 1/100, 1/1000, 1/10,000, 1/100,000 and 1/1,000,000. To enable pellicle formation, tubes were incubated for 3 days under static conditions at 30°C. After 3 days of incubation, cultures were photographed under identical conditions. Pellicles were collected and washed/shaken in deionized water at 4°C for 12 hours, twice, Clean pellicles were subjected to lyophilization for 3 days and measured for dry weight. To remove *S. cerevisiae* and *K. rhaeticus* cells associated with the cellulosic matrix, pellicles were immersed in 0.1 M NaOH at 65°C for 4 hours. Washed pellicles were again washed/shaken in $H_2O$ at 4°C for 6 hours, subjected to lyophilization for 3 days, and measured for dry weight.

### Standard co-culture protocol

Unless otherwise stated, all co-cultures were prepared using *K. rhaeticus* Kr RFP and *S. cerevisiae* yWS167 strains, allowing facile strain-specific detection through fluorescence measurements. Triplicate samples of *K. rhaeticus* were inoculated from glycerol stocks into 5 mL YPD medium supplemented with cellulase (1% v/v) and grown in shaking conditions for 3 days. Triplicate samples of *S. cerevisiae* were inoculated from plates into 5 mL YPD medium and grown in shaking conditions for 24 hours. To enable inoculation of co-cultures with equivalent cell densities of different samples, $OD_{600}$ measurements were made and used to normalise pre-culture densities. *K. rhaeticus* pre-cultures were centrifuged at 3220 x g for 10 min and cell pellets resuspended in sufficient volume of YPS medium to result in a final $OD_{600}$ of 2.5. *S. cerevisiae* pre-cultures were diluted in YPS medium to a final $OD_{600}$ of 0.01. To prepare final co-cultures, resuspended *K. rhaeticus* samples were diluted 1/50 and pre-diluted *S. cerevisiae* samples were diluted 1/100 into fresh YPS medium. In instances where strains were inoculated into various different final media, *K. rhaeticus* pellets were resuspended in PBS buffer and *S. cerevisiae* cultures were pre-diluted in PBS buffer. To prepare OptiPrep-containing co-cultures, OptiPrep (D1556, Sigma-Aldrich) was added to YPS media to a final concentration of 45% (v/v). Co-cultures were grown in either 55 mm petri dishes (15 mL) or 12 well cell culture plates (4 mL). Co-cultures were incubated for 3 days at 30°C under static conditions. It is important to ensure that culture vessels are not mechanically disturbed during the incubation period as this can partially submerge the growing BC layer, resulting in the formation of multiple, disconnected BC layers.

### Determining BC pellicle yields

To determine the yields of BC pellicles, pellicle layers were removed from the surfaces of cultures and dried using the 'sandwich method'. Here, pellicles were sandwiched between sheets of greaseproof paper and then further sandwiched between multiple sheets of absorbent paper and finally placed under a heavy weighted object. After 24 hours, fresh sheets of absorbent paper were added and pellicles left for an additional 24 hours. Pellicles dried in this way were then weighed to determine pellicle yields. Importantly, pellicles were not treated with NaOH to lyse and remove cells embedded within the BC matrix. Since the cellular biomass constitutes an integral, functional part of BC-based ELMs is made up of, we chose not to perform NaOH washes.

This method was used to follow the yields of pellicle formation over time. Here, multiple co-cultures were prepared in triplicate using the standard co-culture procedure. Co-cultures were grown in 12 well plate format. At indicated time points, pellicle layers were removed to be dried and weighed.

**Co-culture passage**

To test whether co-cultures could be passaged, initial co-cultures were prepared in triplicate in 15 mL YPS cultures using the standard co-culture protocol. After 3 days incubation at 30°C, photographs were taken of the resultant cultures. To initiate new rounds of growth, pellicle layers were removed and the liquid below mixed by aspiration then diluted 1/100 into fresh samples of 15 mL YPS. This process was repeated over 16 rounds.

To confirm that the initial strain of GFP-expressing *S. cerevisiae* (yWS167) was maintained during passage, samples were plated at the end of each round. Samples from both the liquid below the pellicle and the pellicle layer itself were plated at various dilutions onto YPD-kanamycin plates. To enable plating, pellicles were first digested by shaking gently for 16 hours at 4°C in 15 mL of PBS buffer with 2% (v/v) cellulase from *T. reesei* (Sigma Aldrich, C2730). After 48 hours of incubation at 30°C, plates were imaged for GFP fluorescence. Dilutions were selected which enabled visualisation of single colonies. Initially plates were imaged using a Fujifilm FLA-5000 Fluorescent Image Analyser. However, due to equipment malfunction, later plates were photographed under a transluminator.

**Determining cell distribution in co-cultures**

Cell distributions were determined by plating samples of cells onto solid media and counting the resultant colonies. Pellicle samples were first gently rinsed by inverting ten times in 15 mL PBS and then digested by shaking gently for 16 hours at 4°C in 15 mL of PBS buffer with 2% (v/v) cellulase from *T. reesei* (Sigma Aldrich, C2730). Samples were diluted at various levels into PBS. For *S. cerevisiae* cell counts, samples were plated onto YPD-kanamycin media.

For *K. rhaeticus* cell counts, samples were plated onto SC media lacking all four supplements essential for *S. cerevisiae* yWS167 growth (histidine, leucine, tryptophan and uracil). In all instances, Kr RFP and yWS167 strains were used. Despite the use of selective growth conditions, to ensure the colonies counted were the target strains, plates were scanned for fluorescence using a Fujifilm FLA-5000 Fluorescent Image Analyser. Plate cell counts were used to calculate the original colony forming units (cfu) per unit volume for liquid samples. However, since the exact volumes of pellicle were not measured prior to degradation, it was not possible to calculate the exact cell counts in cfu per unit volume. To enable a rough approximation of the cell counts per unit volume, pellicle volumes were estimated at fixed levels and these values were used to calculate estimated cfu per unit volume. For 15 mL cultures, pellicle volumes were estimated at 4 mL and for 4 mL cultures in 12 well plates pellicle volumes were estimated at 1 mL.

To compare cell counts from mono-cultures and co-cultures of *K. rhaeticus* and *S. cerevisiae*, pre-cultures of *K. rhaeticus* Kr RFP were pelleted and resuspended in PBS buffer and pre-cultures of *S. cerevisiae* yWS167 were diluted in PBS buffer, according to the standard co-culture procedure. Various co-cultures and mono-cultures were then prepared in different media in 15 mL volumes. After 3 days incubation at 30°C, pellicle and liquid samples were prepared, diluted and plated for cell counts.

To determine the reproducibility of co-culture cell counts, co-cultures were prepared according to the standard co-culture protocol in 15 mL cultures on three separate occasions. After 3 days incubation at 30°C, pellicle and liquid samples were prepared, diluted and plated for cell counts.

**Invertase supplementation experiment**

Co-cultures and *K. rhaeticus* Kr RFP mono-cultures were prepared in YPS medium according to the standard co-culture procedure. Recombinant, purified *S. cerevisiae* invertase (Sigma-Aldrich, I9274) was resuspended in 100 mM citrate buffer, pH 4.5 to create a stock solution at a final concentration of 5 U/μL. This stock solution was diluted into YPS medium for a range of final invertase concentrations: 50 mU/mL ($10^{-2}$), 5 mU/mL ($10^{-3}$), 0.5 mU/mL ($10^{-4}$), 50 μU/mL ($10^{-5}$). After 3 days growth at 30 °C, cultures and, where present, pellicles were imaged.

**Supernatant nitrocefin assay**

For culture supernatant assays, WT BY4741, yCG04 and yCG05 *S. cerevisiae* strains were grown in triplicate overnight in YPD liquid medium with shaking. After 16 hours growth, liquid cultures were back-diluted to final $OD_{600}$ = 0.01 in 5 mL fresh YPS medium and grown for 24 hours with shaking. The resultant cultures were centrifuged at 3220 x g for 10 min and the supernatant fractions harvested. Supernatant samples were pipetted in 50 μL volumes into the wells of a 96 well plate. The colorimetric substrate, nitrocefin (484400, Merck-Millipore), was resuspended in DMSO to create a 10 mg/mL working stock. This stock was diluted to 50 μg/mL in nitrocefin assay buffer (50 mM sodium phosphate, 1 mM EDTA, pH 7.4). To start the reaction, 50 μL of nitrocefin at 50 μg/mL was added to each of the samples simultaneously and the absorbance at 490 nm was measured over time. Active β-lactamase converts nitrocefin to a red substrate, increasing the absorbance of light at 490 nm. Therefore, to calculate the relative β-lactamase activity in samples, the rate of change in the absorbance of light at 490 nm was determined. Specifically, the product formation rates were calculated from the gradient over the linear region of a graph plotting fluorescence AU against time.

**Pellicle nitrocefin assays**

For initial pellicle assays (Fig. 2d and 2e), WT BY4741, yCG04 and yCG05 *S. cerevisiae* strains were co-cultured with *K. rhaeticus* (Kr RFP) in triplicate, according to the standard co-culture protocol. Following 3 days growth, pellicles were removed and washed in 15 mL PBS buffer for 30 min with shaking at 150 rpm. Square pieces of pellicle, measuring 5 mm x 5 mm, were then cut using a scalpel. The remainder of the pellicle was dried using the sandwich method. Once dried, pellicles were again cut to produce 5 mm x 5 mm pieces. Dried pellicle pieces were rehydrated by adding 25 μL of PBS buffer and incubating for 30 min. Assays for both wet and dried samples were run by adding 10 μL of nitrocefin, diluted to 1 mg/mL in PBS buffer, to each of the pellicle pieces simultaneously. Initial assays were performed at room temperature. Photographs were taken of pellicles over the course of 35 min to follow the colour change. To provide a quantitative measure of colour change, the ImageJ (NIH) image analysis software was used. Images were first split into individual colour channels. Since yellow-to-red colour change is caused by an increase in the absorbance of green light wavelengths, the green channel was selected. To quantify the yellow-to-red colour change, the green channel intensity was then measured from greyscale-inverted images of pellicle slices over time. Since preliminary results showed that WT pellicles exhibited no colour change, the signal from WT pellicles was used as a baseline value to correct for background levels of green channel intensity.

To determine absolute levels of β-lactamase activity in wet and dried pellicles, a similar protocol was used to create standard curves. Standard curves were prepared using a commercial *E. coli* β-lactamase enzyme (ENZ-351, ProSpec). First, pellicles grown with WT BY4741 *S. cerevisiae* were washed in nitrocefin assay buffer (50 mM sodium phosphate, 1 mM EDTA, pH 7.4). Pellicle pieces measuring 5 mm x 5 mm were cut and weighed to enable determination of the approximate volume of liquid within the pellicle. The remainder of the pellicles were dried using the sandwich method. Once dried, 5 mm x 5 mm pieces of pellicle were cut for dried pellicle standard curves. Dried pellicle pieces were rehydrated by adding 20 μL of nitrocefin assay buffer. Pre-diluted standard β-lactamase samples were then added to pellicle pieces in 5 μL volumes and allowed to diffuse throughout the BC for 30 min. To initiate the reaction, 5 μL aliquots of nitrocefin, diluted to 2 mg/mL in nitrocefin assay buffer, were added to each of the pellicle pieces simultaneously. Samples were incubated at 25°C and photographs taken over the course of the reaction. Again, ImageJ was used to quantify the yellow-to-red colour change at given time points. Time points were chosen to maximise the dynamic range, without reaching saturation. For wet pellicles, it was necessary to use measured weight of pellicle slices to determine the actual final concentration of the standard β-lactamase. Standard curves are shown in Supplementary Fig. 12. Standard curves using fresh wet pellicles, dried pellicles and dried pellicles stored for 1 month or 6 months at room temperature were all prepared according to this method. For long-term storage, dried pellicles were stored in petri dishes at room temperature and protected from light.

Alongside standard curves, pellicles grown with yCG05 *S. cerevisiae* were analysed using an identical protocol. To enable cross comparison with standard curves, negative samples (pellicles from co-cultures with WT *S. cerevisiae*) and positive samples (pellicles from co-cultures with WT *S. cerevisiae* to which a known amount of β-lactamase standard had been added) were run with samples. For samples to which no standard β-lactamase was added, 5 µL of nitrocefin assay buffer was added to maintain equal final liquid volumes. Photographs taken at identical time points were then used with standard curves to calculate absolute values of β-lactamase activity. Again, ImageJ was used to quantify the yellow-to-red colour change. For wet pellicles, it was necessary to use the measured weight of pellicle slices to determine the actual final concentration of enzyme. Again, fresh wet pellicles, dried pellicles and dried pellicles stored for 1 month at room temperature were all assayed according to this method.

**β-lactamase activity retention assay**

To determine the retention of β-lactamase within BC following multiple rounds of washes, nitrocefin assays were performed. Pieces measuring 5 mm x 5 mm were cut from dried pellicles grown with yCG04 and yCG05. All pellicle pieces were rehydrated by incubating in 1 mL of PBS buffer. Pieces were subjected to a variable number of wash steps, where pellicle pieces were incubated in 4 mL PBS buffer at 25°C and 150 rpm for 30 min. After washing, pellicles were assayed for β-lactamase activity. Negative samples (pellicles from co-cultures with WT *S. cerevisiae*) and positive samples (pellicles from co-cultures with WT *S. cerevisiae* to which a known amount of β-lactamase standard had been added) were run alongside all samples. For samples to which no standard β-lactamase was added, 5 µL of PBS buffer was added to maintain equal final liquid volumes. As before, assays were initiated by adding 5 µL of nitrocefin, diluted to 2 mg/mL in PBS buffer, to each of the pellicle pieces simultaneously. Samples were run in batches based on the number of washes. Again, ImageJ was used to quantify the yellow-to-red colour change at given time points. To enable cross-comparison between different assay runs, negative samples were used to subtract background signals and positive samples were used to normalise signals. To ensure that yellow-to-red colour change values were within a range in which there is a linear relationship between β-lactamase activity and the yellow-to-red colour change signal, a standard curve was run. The standard curve ($r^2$ = 0.9571) confirmed that detected yellow-to-red colour change values fell within the linear range.

**X-$\alpha$-gal $\alpha$-galactosidase assays**

A stock solution of X-$\alpha$-galactosidase (Sigma-Aldrich, 16555) was prepared in DMSO at a concentration of 40 mg/mL. For plate assays, 100 µL of X-$\alpha$-gal were spread on plates prior to cell plating and images taken after 3 days growth at 30 °C. For pellicle assays, pellicles grown with *K. rhaeticus* Kr RFP were harvested after 3 days growth following the standard co-culture procedure. Pellicles were then washed in 15 mL 100 mM citrate buffer, pH 4.5 for 30 min with shaking at 150 rpm. Square pieces of pellicle, measuring 5 mm x 5 mm, were then cut using a scalpel. The remainder of the pellicle was dried using the sandwich method. Once dried, pellicles were again cut to produce 5 mm x 5 mm pieces. Dried pellicle pieces were rehydrated by adding 25 µL of 100 mM citrate buffer, pH 4.5 and incubating for 30 min. Assays for both wet and dried samples were run by adding 2.5 µL of X-$\alpha$-gal stock solution and incubating at 25 °C. Images were taken over the course of several hours.

**ABTS laccase activity assays**

Stock solutions were prepared of 2,2'-azino-bis(3-ethylbenzothiazoline-6-sulphonic acid) (ABTS) (Sigma-Aldrich, A1888) at a final concentration of 0.1 M and copper sulphate at a final concentration of 1 M. Laccases are copper-containing enzymes, requiring supplementation of copper for culture and assay conditions. For plate assays, 125 µL of 0.1 M ABTS and 25 µL of 1 M $CuSO_4$ were spread on plates prior to cell plating and images taken after 3 days growth at 30 °C. For pellicle assays, pellicles grown with *K. rhaeticus* Kr RFP were harvested after 3 days growth following the standard co-culture procedure. The only modification was the addition of 1 mM CuSO4 to the culture medium of both *S. cerevisiae* pre-cultures and co-cultures. Pellicles were then washed in 15 mL 100 mM citrate buffer, 1 mM $CuSO_4$, pH 4.5 for 30 min with shaking at 150 rpm. Square pieces of pellicle, measuring 5 mm x 5 mm, were then cut using a scalpel. The remainder of the pellicle was dried using the sandwich method. Once dried, pellicles were again cut to produce 5 mm x 5 mm pieces. Dried pellicle pieces were rehydrated by adding 25 µL of 100 mM citrate buffer, 1 mM $CuSO_4$, pH 4.5 and incubating for 30 min. Assays for both wet and dried samples were run by adding 5 µL of ABTS stock solution and incubating at 25 °C. Images were taken over the course of several hours.

**Scanning electron microscopy (SEM)**

Pellicles were grown for 3 days following the co-culture procedure and washed with deionized water 3 times (shaking at 70 rpm at 4°C for 12 hours per wash) to remove residue YPS or OptiPrep. Washed pellicles were then free-dried with a lyophilizer for at least 48 hours before coated with a gold sputter. Images were taken with a JEOL 6010LA benchtop scanning electron microscope.

**Environmental scanning electron microscopy (eSEM)**

Pellicles were grown and washed as described in the SEM sample preparation. Instead of subjected to freeze-drying, washed pellicles were placed in 6-well plater suspended in transfer buffer containing 2.5% glutaraldehyde (10% EM grade from EMS, 16100) and 200 mM sodium cacodylate buffer at pH 7.2 (EMS, 11655). After 60 minutes of fixation, samples were rinsed twice with the cacodylate buffer, 5-10 minutes each at 4°C, followed by 3-4 times rinsing with distilled water. Rinsed samples should then be subjected to dehydration as soon as possible, where they were serially dehydrated with multiple rounds of EtOH, 5 minutes each (35%, 45%, 55%, 65%, 70%, 85%, 95%, and 100%). After dehydration, samples were transferred to 50% TMS (EMS, 21760) mixed with 50% EtOH and incubated for 15 minutes, followed by transferring to 80% TMS mixed with 20% EtOH and another 15-minute incubation. Finally, samples were transferred to 100% TMS and incubated for 5 minutes, repeated for 3 times, and air-dried overnight in the fume hood. Images were taken with a FEI XL30 ESEM used on low vacuum mode with Backscatter detector (BSE). Dehydrated samples were placed on stub using double-sided conductive carbon tape (EMS). Parameters: Working height < 10 cm; Low pressure setting > 2.5 Torr; Accelerating voltage 15 kV; Magnification > 1000x; Spot size = 3.

**Brunauer-Emmett-Teller (BET) surface area analysis**

Free-dried pellicles were cut into 5 mm x 5 mm piece and placed in sample tube for 1 hour degas at 423 K using a Micromeritics (Atlanta, GA) ASAP 2020 analyser. BET surface area and pore size were then determined with $N_2$ adsorption at 77 K using Brunauer–Emmett–Teller and Barrett–Joyner–Halenda analyses on the same machine.

**Preparing and assaying sense-and-response pellicles**

In yGPH093, transcription from the BED-inducible promoter is controlled by a synthetic transcription factor ($Z_3EV$) consisting of three domains: the Zif268 DNA-binding domain, the human estrogen receptor (hER) ligand binding domain, and the transcriptional activation domain of viral protein 16 ($VP16^{AD}$)[4]. When present, β-estadiol binds to the hER ligand binding domain of $Z_3EV$, releasing it from its basal sequestration in the cytosol and enabling it to translocate into the nucleus. Once in the nucleus, the Zif268 domain binds cognate DNA sequences in engineered promoters and the $VP16^{AD}$ domain activates transcription of downstream genes. As a preliminary test of *S. cerevisiae* sense-and-response in BC pellicles, co-cultures were prepared in triplicate according to the standard co-culture protocol using WT BY4741 and yGPH093 strains. Co-cultures were inoculated into 4 mL YPS-OptiPrep medium in 12 well cell culture plates. After 3 days of growth, pellicles were removed and washed by incubating at 25°C with shaking at 150 rpm in 15 mL PBS. Pellicles were then placed in fresh 15 mL of YPD medium in the presence or absence of 5 nM β-estradiol (E8875, Sigma-Aldrich) and incubated for 24 hours at 30°C and 150 rpm. During growth cells had 'escaped' from biosensor pellicles, making the medium surrounding the pellicles turbid. Therefore, to remove loosely-associated cells, pellicles were washed twice by incubating at for 30 min at 25°C and 150 rpm in 15 mL of PBS buffer. Finally, pellicles were imaged simultaneously for GFP fluorescence under a transilluminator.

Similarly, dried biosensor pellicles were prepared in triplicate according to the standard co-culture protocol using WT BY4741 and yGPH093 or WT BY4741 and yWS890 strains. Co-cultures were inoculated into 4 mL YPS-OptiPrep medium in 12 well cell culture plates. After 3 days of growth, pellicles were dried using the 'sandwich method'. Dried pellicles were then placed in fresh 15 mL of YPD medium in the presence or absence of 5 nM β-estradiol or 50 nM *S. cerevisiae* α–mating factor (RP01002, Genscript) and incubated 24 hours at 30 °C. Notably, to more closely match the potential use of biosensors in an on-site detection setting, pellicles were incubated without agitation in this and all future experiments. Since static growth results in far less growth in the surrounding liquid, pellicles were only briefly washed after incubation by inverting ten times in 15 mL PBS buffer. Finally, pellicles were imaged side-by-side for GFP fluorescence under a transilluminator. To test for stability after long-term storage, pellicles were stored for 4 months at room temperature stored in petri dishes protected from light. These pellicles were cut in half prior to induction, which was performed as above.

The BED-inducible CtLcc1-secreting strain yCG23 was initially screened for laccase induction using a plate-based ABTS assay. Transformants of yCG23 were re-streaked in triplicate on SC $URA^-$ plates supplemented with 125 µL of 0.1 M ABTS and 25 µL of 1 M $CuSO_4$. After 3 days of incubation at 30 °C colonies were imaged. Co-cultures between *K. rhaeticus* Kr RFP and yCG01 or yCG23 were then prepared in triplicate in 12-well plate format, using YPS-OptiPrep medium supplemented with 1 mM $CuSO_4$. After 3 days growth, pellicles were harvested and were washed by incubating for 30 min at 25°C and 150 rpm in 15 mL of 100 mM citrate buffer, 1 mM $CuSO_4$, pH 4.5. Pellicles were then inoculated into 15 mL of fresh YPD supplemented with 1 mM $CuSO_4$ in the presence or absence of 5 nM β-estradiol and incubated at 30 °C for 24 hours statically. After incubation, pellicles were washed by incubating for 30 min at 25°C and 150 rpm in 15 mL of 100 mM citrate buffer, 1 mM $CuSO_4$, pH 4.5. Pellicles were then placed in a 12-well plate and 75 µL of 0.1 M ABTS added to each well to assay for laccase activity. Pellicles were incubated at 25 °C and imaged after 72 hours.

**Determining the viability of *S. cerevisiae* in dried BC pellicles**

Co-cultures were prepared in triplicate according to the standard co-culture protocol. Co-cultures were inoculated into 4 mL YPS-OptiPrep medium in 12 well cell culture plates. Counts of viable *S. cerevisiae* cells within wet and dried pellicles were determined as described in "Determining cell distribution in co-cultures". Dried pellicles were also stored for 1 month at room temperature, and then degraded and plated onto YPD medium. Since one of the triplicate samples produced no colonies, we could not calculate estimated cell counts within pellicles. However, images are presented of the three plates to show that viable cells were indeed recovered from the other two samples (Supplementary Fig. 25D).

**Total cellulase activity assay**

*S. cerevisiae* strains BY4741 and yCelMix were grown overnight in YPS in triplicate with shaking. After 16 hours growth, liquid cultures were back-diluted to final $OD_{600}$ = 0.1 in 5 mL fresh YPS medium with 2 mM L-ascorbic acid (A7506, Sigma-Aldrich) and grown for 24 hours with shaking. The resultant cultures were centrifuged at 3220 x g for 10 min and the supernatant fractions harvested. Supernatant samples were pipetted in 50 μL volumes into the wells of a 96 well plate. The EnzChek® Cellulase Substrate (E33953, Thermo-Fisher) was resuspended in 50% DMSO and diluted 5-fold in 100 mM sodium acetate (pH 5.0). To start the reaction, 50 μL of cellulase substrate was added to the supernatant and let incubated for 30 minutes in dark at room temperature. To build an enzyme activity standard curve, the cellulase from *T. reesi* (C2730, Sigma-Aldrich) was used to prepare a serial dilution in YPS medium and mixed with the substrate at 1:1 ratio. Blue fluorescence (360/460) was detected using a plate reader (Synergy H1, BioTek) after 30 minutes incubation in dark at room temperature. The data from enzyme standards was fit to an exponential model, a*exp(b*x) + c*exp(d*x) in MATLAB. This model was then used to calculate the total cellulase activity of the supernatant from yCelMix (using supernatant from BY4741 as a blank control).

**Pellicle tensile test**

Co-cultures were set up in 40 mL YPS + OptiPrep (plus 2 mM L-ascorbic acid) and grown in square plates (100 mm x 15 mm) for 2 days at 30 °C. Pellicles were then washed in deionized water 3 times (shaking at 70 rpm at 4°C for 12 hours per wash) and dried using the sandwich method described previously but with an extended 3 days drying to ensure water removal. Dried pellicles were cut into 60 mm * 10 mm stripes and their thickness were measured with a micrometer. Tensile test was performed with a Zwick mechanical tester (BTC-ExMacro .001, Roell) following the ASTM D882 protocol at 1 mm/min speed.

**Pellicle rheology analysis**

The rheological properties of washed pellicles were characterized on a rheometer (AR2000, TA Instruments) with a 25 mm ETC aluminium plate (1 mm gap). The strain sweep measurements were taken from 0.01% to 100% strain amplitude at a constant frequency of 1 rad/s while frequency sweep measurements were taken from 0.1 rad/s to 100 rad/s at a constant strain amplitude of 1%. Samples were kept fully-hydrated with deionized water at 25 °C on a Peltier thermoelectric plate.

**Light-inducible circuit promoter characterization**

Yeast strains were grown overnight in YPD in triplicate with shaking. After 16 hours growth, liquid cultures were back-diluted to final $OD_{600}$ = 0.2 in 100 µL fresh YPD and pipetted into the wells of two 96 well plates (duplicates). One of the two plates was wrapped in black aluminium foil as a dark control. Both plates were placed under a LED lamp at 30 °C for 4 hours. Green fluorescence was then measured with a plate reader.

**Light-inducible luciferase assay**

Yeast strains were grown overnight in YPD in triplicate with shaking. After 16 hours growth, liquid cultures were back-diluted to final $OD_{600}$ = 0.2 in 15 µL fresh YPS and pipetted into the wells of two 96 well plates (duplicates for light and dark conditions, as previously described). Plates were placed under a LED lamp at 30 °C for 4 hours. Substrate in buffer from Nano-Glo® Luciferase Assay System (N1120, Promega) were added to the culture at 1:1 ratio at the end of incubation. After incubation in dark for 5 minutes, bioluminescence of the samples was measured with a plate reader.

**Light-inducible pellicle response assay**

Co-cultures were set up using yeast strains BY4741, yNCellulose, and yNSurface along with wildtype *K. rhaeticus* in 10 mL YPS + OptiPrep. For long term exposure experiment, 60 mm petri dishes were prepared as duplicates, one was wrapped in black aluminium foil while the other one was not. The plates were placed under a LED lamp at 30 °C for 3 days. After the incubation, pellicles were flipped so the bottom side was facing up, and transferred onto YPD agar plates. 500 µL of Nano-Glo mix was applied onto the pellicles evenly through the entire surface. After incubation in dark for 10 minutes, bioluminescence of the samples was detected with a ChemiDoc Touch imager (BioRad). For short term exposure experiment (masking), co-cultures were grown in dark at 30 °C for 3 days. Pellicles were flipped so the bottom side was facing up and transferred onto YPD agar plates. A mask made of black aluminium foil with carved pattern in the centre was placed on top of the pellicles. Plates were placed under a LED lamp and incubated at 30 °C for 4 hours. Mask was then removed and 500 µL of Nano-Glo mix was applied onto the pellicles evenly through the entire surface.

After incubation in dark for 10 minutes, bioluminescence of the samples was detected with a ChemiDoc Touch imager.

**Light-patterning on pellicles**

Co-cultures were grown in 100 mm square plates protected from light as previously described. Pellicles were rinsed in PBS, flipped, placed on YPD agar, and placed in an incubator with a projector mounted on top. After incubation under the projected pattern (with no lid to prevent water condensation) at 30 °C for 12 hours or more, 3 mL of Nano-Glo mix was applied onto the pellicles evenly through the entire surface. Bioluminescence images was detected with a ChemiDoc Touch imager after 30 minutes incubation in dark.

### Supplementary Figures and Tables

### Supplementary Table 1. Strains used in this study

| Strains | Description and phenotype | Source |
|---|---|---|
| *Komagataeibacter rhaeticus iGEM* | BC-producing bacterium isolated from kombucha tea | Florea et al.[5] |
| *K. rhaeticus* Kr RFP | Constitutive mRFP expression. *K. rhaeticus* transformed with J23110-mRFP1-331Bb. | Florea et al.[5] |
| Sc BY4741 | MATa his3Δ1 leu2Δ0 met15Δ0 ura3Δ0 | Dharmacon yeast collection |
| Sc YSC6273 | BY4741 sed1Δ::KanMX | Dharmacon yeast collection |
| Sc yWS167 | Constitutive GFP expression, BY4741 transformed with integrative plasmid pWS702 | This work |
| Sc yCG01 | Constitutive secretion of MFa-sfGFP, BY4741 transformed with integrative plasmid pCG01 | This work |
| Sc yCG04 | Constitutive secretion of MFa-BLA, BY4741 transformed with integrative plasmid pCG04 | This work |
| Sc yCG05 | Constitutive secretion of MFa-BLA-CBDcex, BY4741 transformed with integrative plasmid pCG05 | This work |
| Sc yCG16 | Constitutive secretion of MtLcc1 signal peptide-MtLcc1-CBDcex, BY4741 transformed with integrative plasmid pCG16 | This work |
| Sc yCG17 | Constitutive secretion of MFa signal peptide-MtLcc1-CBDcex, BY4741 transformed with integrative plasmid pCG17 | This work |
| Sc yCG18 | Constitutive secretion of CtLcc1 signal peptide-CtLcc1-CBDcex, BY4741 transformed with integrative plasmid pCG18 | This work |
| Sc yCG19 | Constitutive secretion of MFa signal peptide-CtLcc1-CBDcex, BY4741 transformed with integrative plasmid pCG19 | This work |
| Sc yCG20 | Constitutive secretion of Mel1 signal peptide-Mel1-CBDcex, BY4741 transformed with integrative plasmid pCG20 | This work |
| Sc yCG21 | Constitutive secretion of MFa signal peptide-Mel1-CBDcex, BY4741 transformed with integrative plasmid pCG21 | This work |
| Sc yCG23 | β-estradiol inducible secretion of CtLcc1 laccase. BY4741 transformed with integrative plasmid pCG23 | This work |
| Sc yGPH093 | β-estradiol inducible GFP expression. BY4741 transformed with integrative plasmid pGPY093 | Pothoulakis et al.[6] |
| Sc yWS890 | Alpha-factor-inducible GFP expression. "Design 4" strain from Shaw el at. (2019): BY4741 sst2Δ0 far1Δ0 bar1Δ0 ste2Δ0 ste12Δ0 gpa1Δ0 ste3Δ0 mf(alpha)1Δ0 mf(alpha)2Δ0 mfa1Δ0 mfa2Δ0 gpr1Δ0 gpa2Δ0 LexO(6x)-pLEU2m-sfGFP-tTDH1-LEU2 pCCW12-STE2-tSSA1-pPGK1-GPA1-tENO2-pRAD27-LexA-PRD-tENO1-URA3 | Shaw et al.[7] |
| Sc yCelMix | Constitutive secretion of CtCbh1, ClCbh2, SfBgl1, and PaLPMO9H, BY4741 transformed with integrative plasmid pCelMix | This work |
| Sc yCBH1 | Constitutive secretion of CtCbh1, BY4741 transformed with integrative plasmid pCBH1 | This work |
| Sc yCBH2 | Constitutive secretion of ClCbh2, BY4741 transformed with integrative plasmid pCBH2 | This work |
| Sc yBGL1 | Constitutive secretion of SfBgl1, BY4741 transformed with integrative plasmid pBGL1 | This work |
| Sc yEGL2 | Constitutive secretion of TrEgl2, BY4741 transformed with integrative plasmid pEgl2 | This work |
| Sc yLPMO | Constitutive secretion of PaLPMO9H, BY4741 transformed with integrative plasmid pLPMO | This work |

| | | |
|---|---|---|
| Sc yXTH3 | Constitutive secretion of AtXth3, BY4741 transformed with integrative plasmid pXTH3 | This work |
| Sc yTT-GFP | Blue light inducible GFP expression. BY4741 transformed with integrative plasmid pTT-GFP | This work |
| Sc yRR-GFP | Blue light inducible GFP expression. BY4741 transformed with integrative plasmid pRR-GFP | This work |
| Sc yTR-GFP | Blue light inducible GFP expression. BY4741 transformed with integrative plasmid pTR-GFP | This work |
| Sc yRT-GFP | Blue light inducible GFP expression. BY4741 transformed with integrative plasmid pRT-GFP | This work |
| Sc yNCellulose | Blue light inducible NanoLuc-CBD secretion. BY4741 transformed with integrative plasmid pNCellulose | This work |
| Sc yNSurface | Blue light inducible NanoLuc surface display. BY4741 transformed with integrative plasmid pNSurface | This work |
| Sc yNSurfaceΔSED1 | Blue light inducible NanoLuc surface display. YSC6273 transformed with integrative plasmid pNSurface | This work |
| Sc yWO68 | Constitutive expression of mScarlet-I, BY4741 (ΔAga1, ΔAga2) transformed with integrative plasmid pWO68. | This work |

**Supplementary Table 2. Plasmids used in this study**. Links to annotated plasmid sequences are provided for all constructs.

| Plasmid | Construct details | Source |
|---|---|---|
| J23110-mRFP1-331Bb | Constitutive mRFP1 expression from the pSEVA-331Bb backbone plasmid. Expression is driven by the low strength J23110 promoter. Addgene #78277 | Florea et al.[5] |
| pYTK001 | YTK entry vector into which new DNA parts can be cloned using BsmBI golden gate reactions, verified and stored for later assemblies. | Lee et al.[3] |
| pYTK096 | Pre-assembled YTK plasmid containing genetic elements enabling cloning in E. coli and later integrative transformation into the URA3 locus in *S. cerevisiae*. Golden gate assembly allows insertion of YTK type 2-3-4 parts. | Lee et al.[3] |
| pWS041 | Pre-assembled YTK acceptor plasmid into which single gene cassettes can be cloned, consisting of an sfGFP dropout part flanked by YTK connectors ConS and Con1. Backbone enabling propagation in *E. coli* | This study |
| pWS042 | Pre-assembled YTK acceptor plasmid into which single gene cassettes can be cloned, consisting of an sfGFP dropout part flanked by YTK connectors Con1 and ConE. Backbone enabling propagation in *E. coli* | This study |
| pWS702 | YTK single gene cassette plasmid for constitutive GFP expression, pTDH3-sfGFP-tTDH1. Backbone enabling propagation in *E. coli* and markerless integration at the HO locus in *S. cerevisiae* | This study |
| pCG01 | YTK single gene cassette plasmid for constitutive sfGFP secretion, pTDH3-MFa-sfGFP-tTDH1. Backbone enabling propagation in *E. coli* and integration at the URA3 locus in *S. cerevisiae* | This study |
| pCG04 | YTK single gene cassette plasmid for constitutive BLA secretion, pTDH3-MFa-BLA-tTDH1. Backbone enabling propagation in *E. coli* and integration at the URA3 locus in *S. cerevisiae* | This study |
| pCG05 | YTK single gene cassette plasmid for constitutive BLA-CBD secretion, pTDH3-MFa-BLA-CBD-tTDH1. Backbone enabling propagation in *E. coli* and integration at the URA3 locus in *S. cerevisiae* | This study |
| pCG16 | YTK single gene cassette plasmid for constitutive MtLcc1-CBD secretion, pTDH3-MtLcc1$^{SP}$-MtLcc1-CBD-tTDH1. Backbone enabling propagation in *E. coli* and integration at the URA3 locus in *S. cerevisiae* | This study |
| pCG17 | YTK single gene cassette plasmid for constitutive MtLcc1-CBD secretion, pTDH3-MFa-MtLcc1-CBD-tTDH1. Backbone enabling propagation in *E. coli* and integration at the URA3 locus in *S. cerevisiae* | This study |
| pCG18 | YTK single gene cassette plasmid for constitutive CtLcc1-CBD secretion, pTDH3-CtLcc1$^{SP}$-CtLcc1-CBD-tTDH1. Backbone enabling propagation in *E. coli* and integration at the URA3 locus in *S. cerevisiae* | This study |
| pCG19 | YTK single gene cassette plasmid for constitutive CtLcc1-CBD secretion, pTDH3-MFa-CtLcc1-CBD-tTDH1. Backbone enabling propagation in *E. coli* and integration at the URA3 locus in *S. cerevisiae* | This study |
| pCG20 | YTK single gene cassette plasmid for constitutive Mel1-CBD secretion, pTDH3-Mel1$^{SP}$-Mel1-CBD-tTDH1. Backbone enabling propagation in *E. coli* and integration at the URA3 locus in *S. cerevisiae* | This study |
| pCG21 | YTK single gene cassette plasmid for constitutive Mel1-CBD secretion, pTDH3-MFa-Mel1-CBD-tTDH1. Backbone enabling propagation in *E. coli* and integration at the URA3 locus in *S. cerevisiae* | This study |
| pCG22 | YTK single gene cassette plasmid for expression of CtLcc1 signal peptide-CtLcc1-CBD driven by the pZ3 promoter: pZ3-CtLcc1SP-CtLcc1-CBDcex-tTDH1. Backbone enabling propagation in *E. coli* | This study |
| pCG23 | YTK multi gene cassette plasmid for β-estradiol inducible CtLcc1-CBD secretion, cassette 1: pREV1-$Z_3$E-VP16$^{AD}$-tTDH1, cassette 2: pGAL1$^{6xZ3BS}$-CtLcc1 signal peptide-CtLcc1-CBD-tTDH1. Backbone enabling propagation in *E. coli* and integration at the URA3 locus in *S. cerevisiae* | This study |

| | | |
|---|---|---|
| pGPY093 | YTK multi gene cassette plasmid for β-estradiol inducible GFP expression, cassette 1: pREV1-$Z_3$E-VP16$^{AD}$-tTDH1, cassette 2: pGAL1$^{6xZ3BS}$-sfGFP-tTDH1. Backbone enabling propagation in *E. coli* and integration at the URA3 locus in *S. cerevisiae* | Pothoulakis et al.[6] |
| pGPY074 | YTK single gene cassette plasmid for expression of $Z_3$EV driven by the weak constitutive promoter pREV1: pREV1-$Z_3$E-VP16$^{AD}$-tTDH1. Backbone enabling propagation in *E. coli* | Pothoulakis et al.[6] |
| pGPY085 | YTK single gene cassette plasmid for expression of sfGFP driven by the pZ3 promoter: pZ3-sfGFP-tTDH1. Backbone enabling propagation in *E. coli* | This study |
| pWS032 | YTK position 3b part containing the *E. coli* TEM1 beta-lactamase BLA (UniProt: Q6SJ61) mature protein, lacking signal peptide | This study |
| pWS033 | YTK position 3 part containing sfGFP | Shaw et al.[7] |
| pWS034 | YTK position 3b part containing sfGFP | This study |
| pWS433 | YTK position 3a part containing the S. cerevisiae mating-factor alpha (UniProt: P01149) signal peptide | This study |
| pWS930 | YTK position 3a part containing a fusion of the Zif268 DNA binding domain and the ligand binding domain of the human estrogen receptor | Pothoulakis et al.[6] |
| pWS935 | YTK position 3b part containing the VP16 transcriptional activation domain | Pothoulakis et al.[6] |
| pPPK027 | YTK position 4a part containing the CBDcex cellulose binding domain from the *Cellulomonas fimi* Cex exoglucanase (UniProt: P07986) | This study |
| pPPK041 | YTK position 3a part containing the *Myceliophthora thermophila* laccase Lcc1 (UniProt: Q9HDQ0) signal peptide | This study |
| pPPK042 | YTK position 3a part containing the *Coriolopsis trogii* laccase Lcc1 (UniProt: G2QG31) signal peptide | This study |
| pPPK043 | YTK position 3a part containing the *S. cerevisiae* alpha-galactosidase Mel1 (UniProt: P04824) signal peptide | This study |
| pWS1078 | YTK position 2 part containing a modified GAL1 promoter containing six Zif268 binding sequences | Pothoulakis et al.[6] |
| pPPK044 | YTK position 3b part containing the *Myceliophthora thermophila* laccase Lcc1 (UniProt: Q9HDQ0) mature protein, lacking signal peptide | This study |
| pPPK045 | YTK position 3b part containing the *Coriolopsis trogii* laccase Lcc1 (UniProt: G2QG31) mature protein, lacking signal peptide | This study |
| pPPK046 | YTK position 3b part containing the *S. cerevisiae* alpha-galactosidase Mel1 (UniProt: P04824) mature protein, lacking signal peptide | This study |
| pWS473 | Pre-assembled YTK plasmid containing genetic elements enabling cloning in E. coli and later markerless integrative transformation into the HO locus in S. cerevisiae. Golden gate assembly allows insertion of YTK type 2-3-4 parts. | This study |
| pCelMix | YTK multi gene cassette plasmid for constitutive cellulase cocktail secretion, cassette 1: pTDH3-NS-CtCbh1-tENO1, cassette 2: pCCW12-TFP13-ClCbh2-tADH1, cassette 3: pTEF1-TFP19-SfBgl1-tPGK1, cassette 4: pTEF2-TFP19-TrEgl2-tENO2, cassette 5: pPGK1-MFa-PaLPMO9H-tTDH1. Backbone enabling propagation in *E. coli* and integration at the URA3 locus in *S. cerevisiae* | This study |
| pLSR1-CBH1 | YTK single gene cassette 1 plasmid for constitutive Cbh1 secretion, pTDH3-NS-CtCbh1-tENO1 inserted onto pYTKLSR1DO | This study |
| pL1R2-CBH2 | YTK single gene cassette 2 plasmid for constitutive Cbh2 secretion, pCCW12-TFP13-ClCbh2-tADH1 inserted onto pYTKL1R2DO | This study |
| pL2R3-BGL1 | YTK single gene cassette 3 plasmid for constitutive Bgl1 secretion, pTEF1-TFP19-SfBgl1-tPGK1 inserted onto pYTKL2R3DO | This study |
| pL3R4-EGL2 | YTK single gene cassette 4 plasmid for constitutive Egl2 secretion, pTEF2-TFP19-TrEgl2-tENO2 inserted onto pYTKL3R4DO | This study |
| pL4RE-LPMO | YTK single gene cassette 5 plasmid for constitutive LPMO secretion, pPGK1-MFa-PaLPMO9H-tTDH1 inserted onto pYTKL4REDO | This study |

| | | |
|---|---|---|
| pBGL1 | YTK single gene cassette plasmid for constitutive Bgl1 secretion, pTDH3-TFP19-SfBgl1-tTDH1. Backbone enabling propagation in *E. coli* and integration at the URA3 locus in *S. cerevisiae* | This study |
| pCBH1 | YTK single gene cassette plasmid for constitutive Cbh1 secretion, pTDH3-NS-CtCbh1-tTDH1. Backbone enabling propagation in *E. coli* and integration at the URA3 locus in *S. cerevisiae* | This study |
| pCBH2 | YTK single gene cassette plasmid for constitutive Cbh2 secretion, pTDH3-TFP13-ClCbh2-tTDH1. Backbone enabling propagation in *E. coli* and integration at the URA3 locus in *S. cerevisiae* | This study |
| pEGL2 | YTK single gene cassette plasmid for constitutive Egl2 secretion, pTDH3-TFP19-TrEgl2-tTDH1. Backbone enabling propagation in *E. coli* and integration at the URA3 locus in *S. cerevisiae* | This study |
| pLPMO | YTK single gene cassette plasmid for constitutive LPMO secretion, pTDH3-MFa-PaLPMO9H-tTDH1. Backbone enabling propagation in *E. coli* and integration at the URA3 locus in *S. cerevisiae* | This study |
| pXTH3 | YTK single gene cassette plasmid for constitutive Xth3 secretion, pTDH3-MFa-AtXTH3-tTDH1. Backbone enabling propagation in *E. coli* and integration at the URA3 locus in *S. cerevisiae* | This study |
| pYTK-CBH1 | YTK position 3 part containing the *Chaetomium thermophilum* cellobiohydrolase with native signal peptide (NS-CtCbh1)[8], codon-optimized for expression in *S. cerevisiae* | This study |
| pYTK-CBH2 | YTK position 3 part containing the *Chrysosporium lucknowense* cellobiohydrolase with translational fusion partner 13 (TFP13-ClCbh2)[8], codon-optimized for expression in *S. cerevisiae* | This study |
| pYTK-BGL1 | YTK position 3 part containing the *Saccharomycopsis fibuligera* β-glucosidase with translational fusion partner 19 (TFP19-SfBgl1)[8], codon-optimized for expression in *S. cerevisiae* | This study |
| pYTK-EGL2 | YTK position 3 part containing the *Trichoderma reesei* endoglucanase with translational fusion partner 19 (TFP19-TrEgl2)[8], codon-optimized for expression in *S. cerevisiae* | This study |
| pYTK-LPMO | YTK position 3 part containing the *Podospora anserina* lytic polysaccharide monooxygenases with MFalpha signal peptide (MFa-LPMO)[9], codon-optimized for expression in *S. cerevisiae* | This study |
| pYTK-XTH3 | YTK position 3 part containing the *Arabidopsis thaliana* xyloglucan endotransglucosylase/hydrolase with MFalpha signal peptide (MFa-XTH3)[10], codon-optimized for expression in *S. cerevisiae* | This study |
| pTT-GFP | YTK multi gene cassette plasmid for light-inducible GFP expression, cassette 1: pTDH3-LexABD-CRY2-tTDH1, cassette 2: spacer, cassette 3: pLEXA-yeGFP-tTDH1, cassette 4: spacer, cassette 5: pTDH3-VP16-CIB1-tTDH1. Backbone enabling propagation in *E. coli* and integration at the URA3 locus in *S. cerevisiae* | This study |
| pRR-GFP | YTK multi gene cassette plasmid for light-inducible GFP expression, cassette 1: pREV1-LexABD-CRY2-tTDH1, cassette 2: spacer, cassette 3: pLEXA-yeGFP-tTDH1, cassette 4: spacer, cassette 5: pREV1-VP16-CIB1-tTDH1. Backbone enabling propagation in *E. coli* and integration at the URA3 locus in *S. cerevisiae* | This study |
| pTR-GFP | YTK multi gene cassette plasmid for light-inducible GFP expression, cassette 1: pTDH3-LexABD-CRY2-tTDH1, cassette 2: spacer, cassette 3: pLEXA-yeGFP-tTDH1, cassette 4: spacer, cassette 5: pREV1-VP16-CIB1-tTDH1. Backbone enabling propagation in *E. coli* and integration at the URA3 locus in *S. cerevisiae* | This study |
| pRT-GFP | YTK multi gene cassette plasmid for light-inducible GFP expression, cassette 1: pREV1-LexABD-CRY2-tTDH1, cassette 2: spacer, cassette 3: pLEXA-yeGFP-tTDH1, cassette 4: spacer, cassette 5: pTDH3-VP16-CIB1-tTDH1. Backbone enabling propagation in *E. coli* and integration at the URA3 locus in *S. cerevisiae* | This study |

| | | |
|---|---|---|
| pNCellulose | YTK multi gene cassette plasmid for light-inducible Nanoluc-CBM3 secretion, cassette 1: pREV1-LexABD-CRY2-tTDH1, cassette 2: spacer, cassette 3: pLEXA-MFa-NanoLuc-CBD-tTDH1, cassette 4: spacer, cassette 5: pTDH3-VP16-CIB1-tTDH1. Backbone enabling propagation in *E. coli* and integration at the URA3 locus in *S. cerevisiae* | This study |
| pNSurface | YTK multi gene cassette plasmid for light-inducible Nanoluc-SED1 surface display, cassette 1: pREV1-LexABD-CRY2-tTDH1, cassette 2: spacer, cassette 3: pLEXA-MFa-NanoLuc-Sed1-tTDH1, cassette 4: spacer, cassette 5: pTDH3-VP16-CIB1-tTDH1. Backbone enabling propagation in *E. coli* and integration at the URA3 locus in *S. cerevisiae* | This study |
| pLSR1-T-LexABD-CRY2 | YTK single gene cassette 1 plasmid for strong constitutive LexABD-CRY2 expression, pTDH3-LexABD-CRY2-tTDH1 inserted onto pYTKLSR1DO | This study |
| pLSR1-R-LexABD-CRY2 | YTK single gene cassette 1 plasmid for weak constitutive LexABD-CRY2 expression, pREV1-LexABD-CRY2-tTDH1 inserted onto pYTKLSR1DO | This study |
| pL1R2-spacer | YTK single gene cassette 2 plasmid where an empty spacer is inserted on pTYKL1R2DO | This study |
| pL2R3-LexA-GFP | YTK single gene cassette 3 plasmid for light-inducible GFP expression, pLEXA-yeGFP-tTDH1 inserted onto pYTKL2R3DO | This study |
| pL2R3-LexA-Nluc-CBM3 | YTK single gene cassette 3 plasmid for light-inducible NanoLuc-CBM3 secretion, pLEXA-MFa-Nluc-CBM3-tTDH1 inserted onto pYTKL2R3DO | This study |
| pL2R3-Nluc-SED1 | YTK single gene cassette 3 plasmid for light-inducible NanoLuc-SED1 surface display, pLEXA-MFa-Nluc-Sed1-tTDH1 inserted onto pYTKL2R3DO | This study |
| pL3R4-spacer | YTK single gene cassette 4 plasmid where an empty spacer is inserted on pTYKL3R4DO | This study |
| pL4RE-T-VP16-CIB1 | YTK single gene cassette 5 plasmid for strong constitutive VP16-CIB1 expression, pTDH3- VP16-CIB1-tTDH1 inserted onto pYTKL4REDO | This study |
| pL4RE-R-VP16-CIB1 | YTK single gene cassette 5 plasmid for weak constitutive VP16-CIB1 expression, pREV1- VP16-CIB1-tTDH1 inserted onto pYTKL4REDO | This study |
| pYTK-LexABD-CRY2 | YTK position 3 part containing the LexA DNA binding domain and optical dimerizing partner CRY2 (LexABD-CRY2)[11], codon-optimized for expression in *S. cerevisiae* | This study |
| pYTK-VP16-CIB1 | YTK position 3 part containing the VP16 activation domain and optical dimerizing partnerr CIB1 (VP16-CIB1)[11], codon-optimized for expression in *S. cerevisiae* | This study |
| pYTK-PLEXA | YTK position 2 part containing a modified CYC1 minimal promoter containing eight LexA binding sequences | This study |
| pYTK-yeGFP | YTK position 3 part containing yeGFP | This study |
| pYTK-Nluc | YTK position 3b part containing the NanoLuc luciferase reporter, codon-optimized for expression in *S. cerevisiae* | This study |
| pYTK-CBM3 | YTK position 4a part containing the *Hungateiclostridium thermocellum* cellulosome anchoring protein cellulose-binding module (CBM3), codon-optimized for expression in *S. cerevisiae* | This study |
| pYTK-SED1 | YTK position 4a part containing the *S. cerevisiae* Sed1 protein for surface display | This study |
| pYTKLSR1DO | Pre-assembled YTK acceptor plasmid into which single gene cassettes can be cloned, consisting of an sfGFP dropout part flanked by YTK connectors ConS and Con1. Backbone enabling propagation in *E. coli* | This study |

| | | |
|---|---|---|
| pYTKL1R2DO | Pre-assembled YTK acceptor plasmid into which single gene cassettes can be cloned, consisting of an sfGFP dropout part flanked by YTK connectors Con1 and Con2. Backbone enabling propagation in *E. coli* | This study |
| pYTKL2R3DO | Pre-assembled YTK acceptor plasmid into which single gene cassettes can be cloned, consisting of an sfGFP dropout part flanked by YTK connectors Con2 and Con3. Backbone enabling propagation in *E. coli* | This study |
| pYTKL3R4DO | Pre-assembled YTK acceptor plasmid into which single gene cassettes can be cloned, consisting of an sfGFP dropout part flanked by YTK connectors Con3 and Con4. Backbone enabling propagation in *E. coli* | This study |
| pYTKL4REDO | Pre-assembled YTK acceptor plasmid into which single gene cassettes can be cloned, consisting of an sfGFP dropout part flanked by YTK connectors Con4 and ConE. Backbone enabling propagation in *E. coli* | This study |
| pWO68 | YTK single gene cassette plasmid for constitutive mScarlet-I expression, pCCW12-mScarlet-I-tTDH1. Backbone enabling propagation in *E. coli* and integration at the HIS3 locus in *S. cerevisiae* | This study |
| pWS1766 | YTK position 3 part encoding the mScarlet-I protein | This study |
| pWS065 | Pre-assembled YTK acceptor plasmid into which single gene cassettes can be cloned, consisting of an sfGFP dropout part flanked by YTK connectors ConS and ConE. Backbone enabling propagation in *E. coli* | This study |

**Supplementary Table 3. Composition of YTK single gene constructs used in this study**

| | Parts | | | | | |
|---|---|---|---|---|---|---|
| **Plasmid** | **2** | **3a** | **3b** | **4a** | **4b** | **Backbone** |
| pWS702 | pYTK009 | pWS033 | | pYTK056 | | pWS473 |
| pCG01 | pYTK009 | pWS433 | pWS034 | pYTK056 | | pYTK096 |
| pCG04 | pYTK009 | pWS433 | pWS032 | pYTK056 | | pYTK096 |
| pCG05 | pYTK009 | pWS433 | pWS032 | pPPK027 | pYTK066 | pYTK096 |
| pCG16 | pYTK009 | pPPK041 | pPPK044 | pPPK027 | pYTK066 | pYTK096 |
| pCG17 | pYTK009 | pWS433 | pPPK044 | pPPK027 | pYTK066 | pYTK096 |
| pCG18 | pYTK009 | pPPK042 | pPPK045 | pPPK027 | pYTK066 | pYTK096 |
| pCG19 | pYTK009 | pWS433 | pPPK045 | pPPK027 | pYTK066 | pYTK096 |
| pCG20 | pYTK009 | pPPK043 | pPPK046 | pPPK027 | pYTK066 | pYTK096 |
| pCG21 | pYTK009 | pWS433 | pPPK046 | pPPK027 | pYTK066 | pYTK096 |
| pCG22 | pWS1078 | pPPK042 | pPPK045 | pPPK027 | pYTK066 | pWS042 |
| pGPY74 | pYTK027 | pWS930 | pWS935 | pYTK056 | | pWS041 |
| pGPY85 | pWS1078 | pWS033 | | pYTK056 | | pWS042 |
| pCBH1 | pYTK009 | pYTK-CBH1 | | pYTK056 | | pYTK096 |
| pCBH2 | pYTK009 | pYTK-CBH2 | | pYTK056 | | pYTK096 |
| pBGL1 | pYTK009 | pYTK-BGL1 | | pYTK056 | | pYTK096 |
| pEGL2 | pYTK009 | pYTK-EGL2 | | pYTK056 | | pYTK096 |
| pLPMO | pYTK009 | pYTK-LPMO | | pYTK056 | | pYTK096 |
| pXTH3 | pYTK009 | pYTK-XTH3 | | pYTK056 | | pYTK096 |
| pWO68 | pYTK010 | pWS1766 | | pYTK056 | | pWS065 |

**Supplementary Table 4. Composition of YTK multi gene constructs used in this study**

| Plasmid | Cassette 1 | Cassette 2 | Cassette 3 | Cassette 4 | Cassette 5 | Backbone |
|---|---|---|---|---|---|---|
| pCG23 | pGPY074 | pCG22 | NA | NA | NA | pYTK096 |
| pGPY093 | pGPY074 | pGPY085 | NA | NA | NA | pYTK096 |
| pCelMix | pLSR1-CBH1 | pL1R2-CBH2 | pL2R3-BGL1 | pL3R4-EGL2 | pL4RE-LPMO | pYTK096 |
| pTT-GFP | pLSR1-T-LexABD-CRY2 | pL1R2-spacer | pL2R3-LexA-GFP | pL3R4-spacer | pL4RE-T-VP16-CIB1 | pYTK096 |
| pRR-GFP | pLSR1-R-LexABD-CRY2 | pL1R2-spacer | pL2R3-LexA-GFP | pL3R4-spacer | pL4RE-R-VP16-CIB1 | pYTK096 |
| pTR-GFP | pLSR1-T-LexABD-CRY2 | pL1R2-spacer | pL2R3-LexA-GFP | pL3R4-spacer | pL4RE-R-VP16-CIB1 | pYTK096 |
| pRT-GFP | pLSR1-R-LexABD-CRY2 | pL1R2-spacer | pL2R3-LexA-GFP | pL3R4-spacer | pL4RE-T-VP16-CIB1 | pYTK096 |
| pNCellulose | pLSR1-R-LexABD-CRY2 | pL1R2-spacer | pL2R3-LexA-Nluc-CBM3 | pL3R4-spacer | pL4RE-T-VP16-CIB1 | pYTK096 |
| pNSurface | pLSR1-R-LexABD-CRY2 | pL1R2-spacer | pL2R3-LexA-Nluc-SED1 | pL3R4-spacer | pL4RE-T-VP16-CIB1 | pYTK096 |

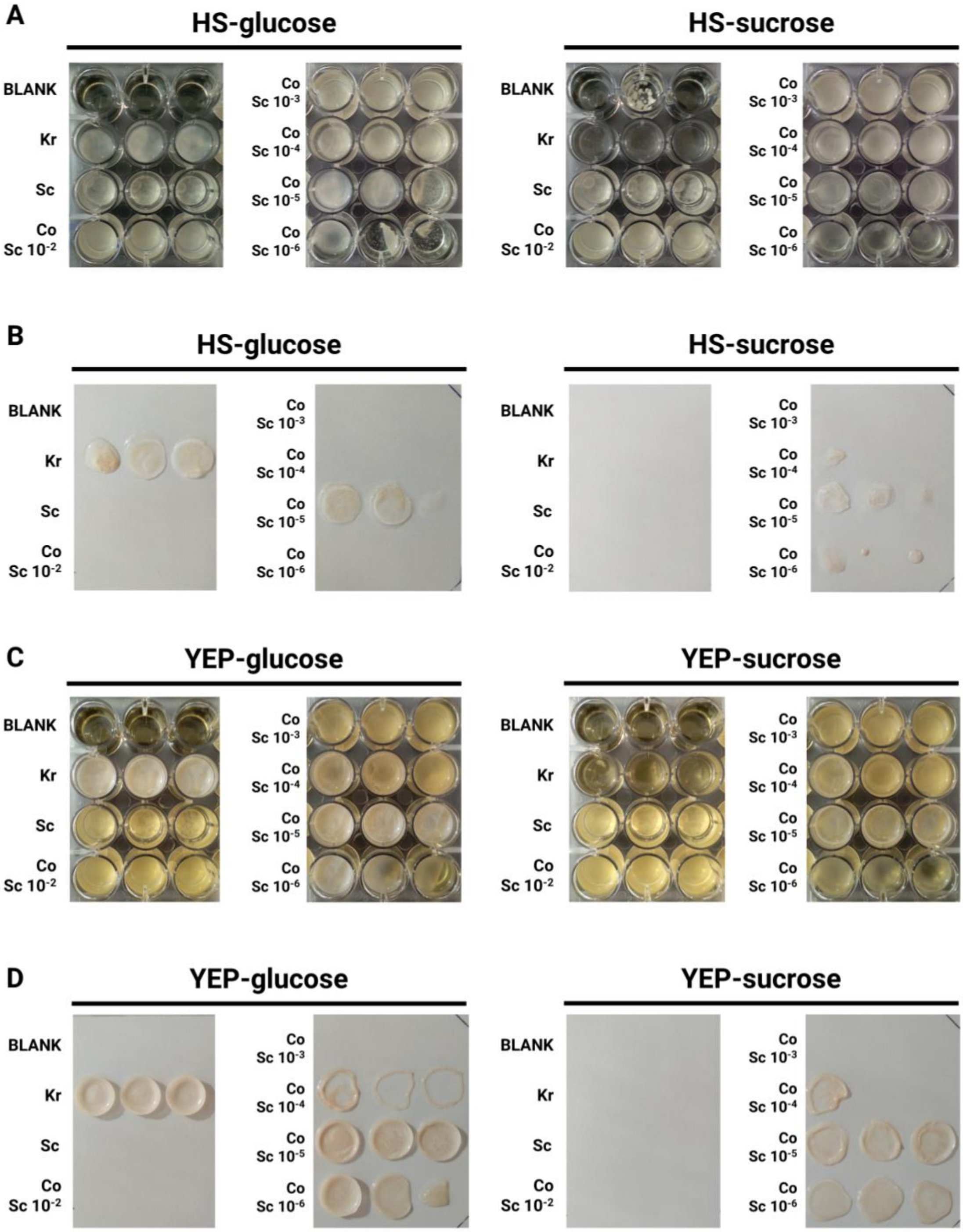


**Supplementary Figure 1. Images of cultures and pellicles from the co-culture condition screen.** *S. cerevisiae* (Sc) and *K. rhaeticus* (Kr) were inoculated in mono-culture or co-culture (Co) in rich yeast media (YEP) or BC-producing bacteria media (HS) with either glucose or sucrose as the carbon source. For co-cultures, the Sc pre-cultures were diluted into fresh medium over a range from 1/100 (Sc $10^{-2}$) to 1/$10^{6}$ (Sc $10^{-6}$). In mono-culture, Sc pre-cultures was diluted 1/100 and Kr pre-cultures was diluted 1/50. As a control for contamination, wells were included in which no cells were inoculated (BLANK). After 4 days incubation at 30°C, images were taken of cultures and then of isolated pellicle layers, where present. Cultures (**A**) and pellicles (**B**) produced in HS media and cultures (**C**) and pellicles (**D**) produced in YEP media.

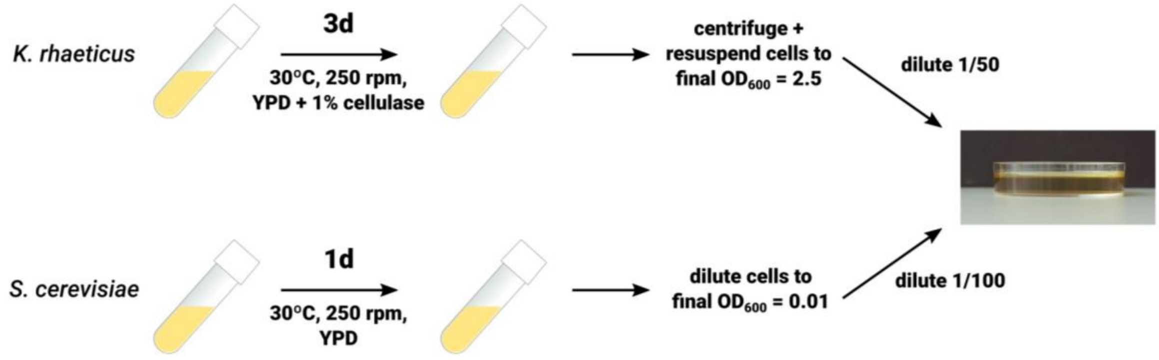


**Supplementary Figure 2. Defining and testing a standard protocol for co-culturing *S. cerevisiae* and *K. rhaeticus*.** Schematic outlining the standard co-culture protocol. *K. rhaeticus* and *S. cerevisiae* are grown in mono-culture under agitation. *K. rhaeticus* cultures are then centrifuged and resuspended in YEP-sucrose (YPS) medium to a final $OD_{600}$ = 2.5 – this step removes trace amounts of cellulase enzyme and normalises cell density. *S. cerevisiae* cultures are normalised by diluting to an $OD_{600}$ = 0.01 in YPS. Normalised *K. rhaeticus* and *S. cerevisiae* cultures are then inoculated into fresh YPS by diluting 1/50 and 1/100, respectively

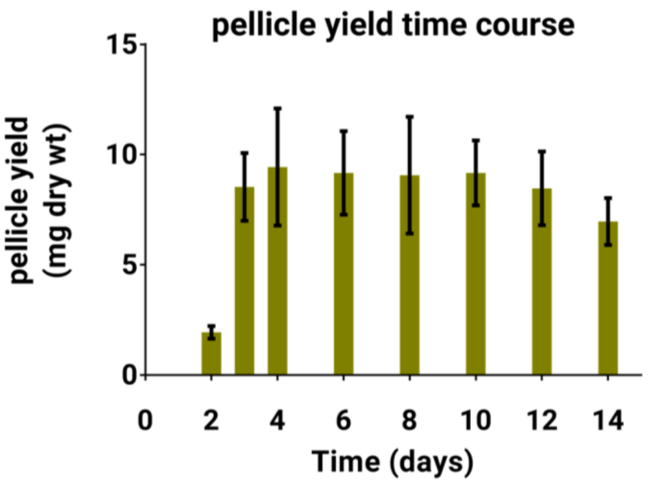


**Supplementary Figure 3. Measuring co-culture pellicle yields.** To follow BC production dynamics over time, co-cultures were prepared following our standard protocol and left incubating over several days. At each time point, pellicle layers were removed and dried. Once dried, pellicles were weighed to determine the pellicle yield. Notably, since pellicles were not treated to lyse and remove cells, this measurement includes contribution from both BC yield and entrapped cells. Pellicle yield rapidly increased between 2 and 3 days, at which point it plateaus. Samples prepared in triplicate, data represent the mean ±1 SD.

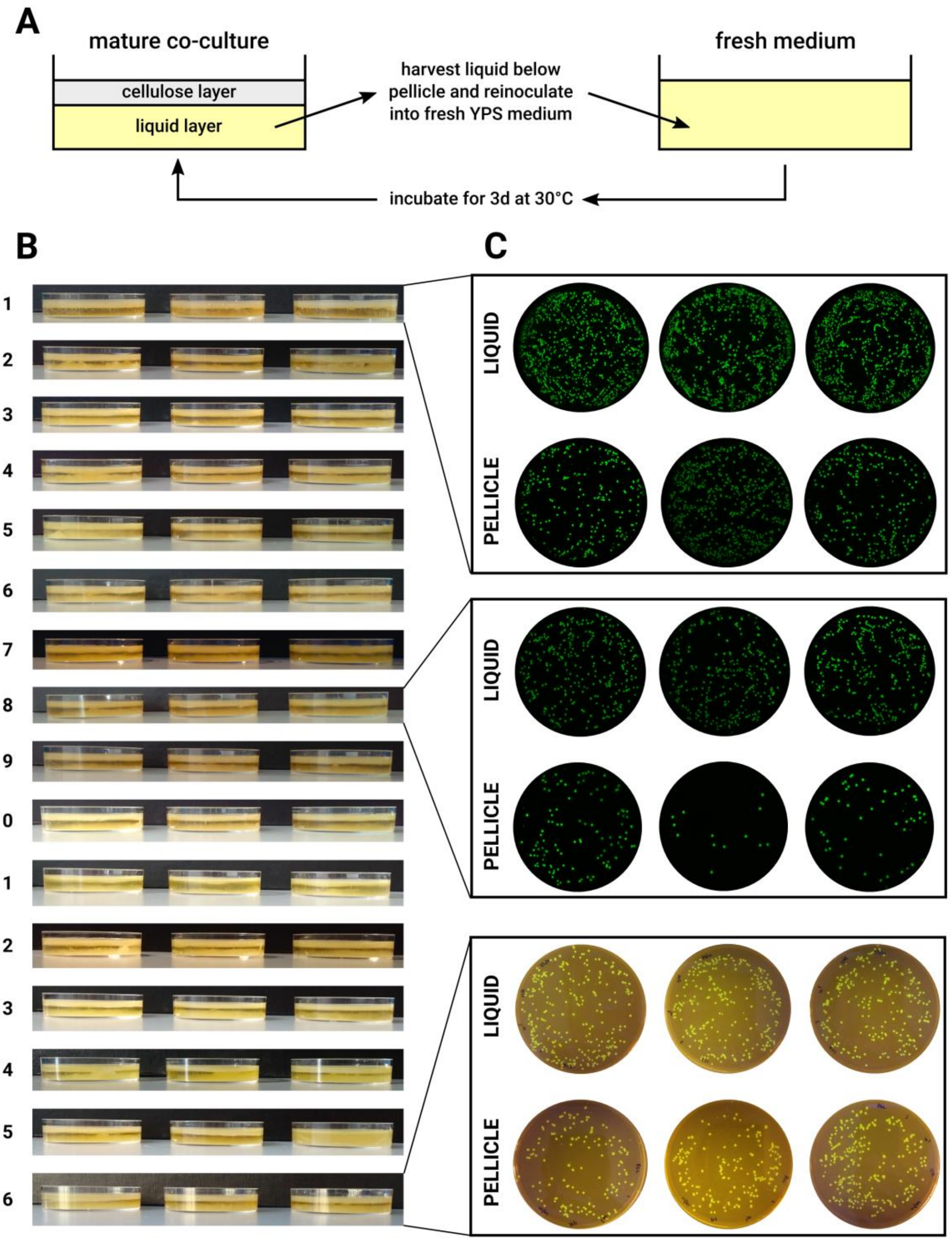


**Supplementary Figure 4. Investigating co-culture stability by passage. (a)** Co-cultures of *S. cerevisiae* yWS167 and *K. rhaeticus* Kr RFP were passaged by iteratively back-diluting liquid from below the pellicle layer in mature co-cultures into fresh YPS medium. **(b)** At each stage, mature pellicles were imaged. Pellicle formation was constant, indicating *K. rhaeticus* was growing well. In addition, a clear sediment was formed below the pellicle, consistent with *S. cerevisiae* growth. **(c)** To confirm the presence of the initial *S. cerevisiae* strain, which expresses GFP, in passage co-cultures, samples of the liquid below the pellicle (LIQUID) and enzymatically-degraded pellicles (PELLICLE) were plated and imaged for GFP fluorescence. In the interest of clarity, plates from only three time points are shown here. The appearance of the final time point is different as it was imaged for fluorescence using different equipment (fluorescence scanner versus imaging under a transilluminator). All images show that the initial GFP-expressing *S. cerevisiae* strain was maintained throughout passage.

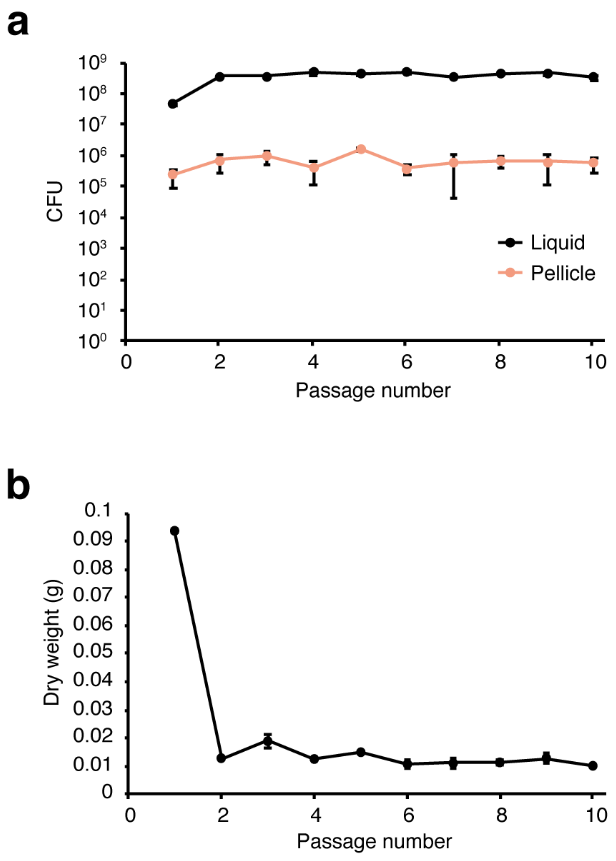


**Supplementary Figure 5. Yeast cell count and pellicle dry weight across 10 passages. (a)** Yeast colony forming unit (CFU) of pellicle from the first passage to the tenth passage. Each passage was inoculated using liquid from the previous passage. Data represent the mean ±1 SD from biological triplicates. **(b)** Pellicle dry weight from the first passage to the tenth passage. Pellicles were free-dried using a lyophilizer. Data represent the mean ±1 SD from biological triplicates.

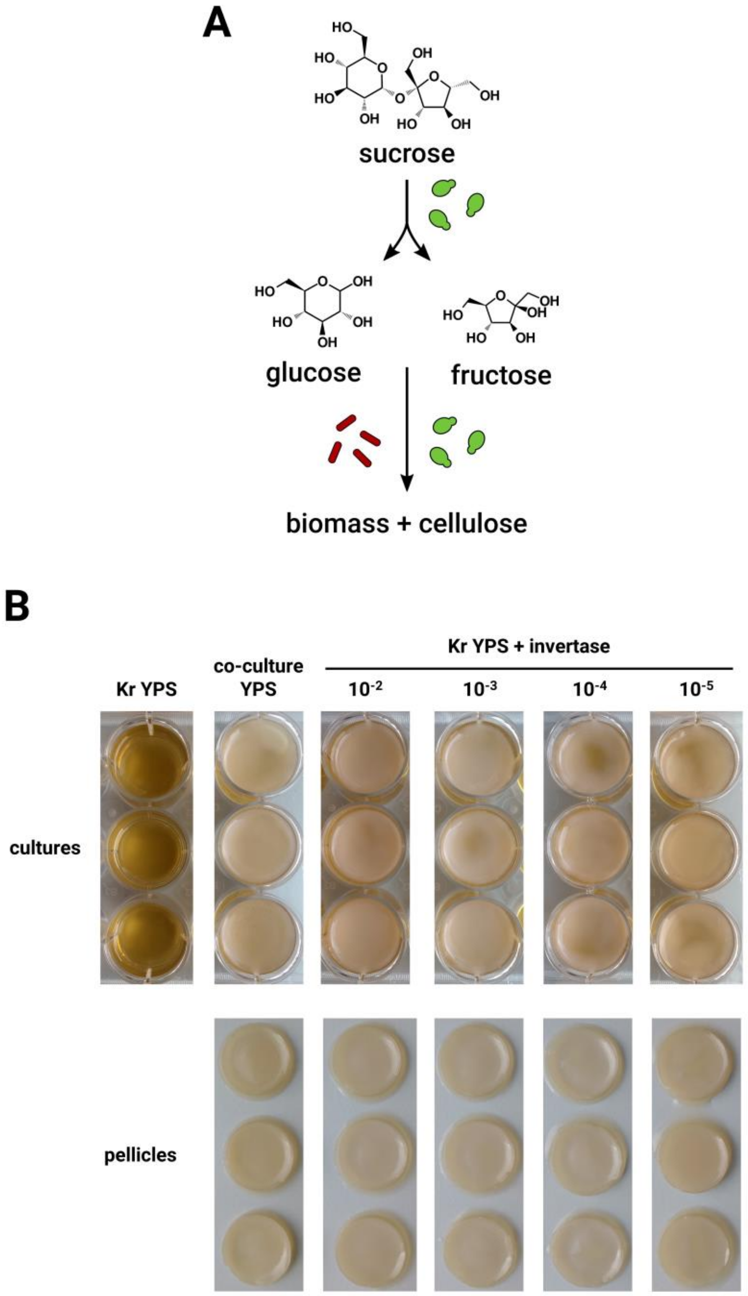


**Supplementary Figure 6. A putative metabolic mechanism for *S. cerevisiae* stimulation of *K. rhaeticus* growth. (a)** Various studies report that yeast (green) in kombucha microbial communities degrade extracellular sucrose to glucose and fructose which both yeast and BC-producing bacteria (red) consume to produce biomass. **(b)** A variety of cultures were prepared in YPS: *K. rhaeticus* Kr RFP mono-culture (Kr YPS), co-cultures of Kr RFP and *S. cerevisiae* yWS167 (co-culture YPS) and mono-cultures of *K. rhaeticus* Kr RFP spiked with a range of dilutions of a stock solution of commercial *S. cerevisiae* invertase at 5000 U/mL concentration (1/100, 1/1000. 1/10,000 and 1/100,000). Images were taken of cultures and, where present, isolated pellicles after 3 days of incubation at 30°C.

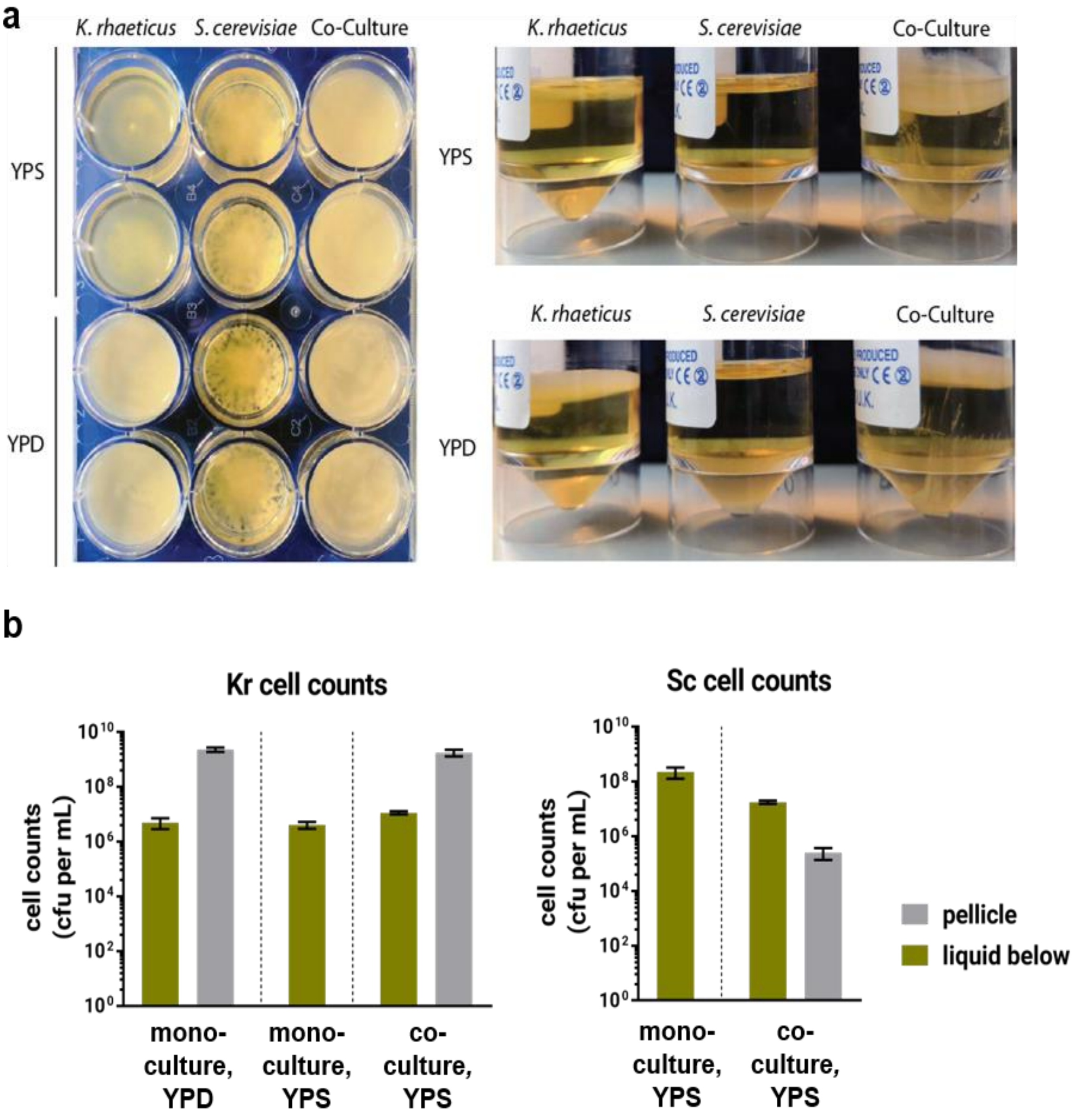


**Supplementary Figure 7. Growing co- and mono cultures of engineerable *S. cerevisiae* and *K. rhaeticus* in YPP and YPS media. (a)** Images of mono-cultures and co-cultures of *K. rhaeticus* and *S. cerevisiae* grown for 3 days. *S. cerevisiae* grows well in both YPD and YPS media, forming a sediment at the base of the culture. *K. rhaeticus* grew well in YPD medium, forming a thick pellicle layer at the air-water interface, but failed to form a pellicle in YPS medium. When co-cultured, in both YPD and YPS, a thick pellicle layer was formed as well as a sediment layer at the base of the culture, indicating both *S. cerevisiae* and *K. rhaeticus* had grown. The left panel shows a top view from the different cultures in a 24-well plate. The right panel shows a side view from the different cultures incubated in 20 ml reaction tubes. **(b)** Cell counts of *K. rhaeticus* Kr RFP and *S. cerevisiae* yWS167 were determined by plating and counting the numbers of cells present in the two phases of co-cultures – the liquid layer and the pellicle layer. Cell counts were determined for *K. rhaeticus* grown in mono-culture in YPD (Kr YPD) or YPS (Kr YPS) or in co-culture with *S. cerevisiae* in YPS (Co YPS). Cell counts were determined for S. cerevisiae grown in mono-culture in YPS (Sc YPS) or in co-culture with K. rhaeticus in YPS (Co YPS). Samples prepared in triplicate, data represent the mean ±1 SD.

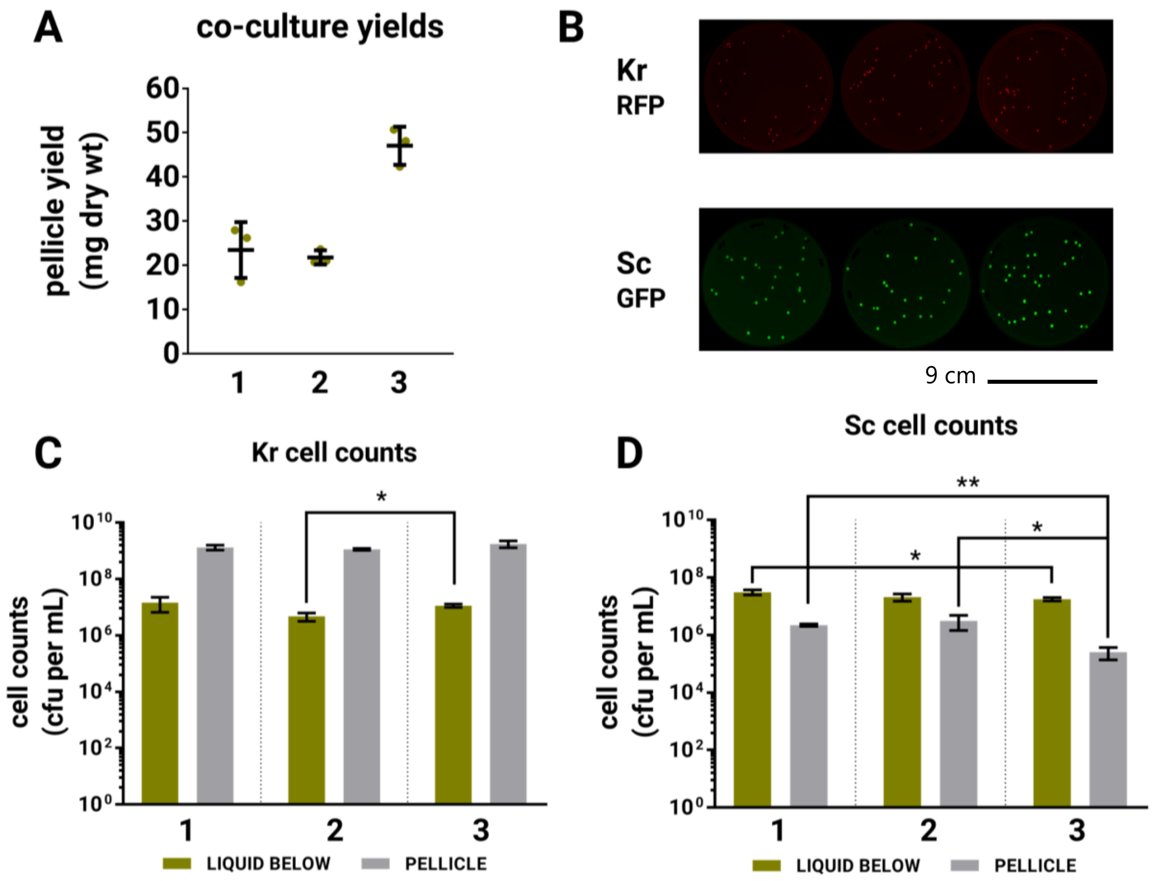


**E**

**Kr cell counts**

| | liquid | | | pellicle | | |
|---|---|---|---|---|---|---|
| | 1 | 2 | 3 | 1 | 2 | 3 |
| mean cell counts (cfu/mL) | 1.47E+07 | 4.75E+06 | 1.14E+07 | 1.33E+09 | 1.14E+09 | 1.77E+09 |
| standard deviation (cfu/mL) | 8.02E+06 | 1.53E+06 | 1.46E+06 | 2.60E+08 | 8.43E+07 | 4.82E+08 |

**F**

**Sc cell counts**

| | liquid | | | pellicle | | |
|---|---|---|---|---|---|---|
| | 1 | 2 | 3 | 1 | 2 | 3 |
| mean cell counts (cfu/mL) | 3.10E+07 | 2.09E+07 | 1.78E+07 | 2.23E+06 | 3.16E+06 | 2.53E+05 |
| standard deviation (cfu/mL) | 6.24E+06 | 5.80E+06 | 2.42E+06 | 2.08E+05 | 1.70E+06 | 1.15E+05 |

**Supplementary Figure 8. Reproducibility of co-culture pellicle yields and cell densities. (a)** Pellicle yields were measured on three separate occasions. For each repeat samples were prepared in triplicate, horizontal bars represent the mean ±1 SD, green circles represent the values of individual samples. **(b)** Cell counts from co-cultures were prepared on three separate occasions by plating onto selective media and scanning for RFP fluorescence for *K. rhaeticus* and GFP fluorescence for *S. cerevisiae*. Cell counts were recorded from both the liquid and pellicle layers for both *K. rhaeticus* **(c)** and *S. cerevisiae* **(d)**. Data here come from one of the three separate experiments. Samples were prepared in triplicate, p-values calculated by unpaired, two-tailed t-test data (* $p < 0.05$ and ** $p < 0.005$), data represent the mean ±1 SD. Since logarithmic scales can mask some of the variation, numerical values of cell counts are included for both *K. rhaeticus* **(e)** and *S. cerevisiae* (**f**).

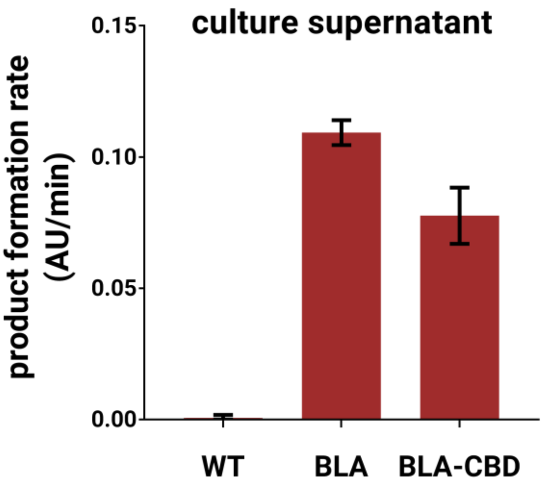


**Supplementary Figure 9. Secreted β-lactamase activity in *S. cerevisiae* mono-culture.** Culture supernatants from WT, BLA and BLA-CBD strains were assayed for β-lactamase activity using the colourimetric nitrocefin substrate. The product formation rate was measured using a plate reader. Samples prepared in triplicate, data represent the mean ±1 SD. Notably, the activity detected from the BLA-CBD secreting strain, yCG05, was reduced compared to the BLA secreting strain, yCG04. Since this assay was performed using undiluted supernatants from 24h cultures, the observed activity will be affected by multiple factors. Therefore, the decrease in β-lactamase activity for BLA-CBD could be due to decreased growth rate, decreased secreted protein yields or an effect of fusion of the CBD to BLA enzyme decreasing its activity by causing steric hindrance, for example.

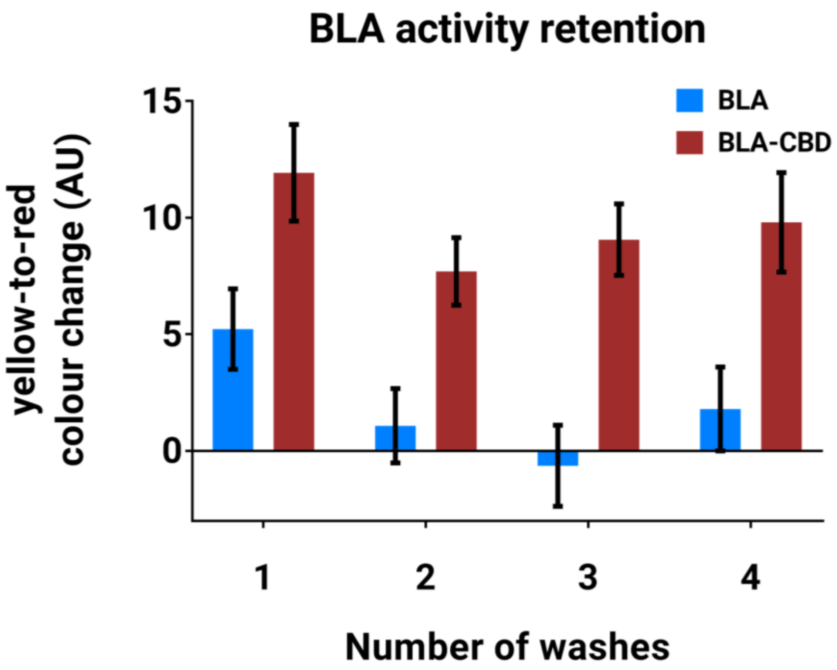


**Supplementary Figure 10. Retention of β-lactamase within functionalised material after washing.** As the BLA enzyme is passively incorporated within the BC matrix by diffusion and the BLA-CBD fusion is specifically bound through the CBD-cellulose interaction, it might, be anticipated that BLA enzyme could leach out of the BC material over time, while BLA-CBD would remain bound stably. To test this, dried pellicles functionalised with BLA and BLA-CBD were subjected to multiple rounds of washes in PBS buffer and then assayed for β-lactamase activity. The activity of β-lactamase in BLA-functionalised pellicles (by nitrofecin assay) fell sharply after washing, by contrast, BLA-CBD-functionalised pellicles retained a greater proportion of their original β-lactamase activity after washing. Samples prepared in triplicate, data represent the mean ±1 SD.

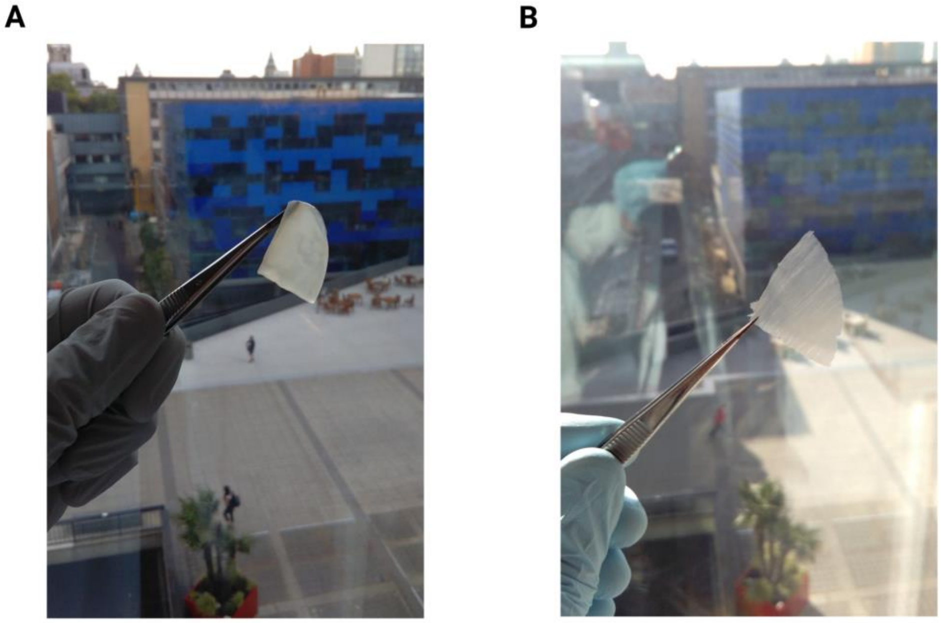


**Supplementary Figure 11. Images of wet and dried BC pellicles. (a)** A piece of BC pellicle in its native, wet stated. **(b)** A piece of a BC pellicle following drying using the sandwich method. Dried pellicles are much thinner than wet pellicles due to water loss and are similar in appearance to thin sheets of paper.

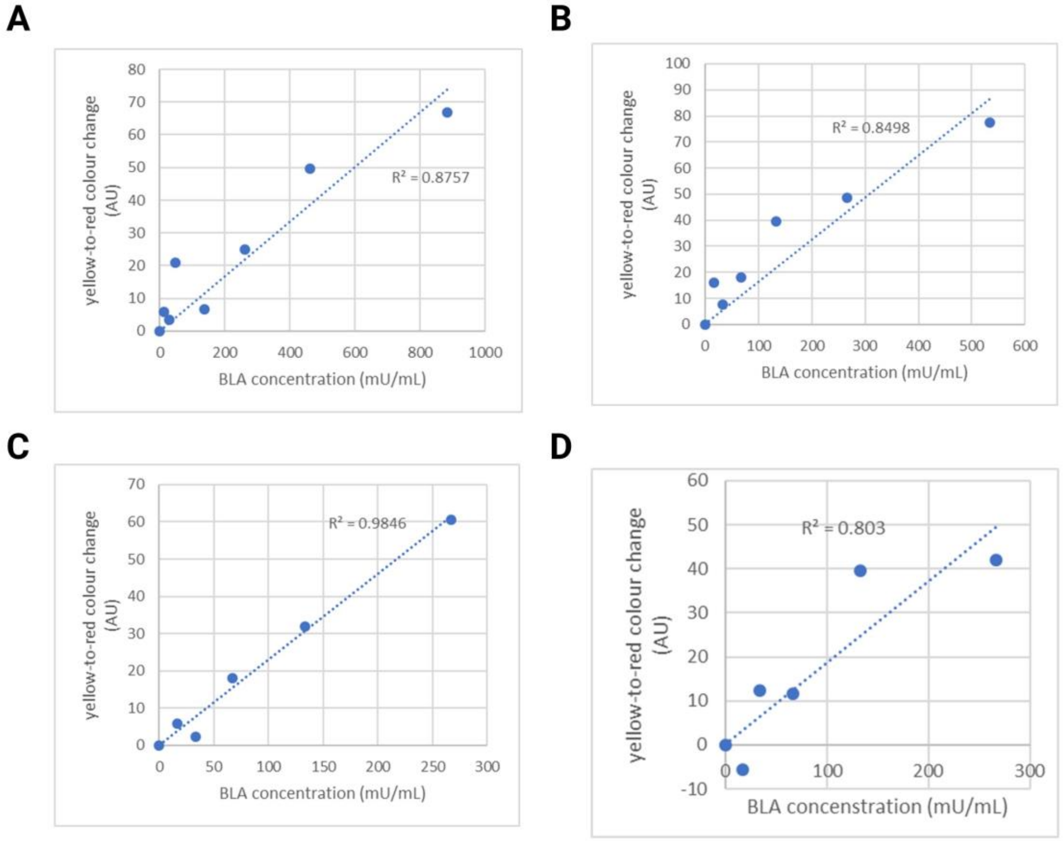


**Supplementary Figure 12. BLA assay standard curves.** To calculate absolute β-lactamase activities, standard curves were run alongside wet (**a**), dry 0 days (**b**), dry 1 month (**c**) and dry 6 month (**d**) samples. To prepare standard curves, pellicles from co-cultures prepared with WT yeast were treated exactly as sample pellicles (i.e. used wet, dried or dried and then stored). Pellicles were then supplemented with the indicated amounts of commercial E. coli β-lactamase enzyme to create known standards. Assays were performed in parallel with samples and images processed identically to generate standard curves as above.

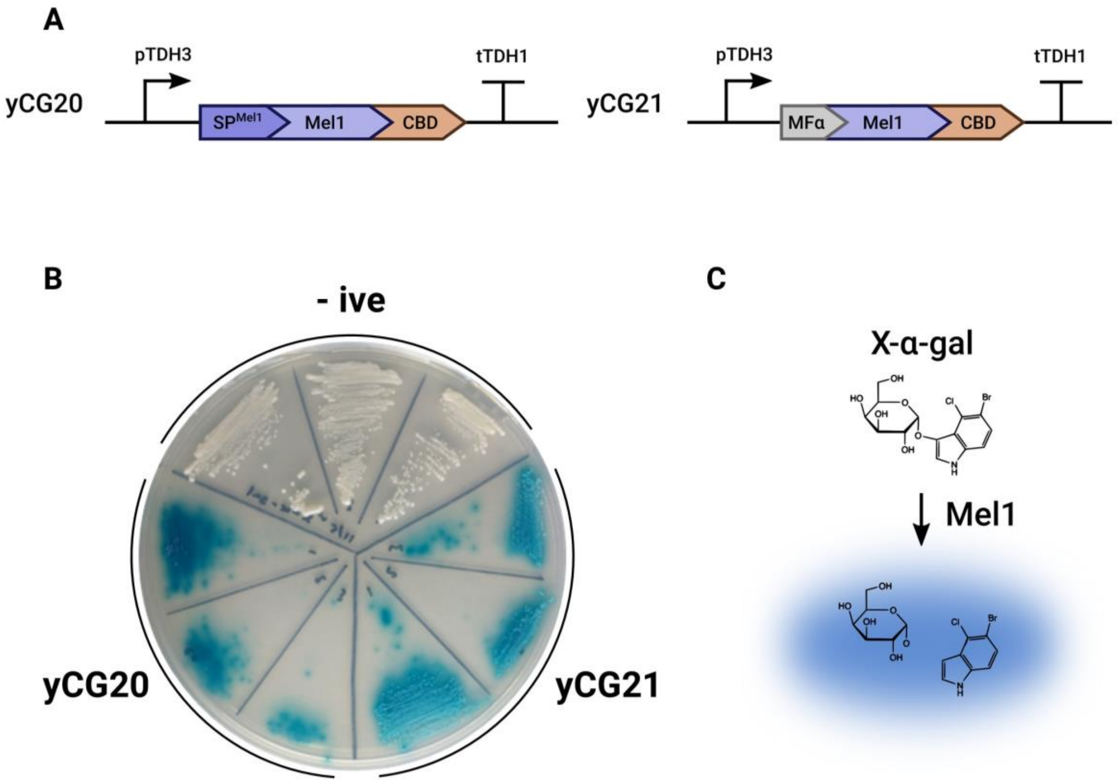


**Supplementary Figure 13. Secretion of the alpha-galactosidase Mel1. (a)** Two engineered strains were generated for Mel1 secretion. The first possessed the Mel1 N-terminal signal peptide and Mel1 catalytic region fused to CBDcex (yCG20). The second possessed the MF$\alpha$ signal peptide fused to the Mel1 catalytic region and CBDcex (yCG21). In both constructs, expression was driven by the strong constitutive promoter pTDH3. **(b)** Strains were screened for Mel1 secretion by a plate-based colourimetric assay. Transformants were re-streaked in triplicates on SC $URA^-$ agar supplemented with the colourimetric reporter X-$\alpha$-gal. After two days growth activity was detectable in the form of halos of blue pigment around colonies of both yCG20 and yCG21. No blue pigment was formed around colonies of the negative control strain, GFP-secreting yCG01 (-ive). Since the growth rate of yCG20 was severely reduced compared to that of yCG21 and yCG01, yCG21 was taken forwards for BC material functionalisation. **(c)** X-$\alpha$-gal is a colourimetric reporter for Mel1; in the presence of active $\alpha$-galactosidase enzymes, X-$\alpha$-gal is converted from a colourless substrate to a blue pigment.

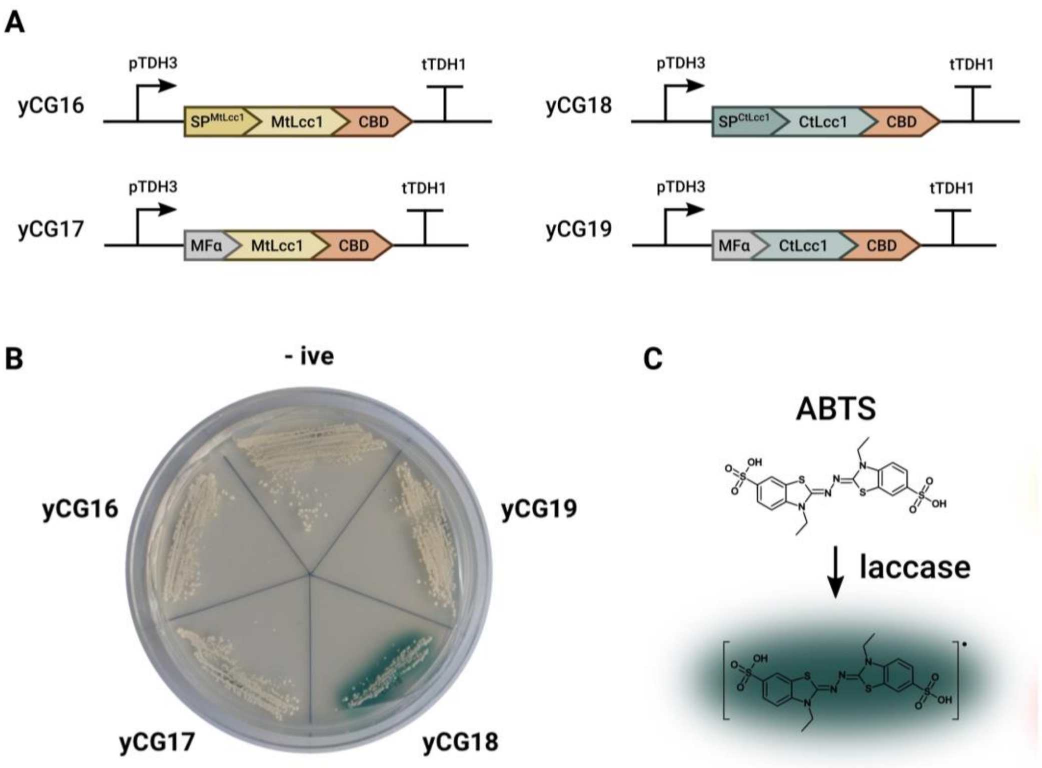


**Supplementary Figure 14. Secretion of fungal laccase enzymes. (a)** Four engineered strains were constructed for laccase enzyme secretion. Two strains were engineered to secrete a laccase from *Myceliophthora thermophila* (MtLcc1) with either the native signal peptide (yCG16) or the MF$\alpha$ signal peptide (yCG17). Two strains were engineered to secrete a laccase from *Coriolopsis trogii* (CtLcc1) with either the native signal peptide (yCG18) or the MF$\alpha$ signal peptide (yCG19). All constructs possessed a C-terminal CBD fusion and were expressed from the strong constitutive promoter pTDH3. **(b)** Strains were screened for laccase secretion by a plate-based colourimetric assay. Transformants were re-streaked on SC URA$^-$ agar supplemented with the colourimetric reporter 2,2'-azino-bis(3-ethylbenzothiazoline-6-sulphonic acid) (ABTS) and $CuSO_4$. After two days growth activity was detectable in the form of halos of green pigment around colonies of only yCG18. By contrast, no green pigment was formed around colonies of the negative control strain, GFP-secreting yCG01 (-ive) nor yCG16, yCG17 or yCG19 (although low level activity could be detected after longer incubation times). Therefore, yCG18 was taken forwards for BC material functionalisation. **(c)** ABTS is a colourimetric reporter for laccase activity; in the presence of active laccase enzyme, ABTS is converted from a colourless substrate to a green pigment.

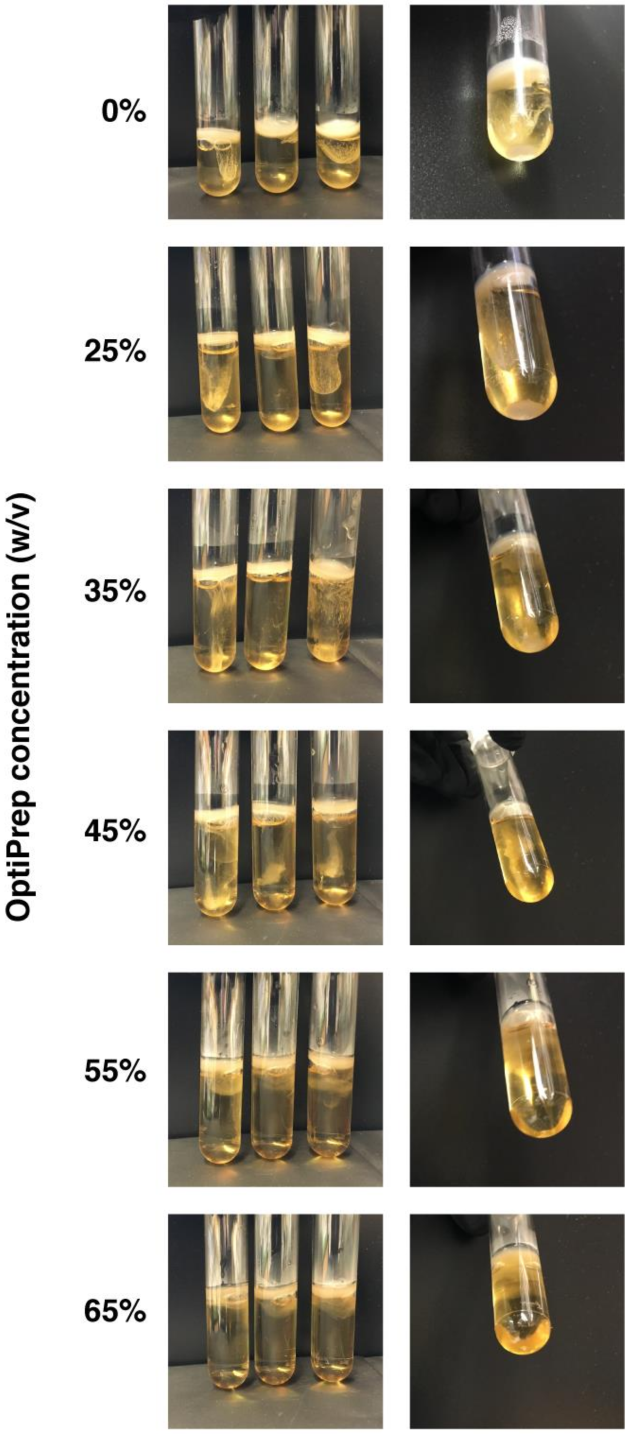


**Supplementary Figure 15. The effect of increasing concentrations of OptiPrep on the sedimentation of cells in co-cultures.** Co-cultures (Sc BY4741 and Kr) were prepared in YPS medium supplemented with the indicated concentrations of OptiPrep. After 3 days growth, samples were imaged in triplicate and the bottoms of individual tubes were imaged. The cell sediment is only apparent at OptiPrep concentrations less than 35%.

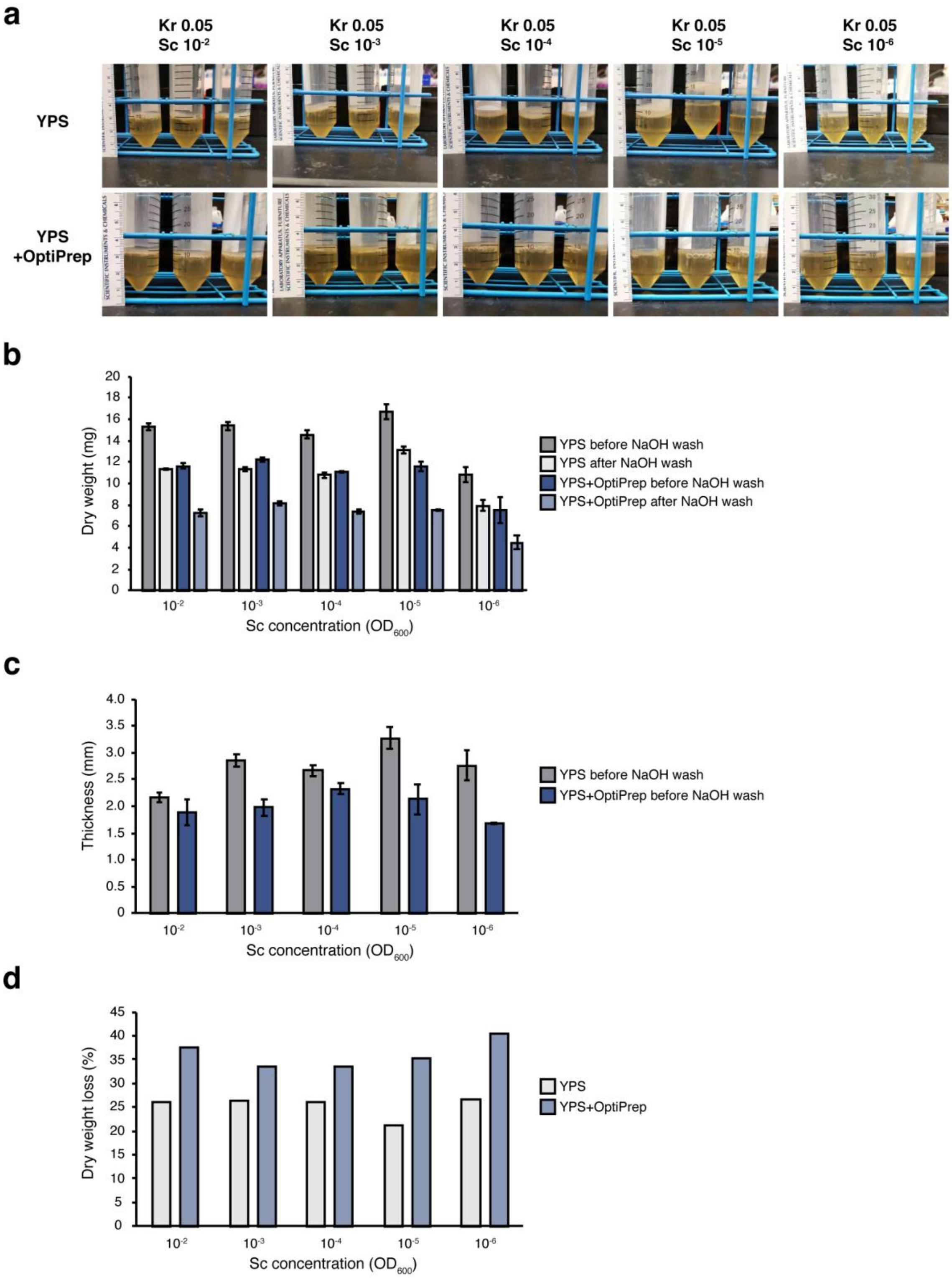


**Supplementary Figure 16. Comparison of co-culture behavior in YPS compared to YPS + OptiPrep media. (a)** Side view of pellicles grown for 3 days with different starting Sc concentrations (unit: $OD_{600}$). **(b)** Dry weights of pellicles grown in YPS and YPS+OptiPrep before and after NaOH wash. (c) Thickness of pellicles grown in YPS and YPS+OptiPrep. (d) Loss in dry weight (% of weight before NaOH wash) of pellicles grown in YPS and YPS+OptiPrep. The extra weight loss in YPS+OptiPrep might originate from the yeast cells incorporated into the bottom surface. Data represent the mean ±1 SD from triplicates.

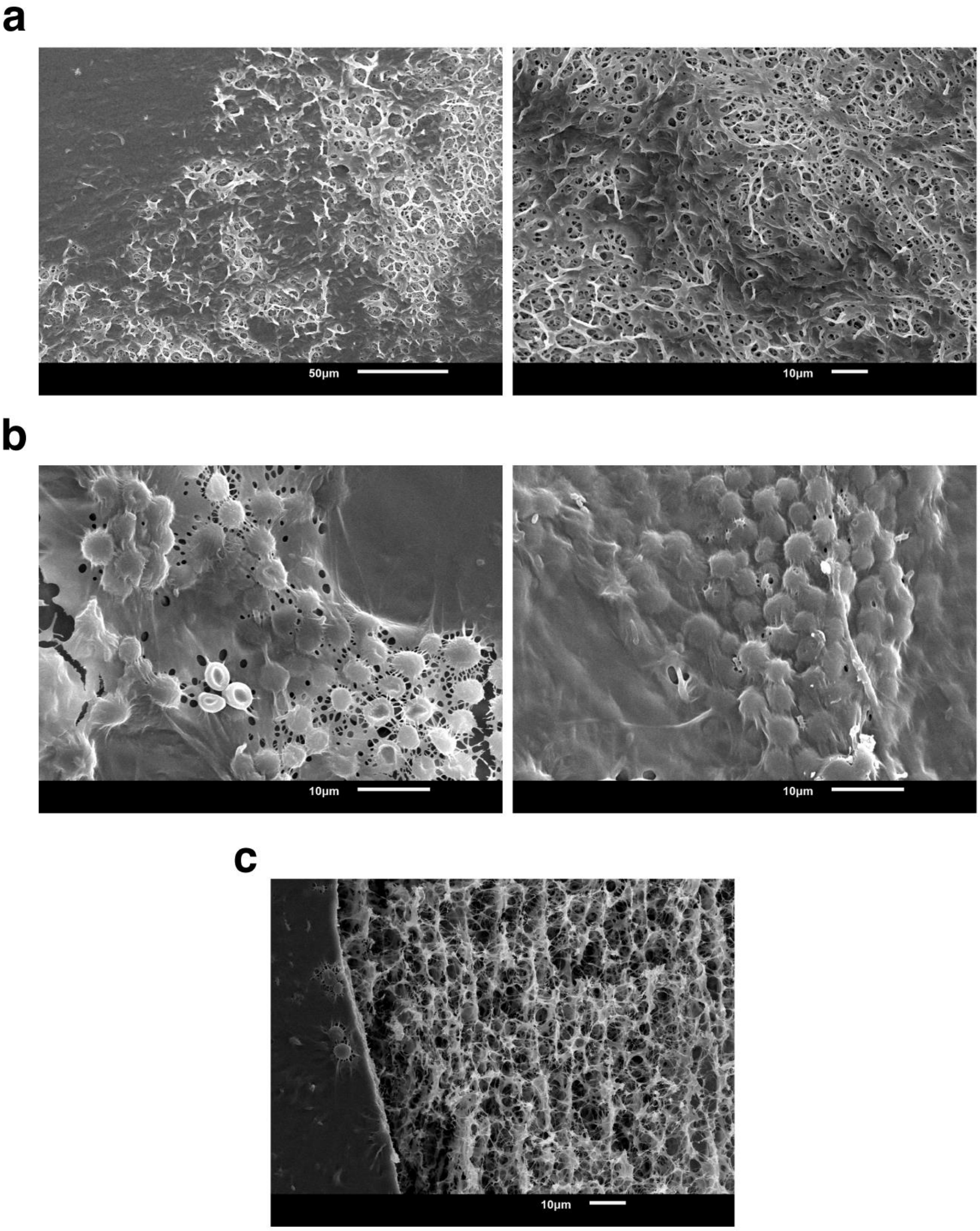


**Supplementary Figure 17. Additional SEM images of dried pellicles grown in YPS + OptiPrep. (a)** Top surface of the pellicle. **(b)** Bottom surface of the pellicle. Spherical yeast cells are trapped in the cellulose matrix produced by rod shape *K. rhaeticus*. **(c)** Cross section of the pellicle. Part of the bottom surface is presented on the left. Notably, the top surface of pellicles is largely evenly-covered with *K. rhaeticus* cells, while the lower surface contains numerous yeast cells localised in colony-like foci.

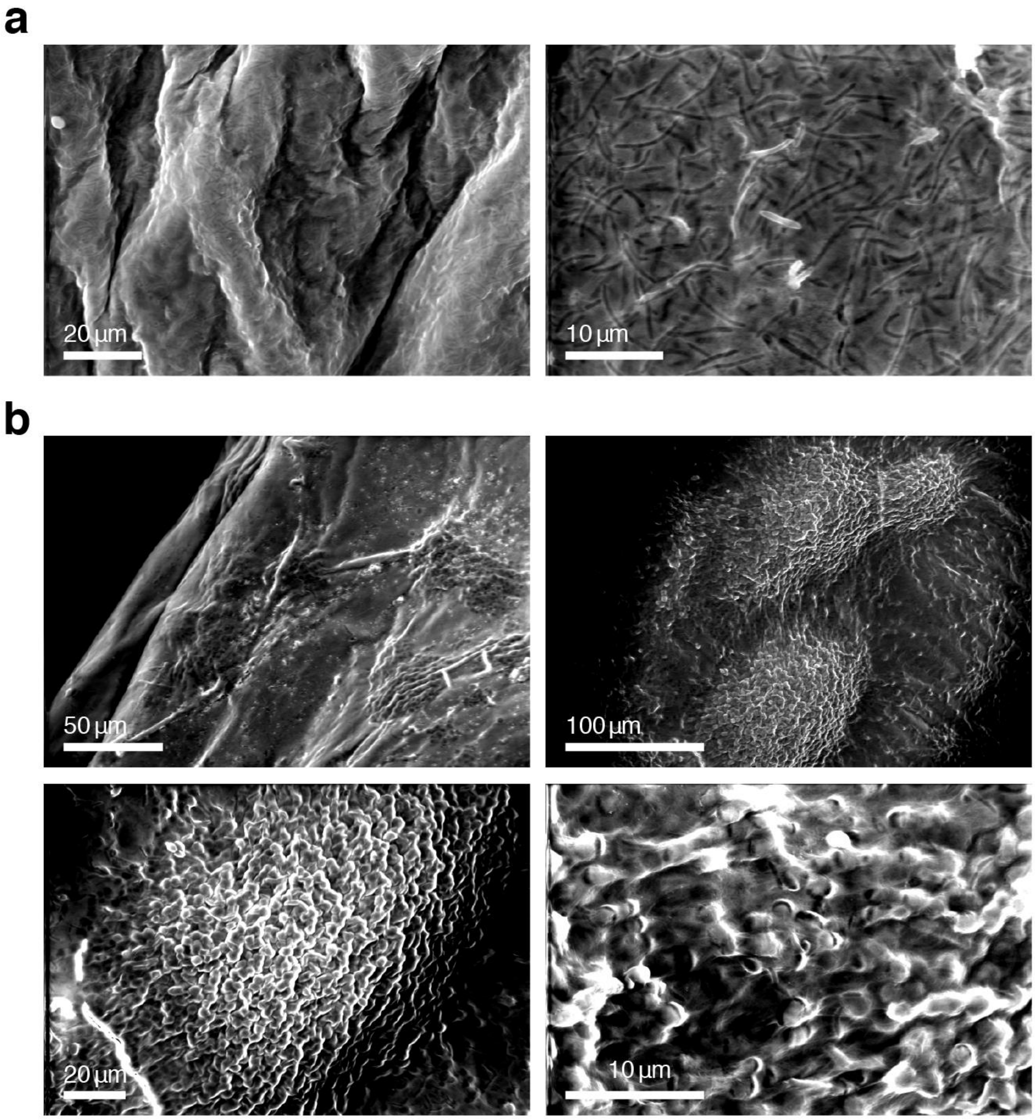


**Supplementary Figure 18. Environmental SEM (eSEM) images showing the native, hydrated structure of pellicles grown in YPS + OptiPrep. (a)** Top surface, where rod-shape *K. rhaeticus* cells covering the entire surface can be easily observed. **(b)** Bottom surface, where numerous yeast cells are localised in colony-like foci.

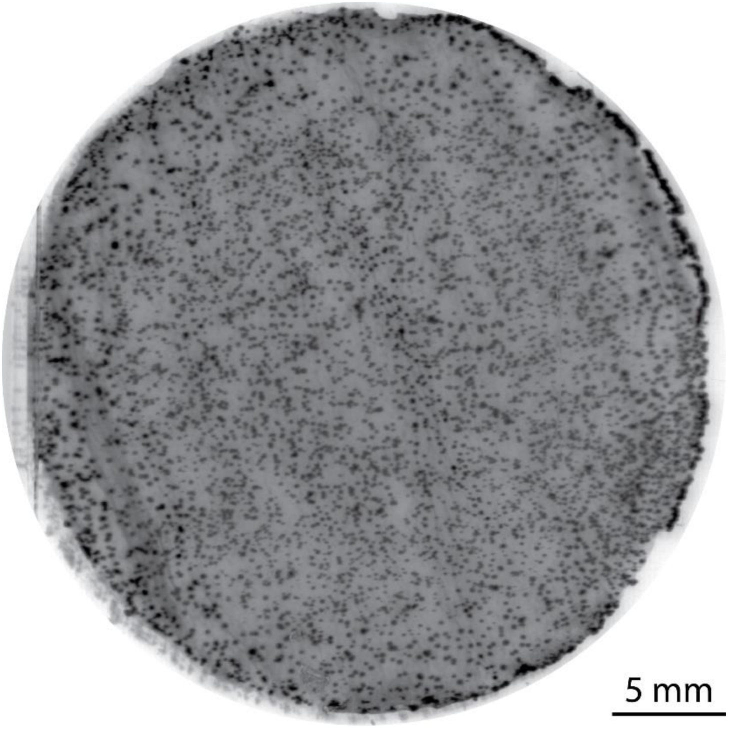


**Supplementary Figure 19. Fluorescence scan of a dried cellulose pellicle containing yeast expressing mScarlet-I.** After 3 days of growth of a static co-culture in YPS media supplemented with 40% (v/v) iodixanol the pellicle was removed from the culture and dried between absorbent papers. The pellicle was then imaged with a laser scanner to obtain a 2D distribution landscape of the incorporated yeast cells. The image was inverted to enhance the visibility of the fluorescent yeast colonies (in black). The darker the spots are the higher is the fluorescence. The pellicle has a diameter of ca. 30 mm as it was grown in a 50 ml falcon.

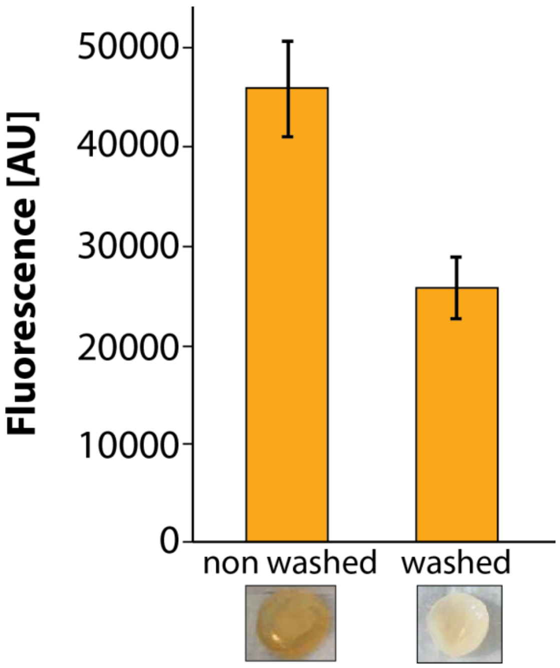


**Supplementary Figure 20. Fluorescence measurement of cell sediment after cellulase digestion of the harvested and dried BC pellicle grown in YPS+OptiPrep.** The co-culture was inoculated in 2 ml media with yeast (final OD700=0.0001) and *K. rhaeticu*s (final OD700=0.05) at the same time and grown statically at 30°C for three days in 24-well plates. Three pellicles (washed with 0.75 x PBS and not washed) were digested separately and the fluorescence of the freed, RFP-expressing yeast cells was determined. The fluorescence was measured with a Synergy HT (BioTek) from the bottom of the plate at 530 nm excitation and 590 nm emission with a gain of 55. The plate was shook for 30 seconds before measuring. Error bars represent the standard deviation of three different digested pellicles**.**

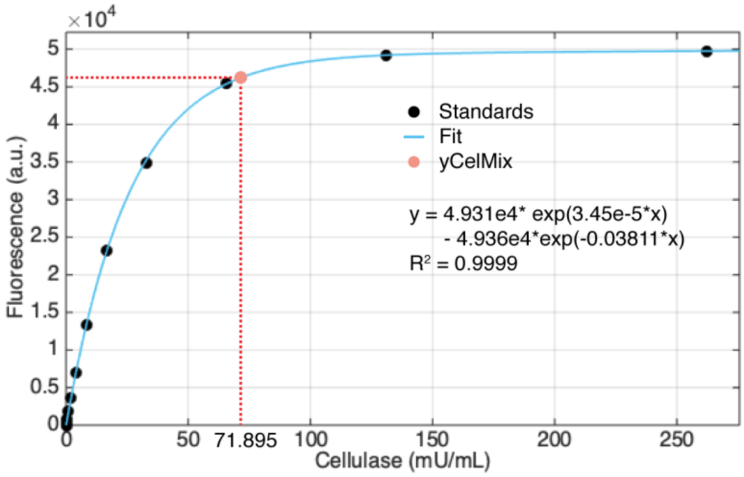


**Supplementary Figure 21. Total cellulase activity of yCelMix.** The total cellulase activity of yCelMix saturated culture was calculated from standards prepared with *T. reesei* cellulase mix. Data represent the mean from biological triplicates.

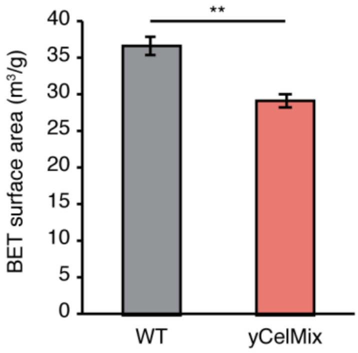


**Supplementary Figure 22. BET surface area is decreased in yCelMix pellicle.** Total BET surface area was calculated from dried pellicles. Data represent the mean ±1 SD from biological triplicates. ** $p < 0.01$.

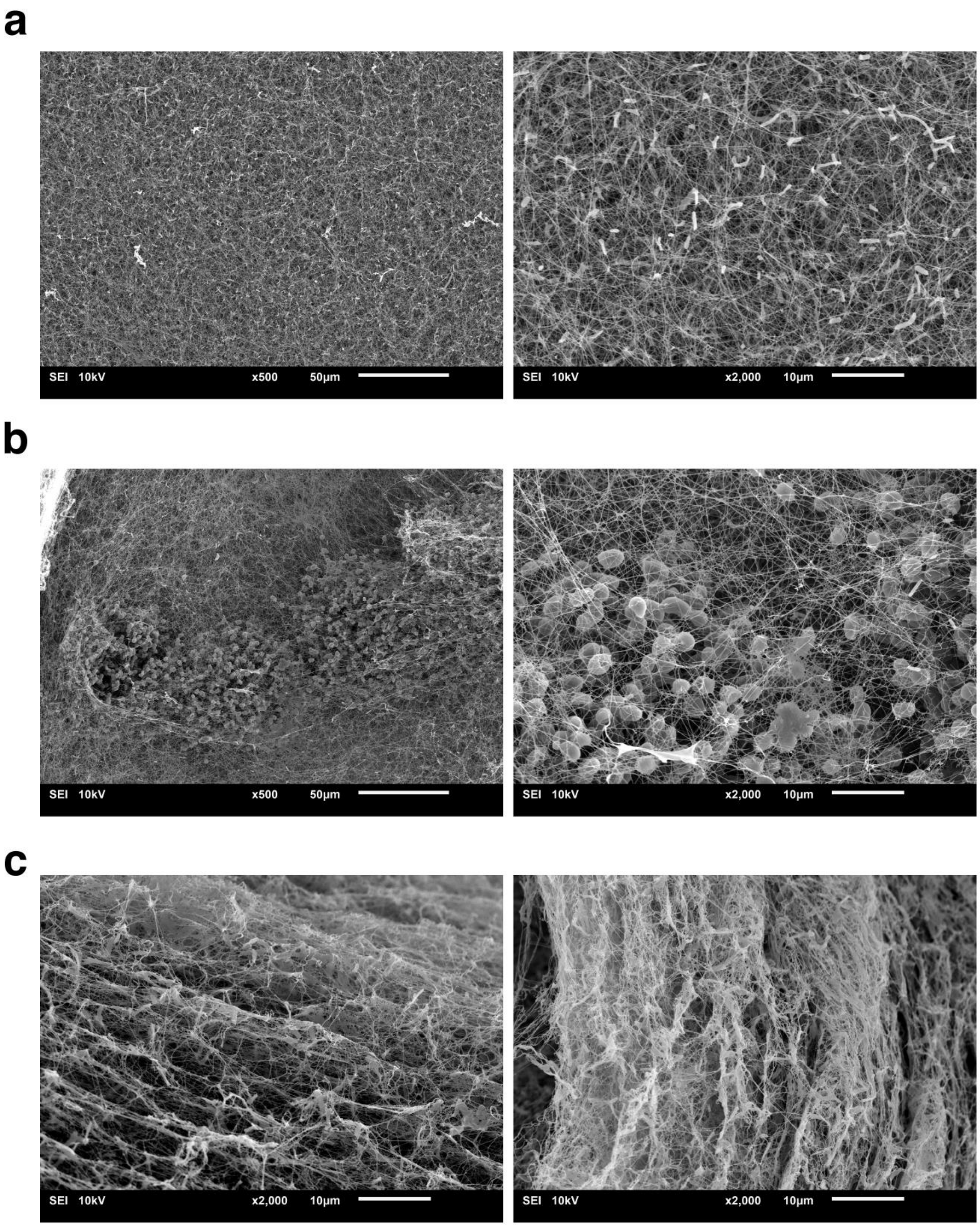


**Supplementary Figure 23. SEM images of dried yCelMix pellicles grown in YPS + OptiPrep. (a)** Top surface of the pellicle. **(b)** Bottom surface of the pellicle. Spherical yeast cells are trapped in the loose cellulose matrix, which is partially degraded by the cellulase cocktail. **(c)** Cross sections of the pellicle.

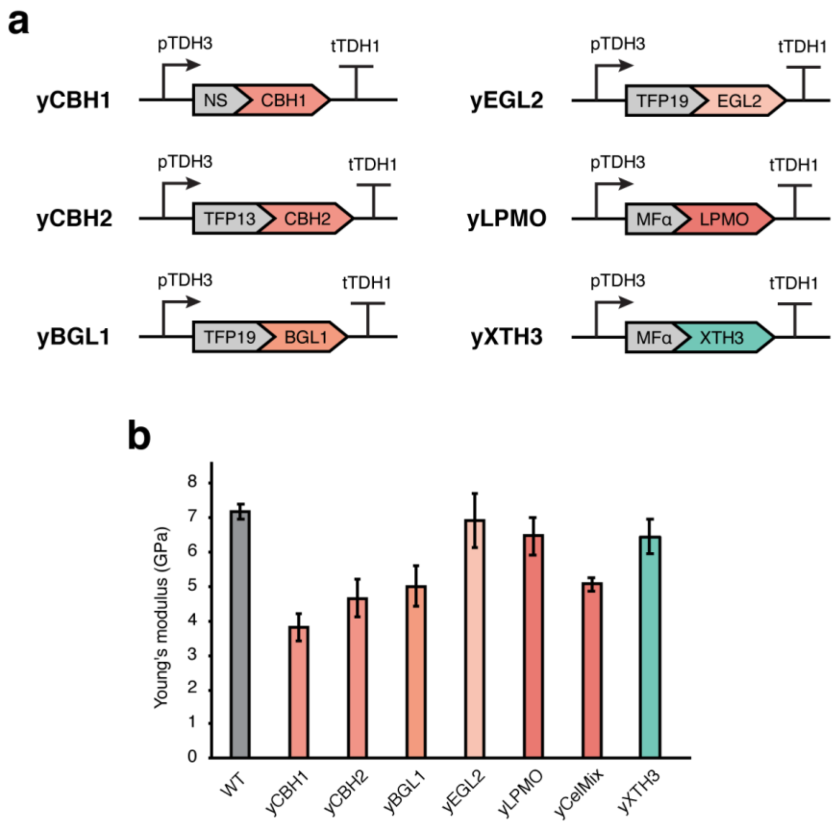


**Supplementary Figure 24. Single cellulase secretion strains and their effect on BC stiffness. (a)** Schematic illustrating the construction of single cellulase secretion strains. XTH3 is an *Arabidopsis thaliana* cell-wall enzyme which catalyses covalent cross-linking between cellulose[10]. Expression of individually secretion-tagged cellulases is driven by the strong constitutive promoter pTDH3. **(b)** Young's moduli of dried pellicles from single cellulase secretion strains and from a cross-linking enzyme secretion strain. Data represent the mean ±1 SD from biological triplicates.

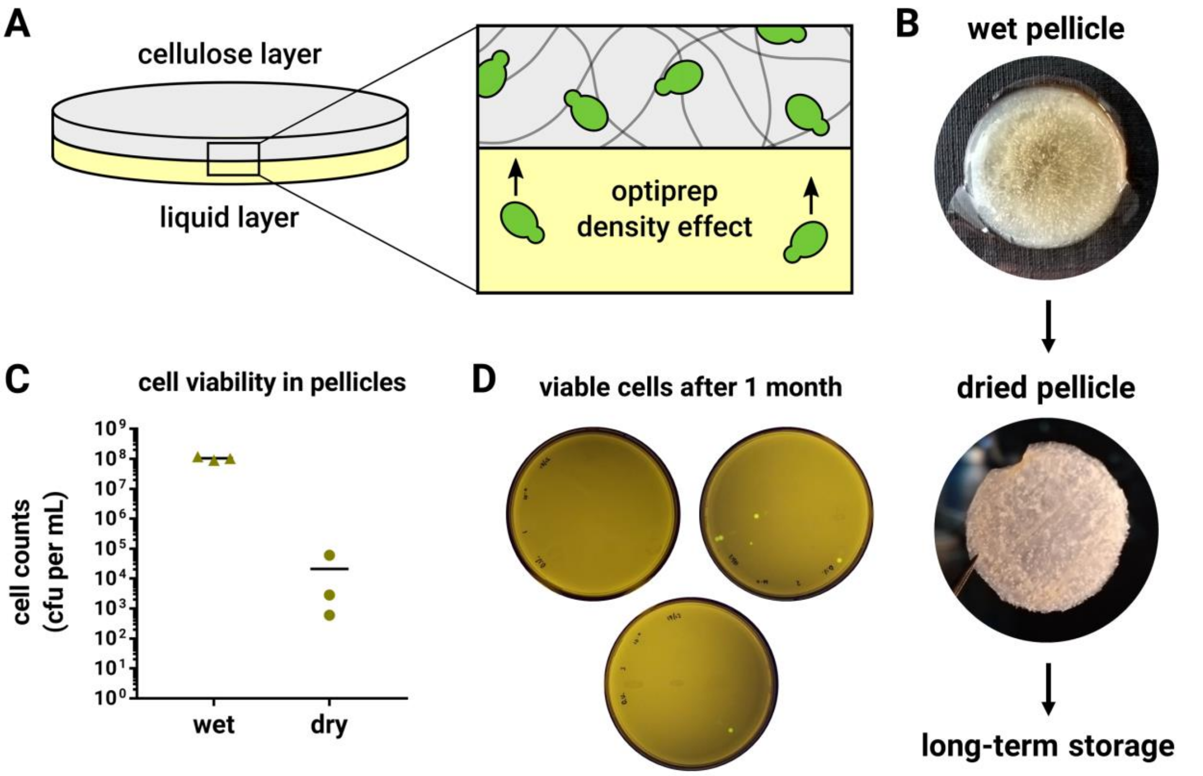


**Supplementary Figure 25. Cell viability in dried pellicles. (a)** Schematic illustrating the effect of Optiprep in the culture medium. By increasing culture medium density, *S. cerevisiae* cells become buoyant, rise to the surface and become incorporated into the BC matrix. **(b)** Pellicles into which *S. cerevisiae* cells have been incorporated can be dried and stored. **(c)** Cell viability was compared between wet and dried pellicles by enzymatically-degraded pellicles, plating and obtain counts of fluorescent *S. cerevisiae* cells. Bars represent the mean and green dots represent individual values. **(d)** After storage for 1 month, dried pellicles were enzymatically-degraded and 100 µL samples plated without dilution. The resultant plates are shown here, imaged for GFP fluorescence under translumination. Viable cells were obtained on two of three plates, indicating that a small number of *S. cerevisiae* cells survive even after 1 month of storage at room temperature.

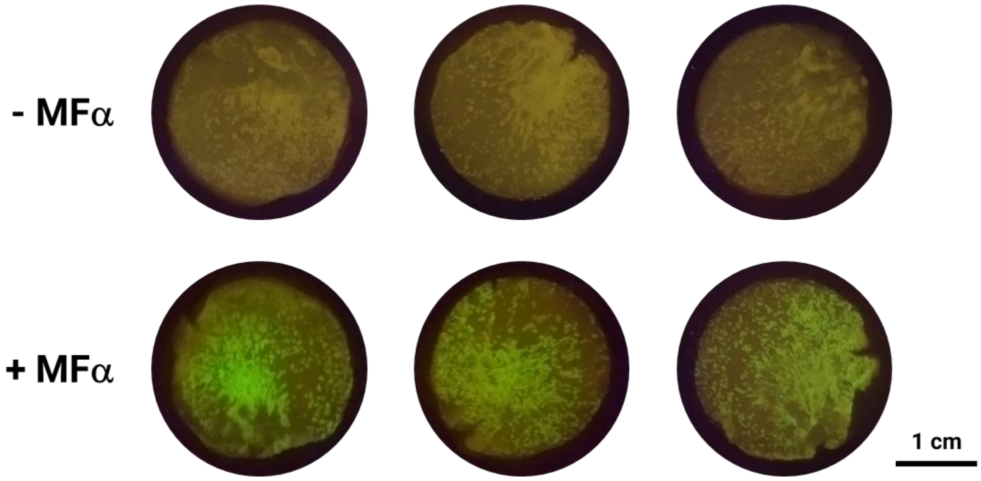


**Supplementary Figure 26. Sense-and-response BC materials using a GPCR-based biosensor strain.** Dried pellicles, into which the GPCR-based, MFα-responsive *S. cerevisiae* strain (yWS890) was incorporated, were incubated in fresh YPD medium without agitation in the presence or absence of MFα. After 24 hours, biosensor pellicles were imaged for GFP fluorescence under a transluminator. Samples prepared in triplicate.

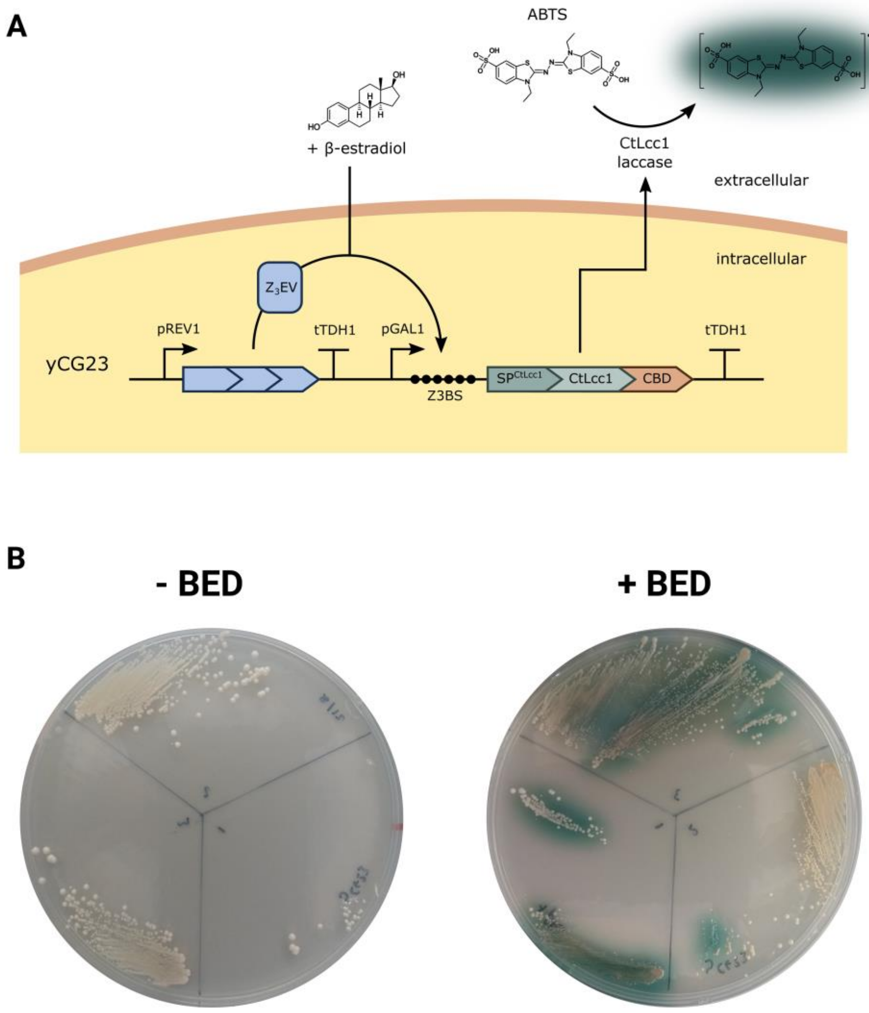


**Supplementary Figure 27. BED-inducible CtLcc1 laccase secretion. (a)** A strain, yCG23, was constructed for engineered CtLcc1 secretion in response to the presence of β-estradiol (BED). A two-gene construct was assembled using the YTK cloning system. The first gene encoded constitutive expression of the BED-responsive synthetic transcription factor $Z_3EV$. The second gene encoded a fusion of the CtLcc1 signal peptide and catalytic region fused to a C-terminal CBDcex domain, all under control of the $Z_3EV$-responsive promoter. In presence of BED, $Z_3EV$ will translocate to the nucleus, bind the Z3 binding sites (Z3BS) upstream of the CtLcc1-CBD ORF and induce transcription. CtLcc1-CBD secretion directed by the MFα signal peptide enables extracellular oxidation of the ABTS substrate to a colourimetric product. **(b)** Transformants were re-streaked in triplicate onto agar containing the colourimetric reporter of laccase activity, ABTS and $CuSO_4$ in the presence or absence of BED. After 3 days growth, a clear halo of green pigment was observed only in presence of BED indicating successful induction of CtLcc1 secretion.

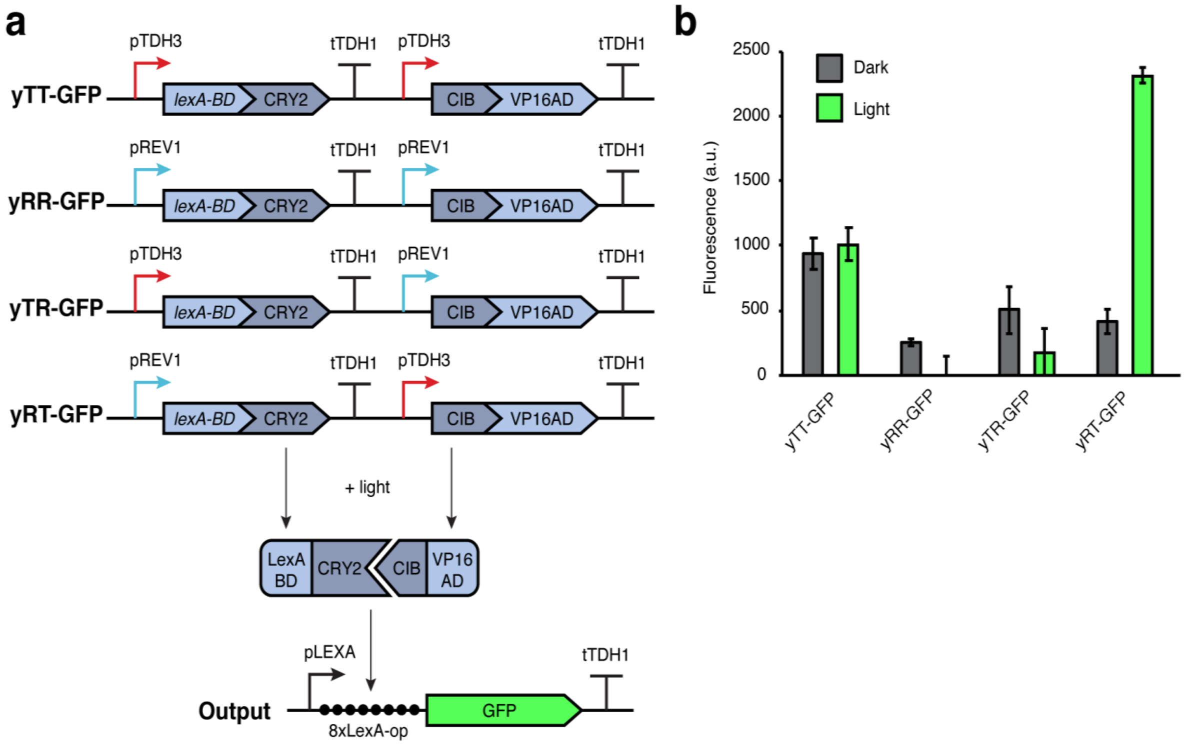


**Supplementary Figure 28. Optimization of light induction. (a)** Schematic illustrating the design of four combinations of promoter strength pairs. The strong constitutive promoter (pTDH3) is marked in red and the weak constitutive promoter (pREV1) is marked in blue. They drive the expression of the DNA-binding component and activation component, which together drive the expression of GFP in the presence of light. **(b)** GFP expression of yeast culture in liquid in dark or after 4 hours of light induction. Data represent the mean ±1 SD from biological triplicates.

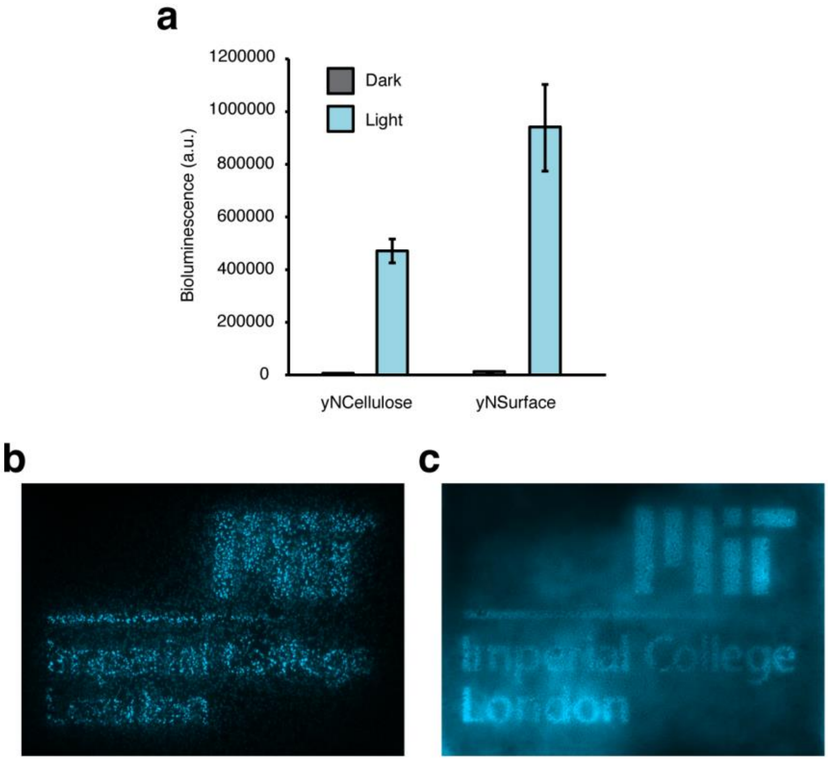


**Supplementary Figure 29. Light induction of luciferase expression and the tuneable resolution on BC living films. (a)** Bioluminescence of yNCellulose and yNSurface liquid culture after 4 hours of light induction. Data represent the mean ±1 SD from biological triplicates. **(b)** yNSurface ΔSED1 pellicle after 12 hours of exposure to projected pattern. This knockout strain showed slower growth compared to yNSurface. Less foci were formed and they are unable to provide enough resolution to reflect the pattern. **(c)** yNCellulose pellicle after 1-day outgrowth followed by 12 hours exposure. Increased yeast cell number on the surface provides higher resolution for patterning. Higher background activity reflects the diffusion of NanoLuc-CBD within the BC material.

### Supplementary References


(1) Chen, C., and Liu, B. Y. (2000) Changes in major components of tea fungus metabolites during prolonged fermentation. *J. Appl. Microbiol. 89*, 834–839.

(2) Bamba, T., Inokuma, K., Hasunuma, T., and Kondo, A. (2018) Enhanced cell-surface display of a heterologous protein using SED1 anchoring system in SED1-disrupted Saccharomyces cerevisiae strain. *J. Biosci. Bioeng. 125*, 306–310.

(3) Lee, M. E., DeLoache, W. C., Cervantes, B., and Dueber, J. E. (2015) A Highly-characterized Yeast Toolkit for Modular, Multi-part Assembly. *ACS Synth. Biol. 4*, 975–986.

(4) McIsaac, R. S., Gibney, P. A., Chandran, S. S., Benjamin, K. R., and Botstein, D. (2014) Synthetic biology tools for programming gene expression without nutritional perturbations in Saccharomyces cerevisiae. *Nucleic Acids Res. 42*, 1–8.

(5) Florea, M., Hagemann, H., Santosa, G., Abbott, J., Micklem, C. N., Spencer-Milnes, X., de Arroyo Garcia, L., Paschou, D., Lazenbatt, C., Kong, D., Chughtai, H., Jensen, K., Freemont, P. S., Kitney, R., Reeve, B., and Ellis, T. (2016) Engineering control of bacterial cellulose production using a genetic toolkit and a new cellulose-producing strain. *Proc. Natl. Acad. Sci. 113*, E3431–E3440.

(6) Pothoulakis, G., and Ellis, T. (2018) Synthetic gene regulation for independent external induction of the Saccharomyces cerevisiae pseudohyphal growth phenotype. *Commun. Biol. 1*, 7.

(7) Shaw, W. M., Yamauchi, H., Mead, J., Wigglesworth, M., Ladds, G., and Correspondence, T. E. (2019) Engineering a Model Cell for Rational Tuning of GPCR Signaling. *Cell 177*, 782-796.e27.

(8) Lee, C.-R., Sung, B. H., Lim, K.-M., Kim, M.-J., Sohn, M. J., Bae, J.-H., and Sohn, J.-H. (2017) Co-fermentation using Recombinant Saccharomyces cerevisiae Yeast Strains Hyper-secreting Different Cellulases for the Production of Cellulosic Bioethanol. *Sci. Rep. 7*, 4428.

(9) Villares, A., Moreau, C., Bennati-Granier, C., Garajova, S., Foucat, L., Falourd, X., Saake, B., Berrin, J.-G., and Cathala, B. (2017) Lytic polysaccharide monooxygenases disrupt the cellulose fibers structure. *Sci. Rep. 7*, 40262.

(10) Shinohara, N., Sunagawa, N., Tamura, S., Yokoyama, R., Ueda, M., Igarashi, K., and

Nishitani, K. (2017) The plant cell-wall enzyme AtXTH3 catalyses covalent cross-linking between cellulose and cello-oligosaccharide. *Sci. Rep. 7*, 46099.

(11) Pathak, G. P., Strickland, D., Vrana, J. D., and Tucker, C. L. (2014) Benchmarking of Optical Dimerizer Systems. *ACS Synth. Biol. 3*, 832–838.
